# Molecular Process Diagram: a precise, scalable and compact visualization of rule-based models

**DOI:** 10.1101/503359

**Authors:** Dan Vasilescu, James Greene, James C. Schaff, Ion I Moraru, Michael L. Blinov

## Abstract

Rule-based modeling allows representation and simulation of biological systems where molecular features (such as protein domains) and feature-specific details of molecular interactions are accounted for. The rule-based description is precise and can define very fine molecular details (e.g. how phosphorylation status of a single residue in a multi-protein complex can affect affinity of another binding site of another protein within the same complex), but makes it difficult to effectively combine the assumptions scribed within the multiple rules of a model into a diagrammatic view. Various visualization schemas have been suggested, but they are all highly rule-based centric (e.g. a visual list of unconnected rules, extended contact maps, or atom-rule graphs). None of them match the clarity of traditional reaction/pathway diagrams, where a researcher can easily visually track the transitions and modifications of chemical species occurring in the biological systems being modeled. Here we *present a novel approach and software for precise, scalable and compact representation of rule-based models* that we call Molecular Process Diagram. It is based on the three basic elements: interacting molecular complexes, molecular sites directly modified by a rule, and molecular sites that are not modified but contribute to a rule mechanism (e.g. a site that in a phosphorylated state changes binding affinity of another site). Multiple levels of resolution are available: pathway-like diagram of interactions among molecules, optional site-specific interactions, and additional contingencies for interactions. Inclusion of molecular sites enables unambiguous reconstruction of the rule descriptions from the visual diagram without additional supporting documentation, while still keeping a pathway-like visual appearance. *The proposed approach for visualization has been implemented in the Virtual Cell (VCell) modeling and simulation framework*. Our Molecular Process Diagrams extend the notion of Systems Biology Graphical Notation (SBGN) process diagrams and use SBGN-compliant conventions.

**Summary:** Kinetic models have provided significant insights into biological regulatory mechanisms even though they typically did not take into consideration the details of protein subcomponents such as binding domains and phosphorylation sites. However, these details are often required for an accurate understanding of the events that occur during cell signaling. Without such detailed understanding, intervention strategies to act on signaling pathways in pathological conditions are bound to have limited success. This need to include site-specific details into models led to the advance of rule-based modeling. While rules describe the details of interactions with unmatched precision, they often obscure the “big picture”, i.e. a pathway-like description of the information flow through the biological system. An intuitive visual diagram is crucial for understanding the assumptions embodied into a model. Here we present a novel approach and software for precise, scalable and compact representation of rule-based models that we call **Molecular Process Diagram**. It allows visualizing in a pathway-like diagram of the interacting molecules, the molecular sites modified, and the molecular sites that affect the interactions. The approach is implemented in the Virtual Cell (VCell) modeling and simulation framework and suggested as an extension for the Systems Biology Graphical Notations (SBGN) standard.

## Introduction

Kinetic models are important for predicting the dynamics and understanding the mechanisms of many biological processes. Biological regulatory mechanisms are largely governed by interactions among biomolecules. Most large biomolecules (proteins, RNA, DNA) contain multiple functional components, such as protein phosphorylation residues and SH2/PTB domains (1–4). Moreover, it has been known for a while that interactions among biomolecules generally depend on site-specific details. An early example was provided by elucidating receptor tyrosine kinase signaling, such as by the Epidermal Growth Factor (EGF) receptor: binding of adapter proteins to the receptor is mediated by the phosphorylation of receptor tyrosine residues and the affinity of the adapter proteins’ SH2 or PTB domains (5). Similarly, during IgE receptor (FceRI) signaling, which triggers allergic reactions, activation of kinase Syk is conditioned on phosphorylation by two other kinases of the tyrosine residues in the linker region and in the activation loop (6,7). Information about such site-specific context of interactions is increasingly becoming available for many systems, and so are models that take it into account. However, including site-specific details into conventional models (which use the explicit *reaction network approach*) leads to an excessively large number of molecular species and reactions to be considered, due to the combinatorial nature of protein modifications and protein complexes that can be generated (8,9). This exposes an inherent limitation of conventional models where the full list of the possible “reactions” (interaction kinetics) to be translated into the corresponding (stochastic or deterministic) mathematical equations must be specified. A rule-based approach (8,10–13) can overcome this limitation and enables a compact description, analysis and simulation of biomolecular interactions accounting for all site-specific details. Moreover, a rule-based model can serve as a compendium of information about interactions among multi-site molecules, with precise description of all modeling assumptions (14). However, a detailed model is of limited use unless it is presented in an understandable manner. The proteins and interactions included in a model, as well as the justification for modeling assumptions, should be communicated clearly and precisely if a model is to be understood, evaluated, and reused. Diagrams of reaction networks and pathways have long served as the de-facto standard for description of biological events. However, such diagrammatic presentations are not suitable for illustrating rule-based models.

Rule-based modeling capabilities were recently added to the well-established Virtual Cell (VCell) modeling and simulation framework (15). To enable a user-friendly specification and analysis of rule-based models, we implemented a graphical user interface that allows a visual specification of reaction rules (16). However, as we will discuss later, the implementation had the same shortcoming as previous rule-based visual representations: it was not possible to combine reaction rules into a diagram resembling a reaction network or a pathway. To overcome this limitation, we designed, developed and implemented a novel approach for precise, scalable and compact representation of rule-based models. In this manuscript we introduce the concept of a **Molecular Process Diagram (MPD)**, which combines features of reaction network visualization with details specific to rule-based modeling. Our approach allows visualizing site-specific details of molecular interactions within the intuitive paradigm of pathway visualization.

Next we provide a background on rule-based models and existing approaches to visualize them. We then give an overview of the basic principles of Molecular Process Diagrams and their use in the Virtual Cell software. We discuss how our visualization fits into the existing community standard for visualization, the Systems Biology Graphical Notation (SBGN, (17)) and propose SBGN-style notations consistent with Process Diagrams conventions.

### Pathways and networks visualizations

Traditionally, pathways were represented as graphs – most often bipartite graph where two types of nodes represent molecular entities and processes, respectively, and the edges connect molecular entities to the processes they participate in. A mathematical model of a pathway and the pathway itself are usually visually represented in the same style. Traditionally in the model the nodes are chemical species (equivalent to molecular entities) and reactions (equivalent to processes). Consider, for example, a pathway and a model of the initial events in signaling by Epidermal Growth Factor (EGF) receptor (EGFR) tyrosine kinase (**Fig. 1**). The first four steps in this pathway are: (1) EGF ligand binding to EGF receptor (EGFR); (2) dimerization of EGF-receptor complexes; (3) transphosphorylation of EGFR tyrosines by receptor tyrosine kinase; and (4) unprotected EGFR tyrosine dephosphorylation by phosphatases. These steps are usually represented by a pathway diagram that may merge some of these steps together, as illustrated in a pathway from PantherDB database (18) (**Fig. 1A**). Figure **1B** illustrates a simplified reaction network model built in the Virtual Cell (VCell) modeling and simulation framework corresponding to these three pathway steps.

**Figure 1.**
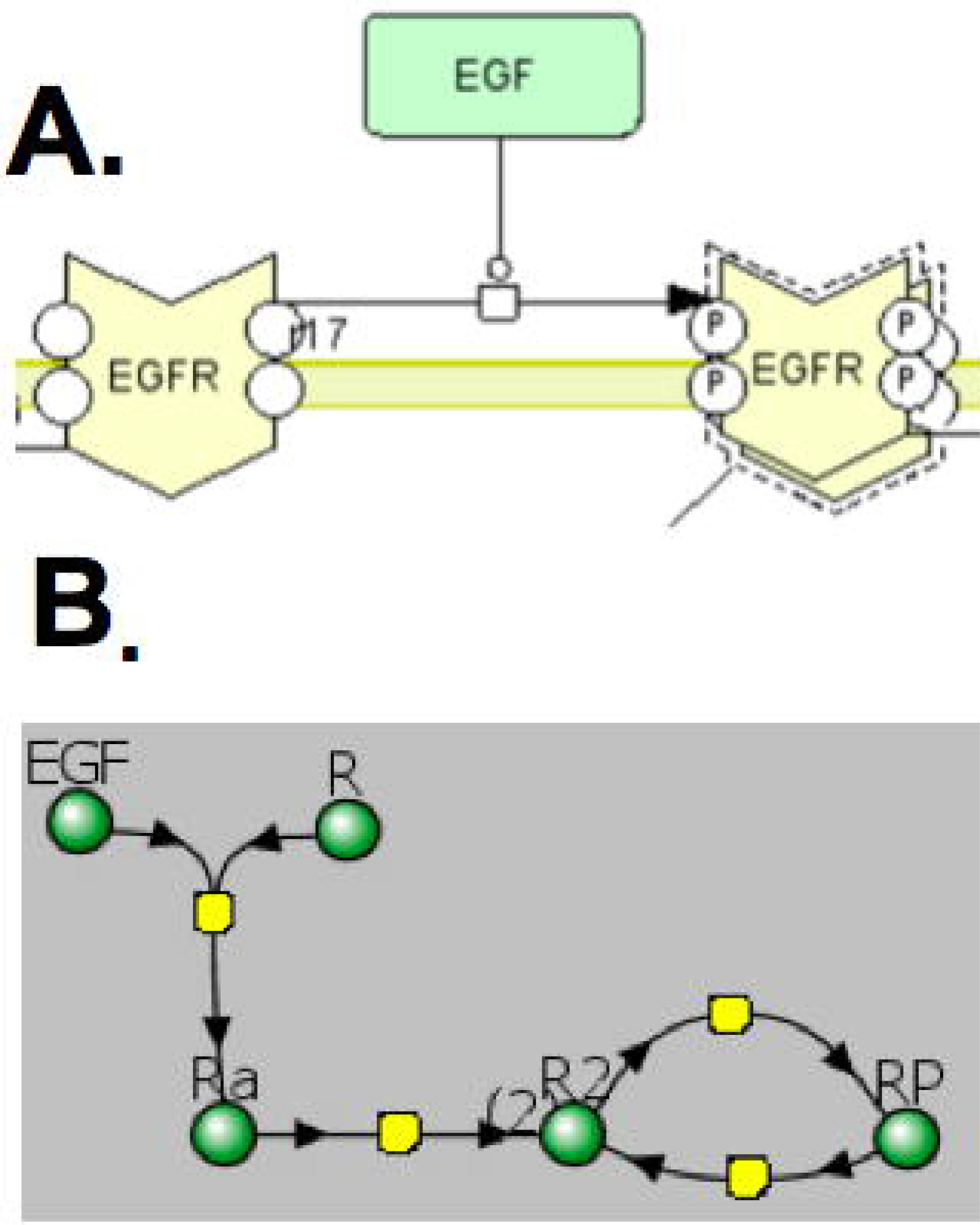
Visual representations of pathways and reaction networks models. **A.** A screenshot of PantherDB illustrating EGF receptor dimerization and autophosphorylation stimulated by EGF binding. **B.** The corresponding reactions as depicted in a VCell model: receptor R binds ligand EGF forming species Ra (a complex of EGF-EGFR), which undergoes dimerization to form species R2 (EGF-EGFR dimer), which undergoes phosphorylation to form species RP (fully phosphorylated dimer).

### Complexity of biological systems

In the conventional modeling approach, a reaction network consists of chemical species and reactions among them. Each chemical species has to be created, named uniquely, and reactions specified and assigned to appropriate locations such as membranes or volumetric compartments. For each interaction described in the model, the user chooses the appropriate kinetic laws and relevant parameters. This approach is implemented in many popular software environments such as VCell (15), COPASI (19) or CellDesigner (20). However, the manual specification of each and every model component leads to obvious limitations on the number of species and reactions that can be modeled. Modelers often introduce bias by imposing ordering of processes and lumping of species in order to limit the number of species in the model. Without such simplifying assumptions the number of molecular species and reactions needed may be completely intractable: in our example of EGFR signaling, the EGF receptor has nine tyrosines subject to independent phosphorylation and binding events. Thus, 3^9^ = 19,683 different EGF receptor complexes can possibly exist (each site tyrosine be in three states – unphosphorylated and unbound, phosphorylated and unbound, and phosphorylated and bound to its binding partner) but modelers will typically consider a single species of fully phosphorylated receptor. This combinatorial complexity explodes once we consider binding partners that can undergo modifications and binding to other biomolecules. Nevertheless, the real scope of complexity in a biological system may lie well below such high numbers of possibilities because of spatial and structural constraints that may limit individual molecular interactions. However, it is still desirable to be able to explore the whole multitude of all possible complexes and phosphoforms in order to eliminate bias and capture critical features of variability in signaling (9). The reaction network approach is not capable of providing this ability, and as a result an alternative approach of rule-based model description was developed.

### Rule-based approach

The rule-based approach provides an opportunity to consider the whole nomenclature of potential molecular complexes and interactions among them that can potentially occur. A model is defined by reaction rules, which are precise statements about biomolecular interactions and modifications. Given a set of species, a reaction rule identifies those species that have the features required to undergo a particular transformation from reactants to products. Interactions represented in a reaction rule depend only on specifically indicated features. Thus, multiple individual reactions involving distinct species can be in fact defined by a single reaction rule (**Fig 2**). The same four pathway steps in the EGFR signaling can be described by the following four rules that repeat the pathway steps but provide a lot more information of what is important and what not for the interactions. Rule (1) states that the EGF ligand binds to an EGFR monomer conditional on the existence of an available extracellular site but independent of the state of any other biomolecule in the system (including the state of the intracellular portion of the receptor itself – e.g. whether some of the receptor tyrosine residues are phosphorylated or whether some binding sites are occupied by their binding partners). This reaction rule corresponds to various individual binding and unbinding reactions of EGF with receptor-monomers that have only one feature in common: EGFR is not currently bound to another ligand. Rule (2) states that a receptor dimerizes with another receptor conditional on ligands being bound to the extracellular site on both receptors but independent of the state of other biomolecules in the system or the states of the receptor intracellular sites. Rule (3) states that tyrosine residues of a receptor are trans-phosphorylated by receptor tyrosine kinase conditional on the receptor being part of a dimer complex (bound to another receptor at the trans-membrane portion) but irrespective of the state of receptors or other biomolecules. Rule (4) states that phosphorylated tyrosine residues of receptor can be dephosphorylated conditional on being unprotected (not bound to any other protein) but independent of the state of other sites. **Figure 3** illustrates all four rules using the conventions introduced in the VCell modeling and simulation framework (16).

**Figure 2.**
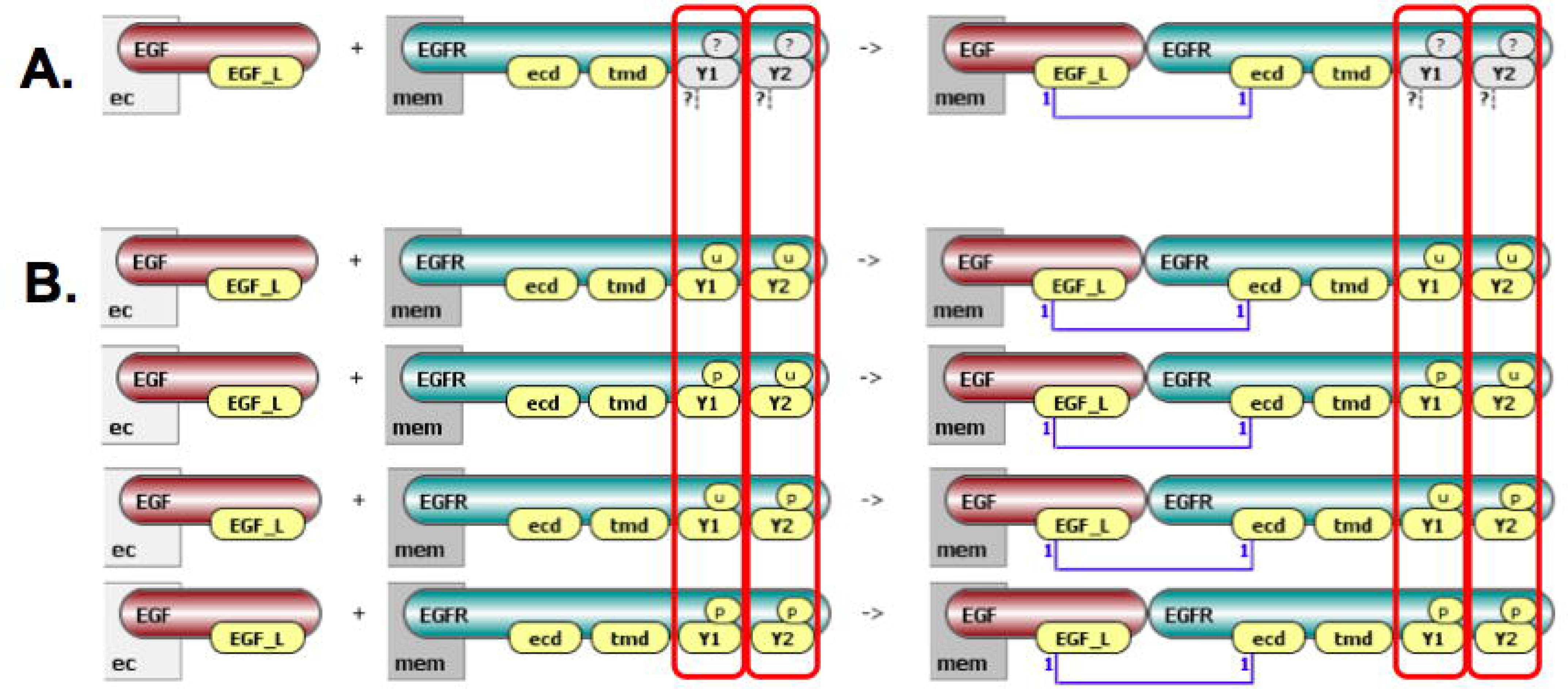
Ligand-receptor binding visualized in VCell. **A**. The rule for receptor ligand binding. The rule has two reactant patterns (ligand EGF and receptor EGFR), and one product pattern (a complex of EGF and EGFR). EGFR has four sites: “ecd” stands for extracellular domain, which binds to EGF; “tmd” stands for trans-membrane domain, which can bind to another EGFR molecule; “Y1” and “Y2” are tyrosines which can be phosphorylated and can bind other molecules. Yellow shapes denote sites that must be unbound while white shapes denote sites that do not affect the reaction rule (both tyrosines can be in any states – question mark above site – and can be bound or unbound – question mark underneath site). **B**. Four possible individual reactions that correspond to this rule, with tyrosines being in all combinations of phosphorylated and unphosphorylated states. “Mem” underneath molecule box stands for a membrane location, while “ec” means extracellular location.

**Figure 3.**
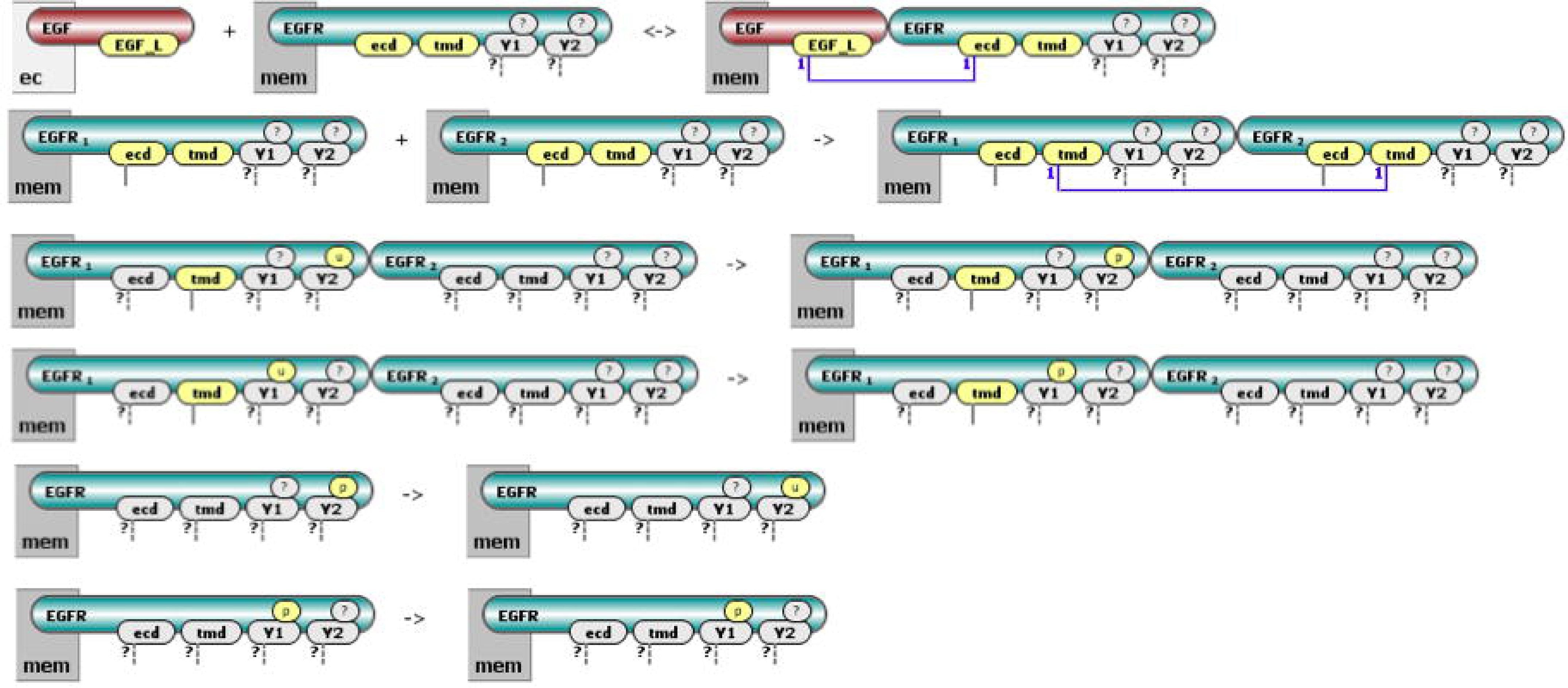
VCell cartoons describing the initial steps in EGFR signaling with six rules: a rule for ligand-receptor binding, one for EGFR dimerization at “tmd” site (“ecd” site shown as having to be bound to ligand on both receptors), two rules for tyrosine phosphorylation (receptor must be in dimer form), and two rules for tyrosine dephosphorylated (receptor must be monomer).

A simple way to use the rule-based approach is to translate the reaction rules into machine-generated conventional reaction networks which are then being simulated, as implemented in the BioNetGen software (10,21,22). This relieves the effort to manually generate the large number of all individual species that can potentially exist during the simulation time and the simulation results can be post processed to produce timecourses of observables of interest (typically functions that group molecular species that share certain features of interest – e.g. the total number of molecules in a specific phosphoform). However, this approach will be of little value if the number of possible complexes and reactions is so large that the generated network cannot be effectively simulated. On the other hand, in these extreme cases the biological reality is that not all of the possible molecular states are actually being formed, as this number is obviously bound by the total number of molecules present in the system. Since the total number of proteins and other biomolecules in typical models of intracellular system is below what can be handled by stochastic simulations run on modern computers, a different approach for using the rule-based description was later developed: network-free simulations (11), which are agent-based simulations where the agents are all individual molecules of the system. All possible individual species do not need to be generated, and only observables are being tracked over the timecourse of the simulation. Several other software tools now also implement rule-based modeling capabilities including network free simulations: KaSim (23,24), Simmune (25,26), rxncon (27) and VCell (16). The rule-based description is also used as a compendium of information about molecular interactions on a site-specific level (14,28).

### Visualization of rule-based models

One of the drawbacks of the rule-based modeling paradigm is that the representations of the new modeling concepts being introduced (i.e. molecules, sites, bonds, states, patterns and rules) must be learned, and they are being presented differently by the various modeling frameworks. A rule-based model is usually coded in a specially designed language, such as BioNetGen Language (BNGL,(21)), Kappa (23,24), PySB (29) or rxncon (27). This approach requires learning of a formal, domain-specific language, similar in usage to writing a computer program, and introduces a steep learning curve for a modeler inexperienced in the language of choice. Therefore, several graphical representations have been proposed to ease the design of rule-based models.

Expanding on the example of EGFR signaling will illustrate the complications that arise when trying to visualize rule-based models. The initial signaling events described are followed by additional steps of recruitment of Grb2 and Shc adapter proteins to the phosphorylated receptors: phosphorylation of Shc, recruitment of Grb2 to phosphorylated Shc and sequential recruitment of Sos guanine nucleotide exchange factor to the membrane via its interaction with Grb2 (30). The full model accounts for 3749 interactions among 356 species, and describes a large number of interactions that a molecular complex can undergo: there may be up to ten adapter proteins bound to two distinct tyrosines per receptor in a dimer, dimers can break up into monomers, lose ligands, but still function as protein complex assembly, etc. This precludes the standard approach of visualizing each and every reaction, and cartoons for visualization of individual rules were initially suggested (25,31–33). In **Figure 2** we used conventions for direct graphical specification of individual rules: every molecular entity (protein, biomolecule, DNA) is represented as a “molecule class” – a container with components (“sites”) that define the particular features of this entity; any species can be represented as a graph consisting of one or more containers connected together via site-site bonds. Each reaction rule is thus specified as a separate cartoon describing reactants and products. This approach was implemented in Simmune (25,26,34), and more recently in VCell (16) (**Fig 3**). Such visualization of individual rules is perfectly appropriate to capture the details of every interaction in easily understandable pictograms that do not require one to be familiar with arcane symbolic languages. Unfortunately, individually illustrating every rule in models composed of large number of rules results in a long sequence of diagrams that are locally comprehensible but globally incomprehensible (see examples in **Supplemental Material**), and runs counterintuitive for most modelers who are used to think in terms of networks and pathways.

Several approaches have been suggested to assemble graphical rules into a connected “pathway-like” diagram. A reaction network looks similar to a pathway description, because a product of one reaction is a chemical species that can participate in another reaction. Similar to conventional reactions in a network, rules have reactant patterns and product patterns referred by the rules they participate in. However, rules operate on sets of species, and they can draw their reactant species from products species of multiple other rules. Thus it is rare that the output product of one rule can be used directly as the input to another rule (35). A direct graph translation where each node corresponding to a distinct reactant or product pattern in the set of reaction rules would be disconnected (**Fig. 4**).

**Figure 4.**
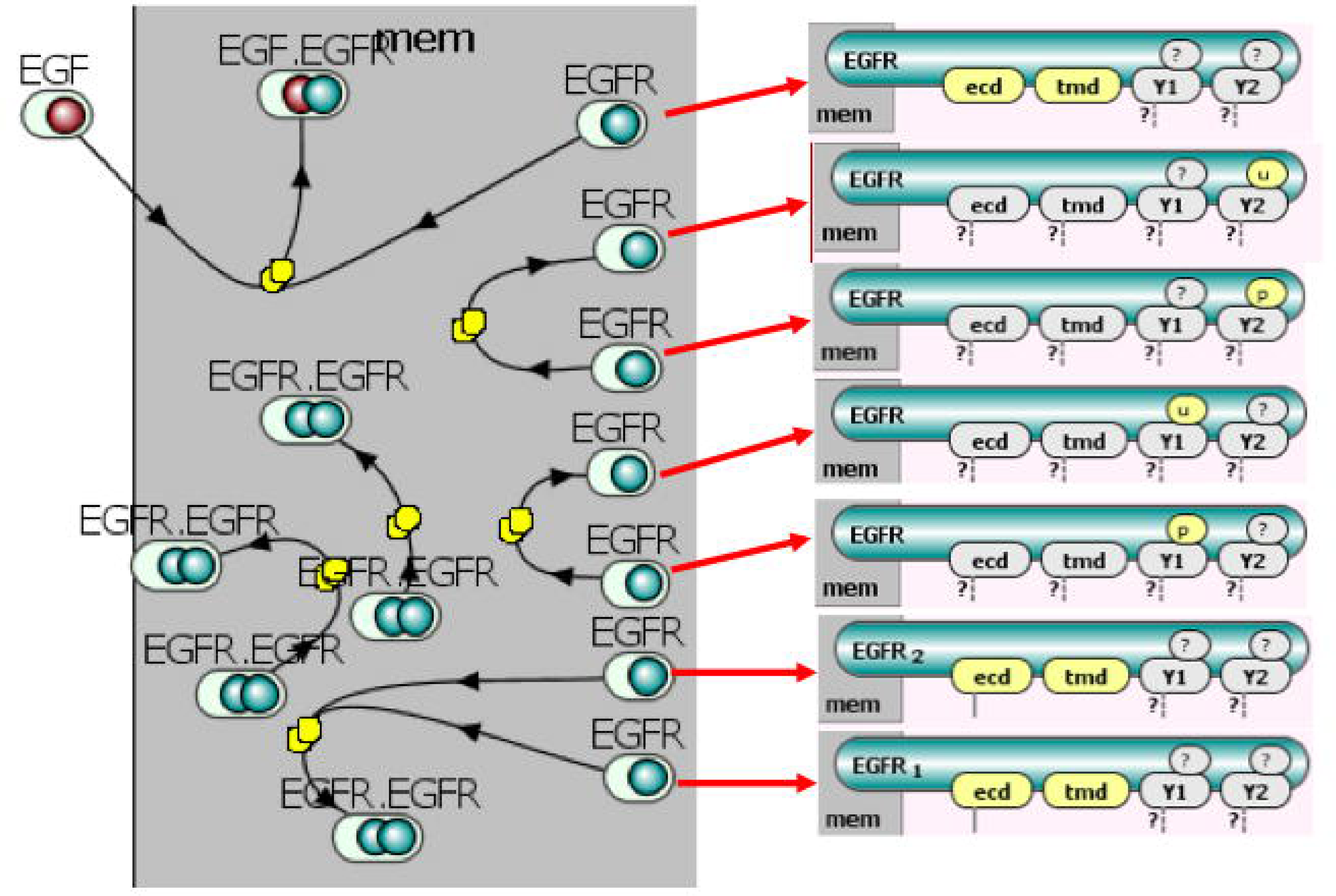
Direct graph representation of rules. The resulting bipartite graph shown at left is comprised of nodes for reaction rules and nodes for distinct reactant and product patterns is usually highly disconnected, because many (in this case all) of the reactant and product patterns from different rules are slightly different. This is illustrated on the right by showing the detailed state of monomeric EGFR in seven different patterns.

One approach is based on displaying molecules and site-site interactions among these molecules. A contact map (24) representing all possible interactions among sites of molecules was followed by an extension to molecular interaction maps (MIM, (36,37)), and extended contact maps (ECM, (38)). These representations are really compact, they showcase molecules, their internal states, and the possible interactions among molecules represented by rules, and they do provide a connected view of model interactions. However, there are several disadvantages. For one, they fail to pin-point the temporal order of interactions. Second, and more importantly, the conditions under which each interaction may happen is not easy to understand and track back to the reaction rules (36); as a result the set of rules used to create the model generally cannot be uniquely recovered without additional information given. Overall, the major weakness of these visualization approaches is that the information flow through the system is obscured: contact or molecular interaction maps are drastically different from the commonly accepted representation of signaling by a graph that can be traversed to explore individual signaling paths.

Most recently, three other visualization approaches were suggested to remedy this issue, attempting to link some elements of the rules into a connected network. First, the rule influence diagram (39) is a graph with nodes being rules and edges displaying rules influence on each other. Second, the atom-rule graph (AR graph,(40)) is a bipartite graph where one type of nodes (“atoms”) represents a class of molecular features (which can be a single site, a specific state of a single site, or a complex of multiple sites), and another type of nodes (“rules”) represents reaction rules. Three types of edges connect “atom” and “rule” nodes: a reactant or a product edge is drawn if an instance of the atom is modified on the left or right side of the rule, respectively; and a context edge is drawn if a rule is contingent on the atom. Both the rule influence diagrams and AR graphs are capable of representing all elements of a rule-based description, but they rely heavily on the user being familiar with rule-based descriptions, and they are still not intuitive to a biologist familiar with signaling pathways and reaction network representations. Third, the Simmune NetworkViewer (26,34) is a bipartite graph where one type of nodes represents reactant and product patterns that have the same molecules and bonds, but differ in internal states, while the second type of nodes represent the rules that patterns participate in. This approach is the most intuitive one and resembles a pathway description, but it obscures causal dependencies on internal states and is prone to combinatorial explosion in the number of nodes when molecules have multiple ways of being connected within a molecular complex.

Ideally, a visualization for rule-based models should be: (1) Intuitive – resembling a traditional reaction network or pathway diagram; (2) Precise – unique for a given set of rules; (3) Unequivocal – the original set of rules can be inferred back from the pictogram; (4) Scalable – have different levels of resolution from a model overview to full details; and (5) Compact – do not exhibit combinatorial complexity but grow linearly with respect to the number of rules. We believe that the novel approach to visualize rule-based models presented below satisfies these requirements.

## Methods

### Molecular Process Diagram

The overall goal of our visualization approach is to represent a rule-based model as a connected diagram that carries all the information about the reaction rules but also provides a view of information flow through the system. We define a “molecular complex” as all the molecules that are specified by a rule pattern. For example in the case of EGFR signaling the receptor dephosphorylation rule will have as both reactant and product molecular complexes a single EGFR molecule, whereas the dimerization rule will have as a product a molecular complex consisting of two EGFR molecules. We then construct a **Molecular Process Diagram** (MPD) as a bipartite graph with two types of nodes: “molecular complex” and “process”. Thus, the MPD representations of a rule-based model will be a graph with a number of process nodes equal to the number of rules and a number of molecular complex nodes no larger than the sum of the number of reactant patterns and the number of product patterns present in the rule set. The MPD representation of the system of 6 rules represented as 6 cartoons in Figure 2 is a bipartite graph with 6 process nodes and 4 molecular complexes (**Fig. 5A**).

This representation looks quite similar to a pathway: one can easily see that EGF binds to EGFR to form the EGF-EGFR molecular complex, the EGFR receptor dimerizes, the EGFR molecules in the dimer can undergo two distinct types of modifications, and the monomeric EGFR receptor can also undergo two modifications. While this representation is not showing all the site-specific information (we cannot immediately infer what those modifications are – which sites are being modified and how), it is very intuitive and captures a logical overview of the signaling pathway.

### Adding site-specific details to Molecular Process Diagrams

To view the site-specific information one just has to click on a process node in the MPD to expand it into a cartoon presenting the details of the corresponding reaction rule (**Fig. 5B**); similarly, every molecular complex node can be clicked and expanded to show the set of exact reactant and product patterns. The VCell-implemented Molecular Process Diagram thus provides a concise and intuitive representation of the rule-based model, while individual cartoons provide the detailed descriptions of all model assumptions. An additional problem is the fact that it is not trivial to create effective cartoons; just as for the textual BNGL representation of rules, often the sheer number of details can make it difficult to identify the most salient molecular features of the reaction rule. To address this problem, we implemented optional highlighting of different features: one can turn on or off highlighting of *the interacting molecules* (which can be displayed using distinct colors like in **Figure 5** or all gray like in **Figure 6**), of *the sites being modified* by the rule (**Fig. 6B**), and of *the molecular context* – i.e. sites that do not change their state, but the process requires them to be in a particular state (**Fig. 6C**).

**Figure 5.**
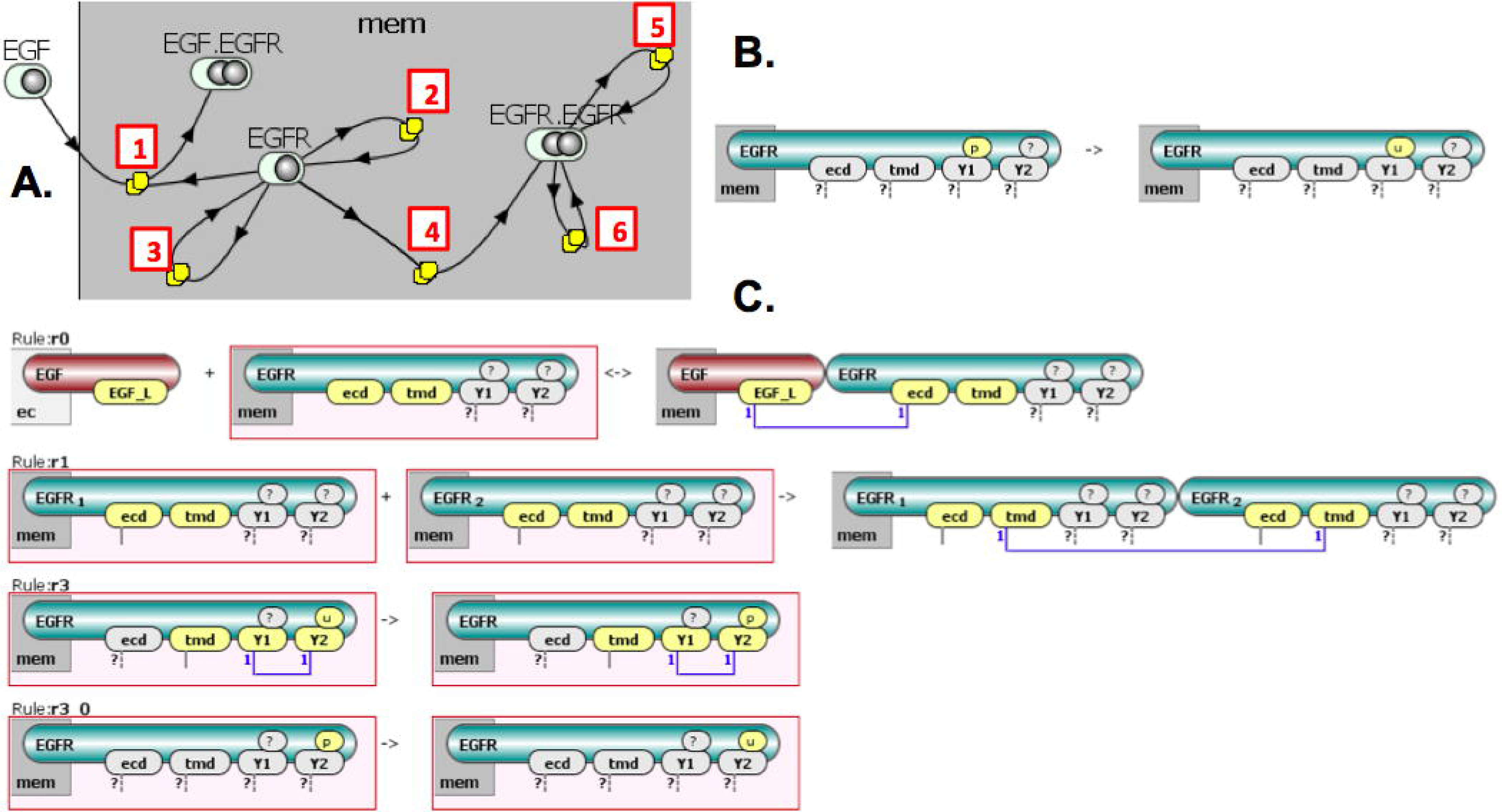
VCell layered graphical representation of a rule-based model. **A**. The pathway-like Molecular Process Diagram shows that EGF binds to EGFR [1] to form an EGF-EGFR molecular complex; EGFR can undergo two modifications [2,3] and can dimerize [4]; the EGFR-EGFR dimer also can undergo two other modifications [S,6]. **B**. Bly clicking on a reaction rule node in the diagram, one can see site-specific details of the rule (shown here is unimolecular interaction [3] by dephosphorylation of site “Y1”). **C**. By clicking on a molecular complex node, one can see the different state patterns of the molecular complex from all the different rules it participates in, with the particular complex highlighted by a red box (shown here is EGFR which participates in four rules).

**Figure 6.**
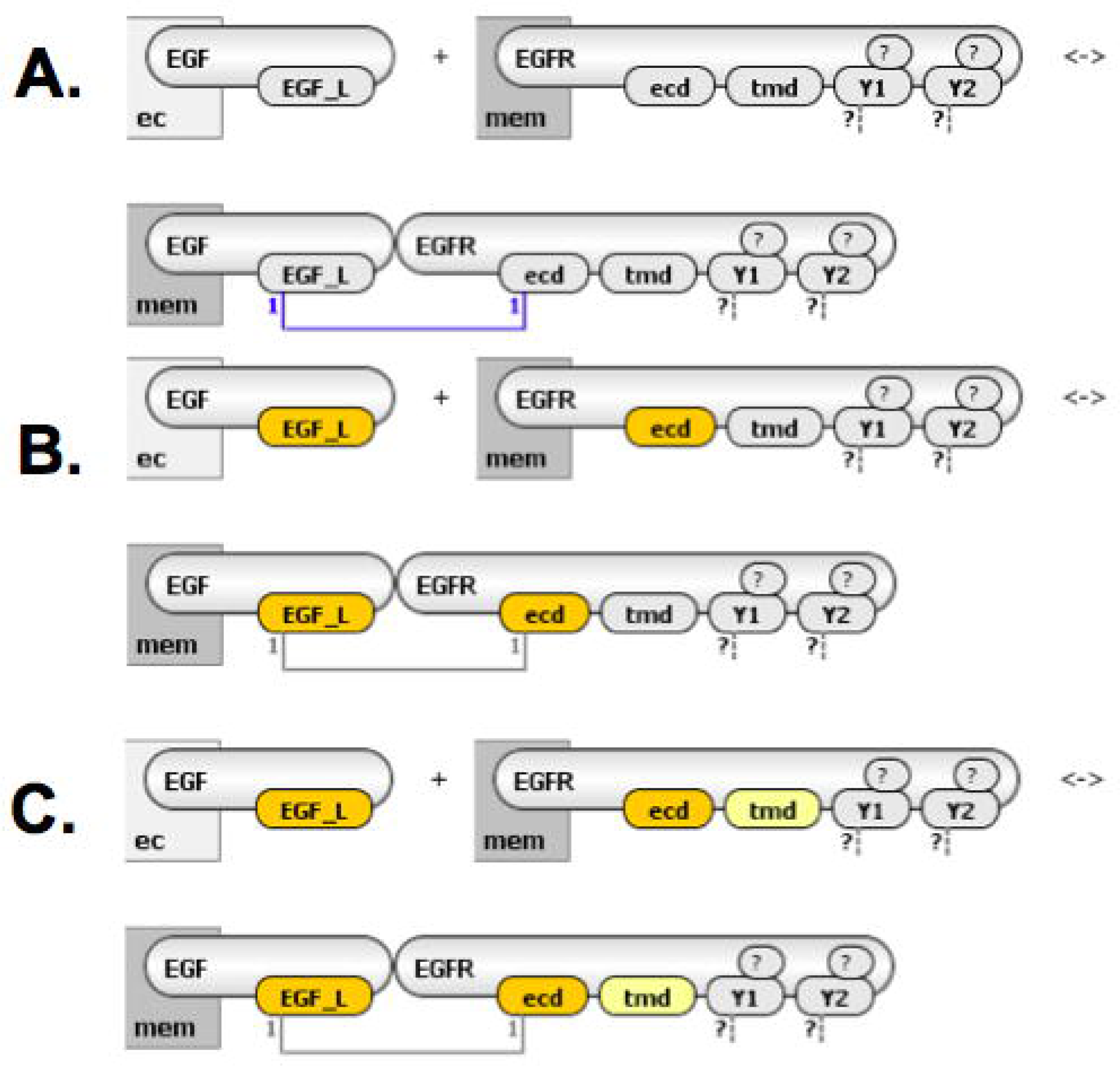
Highlighting specific details of reaction rules in VCell. **A**. Interacting molecules with all sites are shown but no highlighting. **B**. Sites being modified as a result of the rule are highlighted in bright yellow, while all sites that remain unchanged are not colored. One can immediately notice what is happening (“r” site of ligand and “ecd” site of receptor change from “unbound” to “bound”). **C**. Additional highlighting in pale yellow of sites that remain unchanged but need to be in a particular state for the rule to take effect (molecular context). One can easily identify the molecular features required for the interaction to happen (“tmd” site of receptor must be unbound).

### SBGN-compliant Molecular Process Diagrams

The Systems Biology Graphical Notation (SBGN (17)) is a community effort to develop standard graphical languages for representing biological processes and interactions. The challenge is that the proposed notations should be human readable (41) and visually appealing but at the same time machine readable (which imposes certain limitations, e.g. glyphs have to be color- and width-agnostic, the notion “near” cannot be used, etc.). There are three orthogonal visual languages defined in SBGN: Process Description (PD), Entity Relationship (ER), and Activity Flow (AF). A PD diagram depicts how physical entities transition from one form to another as a result of different influences; it represents all the molecular processes and interactions taking place between biochemical entities, and their results, and it demonstrates the temporal order of molecular events occurring in biochemical reactions. The PD diagrams are most relevant to the visualization of rule-based models proposed here. They are conceptually similar to the “classical drawings” of signaling or metabolic pathways that one would find in textbook cartoons, and the SBGN PD language is used by various pathway databases (e.g. PantherDB (18), Reactome (42), GeneXplain (43)) and supported by a number of modeling software tools (e.g. CellDesigner (20), COPASI (19), JWS online (44), BioUML (45), BIOCHAM (46)). Many pathway visualization tools that support Biological PAthway eXchange (BioPAX) standard (BioPAX (47)) provide visualization in the form of SBGN-PD (e.g. VISIBIOweb (48), CHiBE (49), Cytoscape plugins (50), SBGNViz (51)). A support for visualization tools that implement SBGN is provided by dedicated software library libSBGN implementing a community-developed markup language SBGN-ML (52).

An SBGN PD diagram is a bipartite graph with two classes of nodes: entity pools and processes. Nodes may reside in different compartments and each class of nodes uses several different glyph types that convey additional information about the corresponding molecular entity (simple chemical, macromolecule, complex, etc.) or process (transition, association, dissociation, etc.). The edges of the graph are one of several types of connecting arcs (production, modulation, stimulation, etc.). The biological meaning of the molecular interactions depicted in a PD diagram is thus gleaned from these visual representations of the specific types of entity pool nodes, process nodes, and connecting arcs. For the purpose of representing rule-based constructs, we limit ourselves to just one type of process node (transition) and two types of connecting arcs (consumption and production). However, the use of other glyphs may enrich the diagram and still keep it in correspondence with a rule-based model. On the other hand, with regard to entity pool nodes, different types will be used (such as simple chemical, macromolecule, etc.) and several entity pool nodes may be assembled into container nodes representing complexes.

Our suggested Molecular Process Diagrams for rule-based models is using the SBGN PD diagram approach, utilizing existing glyphs and syntax of SBGN-PD, but introducing new semantics. Similar to SBGN PD diagrams, a Molecular Process Diagram includes information about location (compartments where entities reside), about the type of physical entities (proteins, small molecules, dimers, etc.), and about the types of interactions among molecules (modification, translation, transcription, etc.). We can create SBGN PD-compliant representations at different levels of resolutions, starting with the coarse-grained MPD and including additional details corresponding to the different highlighting options of an expanded process node (refer to **Figures 5** and **6** above).

## Results and Discussion

In **Figure 7** we present the most coarse-grained level of a Molecular Process Diagram drawn in yED (a software tool that can produce fully SBGN-compliant visualizations; YWorks), which corresponds to the VCell MPD diagram shown in **Figure 5A**. The SBGN PD multimer glyph is used for the receptor dimer and state variables are used to declare the binding and modification sites. To declutter the diagram molecular sites are only shown when an individual molecule is displayed and not when it is a participant in a multimer or complex.

**Figure 7.**
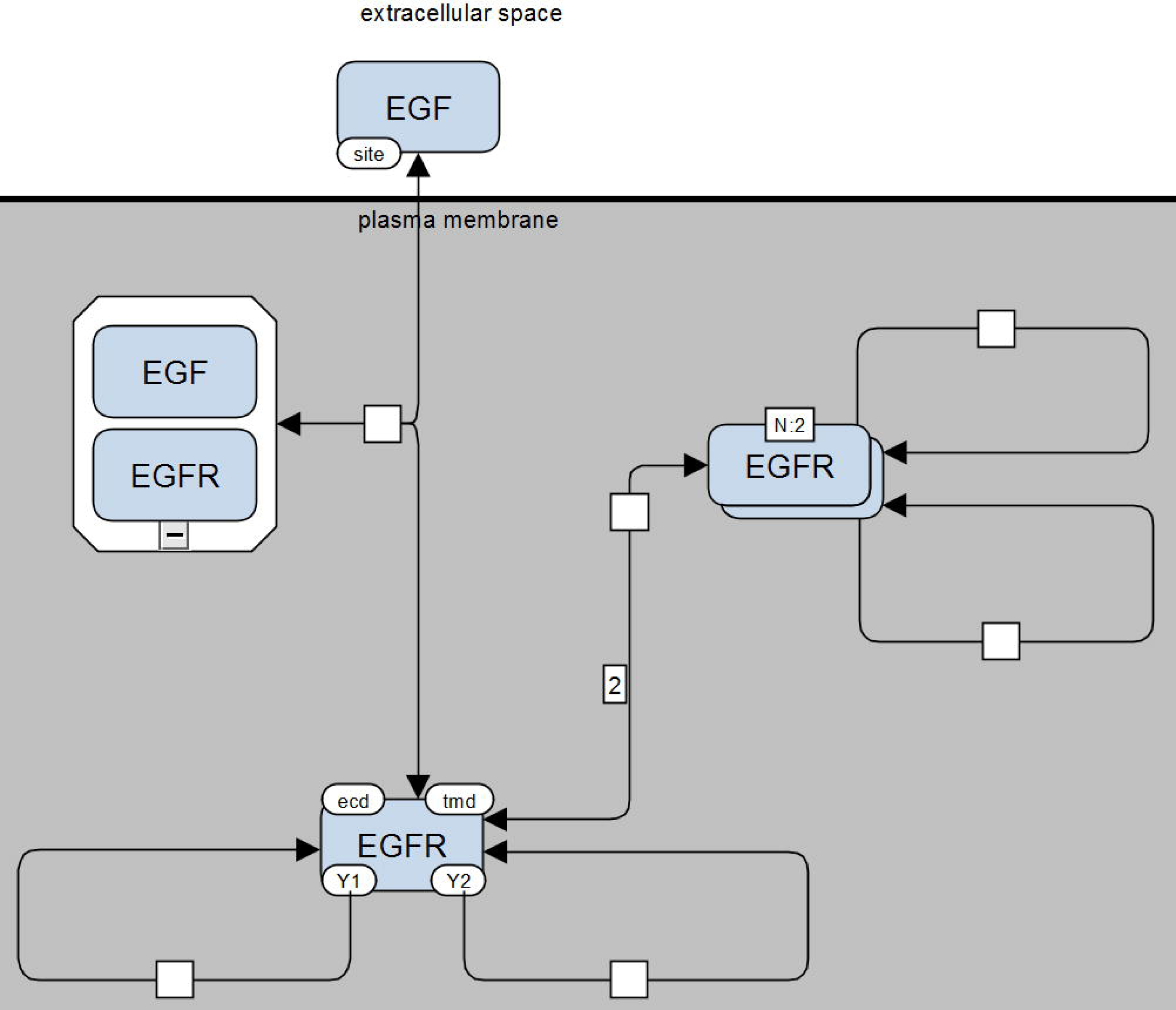
Molecular interactions of a Molecular Process Diagram drawn in yED. This diagram corresponds to the VCell version of the diagram shown in **Figure 5A**. It is fully compliant with SBGN-PD. The macromolecule glyph is used for EGF and EGFR, the container node glyph for the EGF-EGFR complex, and the multimer glyph for the EGFR dimer. Molecular sites (“site”, “ecd”, “tmd”, “Yl” and “Y2”) are drawn using state variable glyphs (note that they are shown only once in a diagram, where an individual molecule is being displayed and connectivity of complexes is not detailed). Each process node corresponds to a rule, and the rule stoichiometries that have values other than 1 are displayed on arcs (note “2” shown on consumption arc of dimerization rule).

In **Figure 8** we present a more detailed version that adds sites being modified by a rule. These are presented at both sides of the corresponding process node in SBGN-compliant information boxes. Sites defined in the reactant patterns are located on the consumption arcs while sites defined in the product patterns are located on the production arcs. Site state is being labeled consistent with SBGN conventions (e.g. we use P@Y1 to indicate site Y1 is in a phosphorylated state). The SBGN requirements for machine processing are being met by placing the information boxes for states such that they touch the process nodes. To declutter diagram, sites that participate in binding-unbinding reactions are shown only on the consumption arc: the convention is that these sites are unbound before the process takes place and become bound as a result.

**Figure 8.**
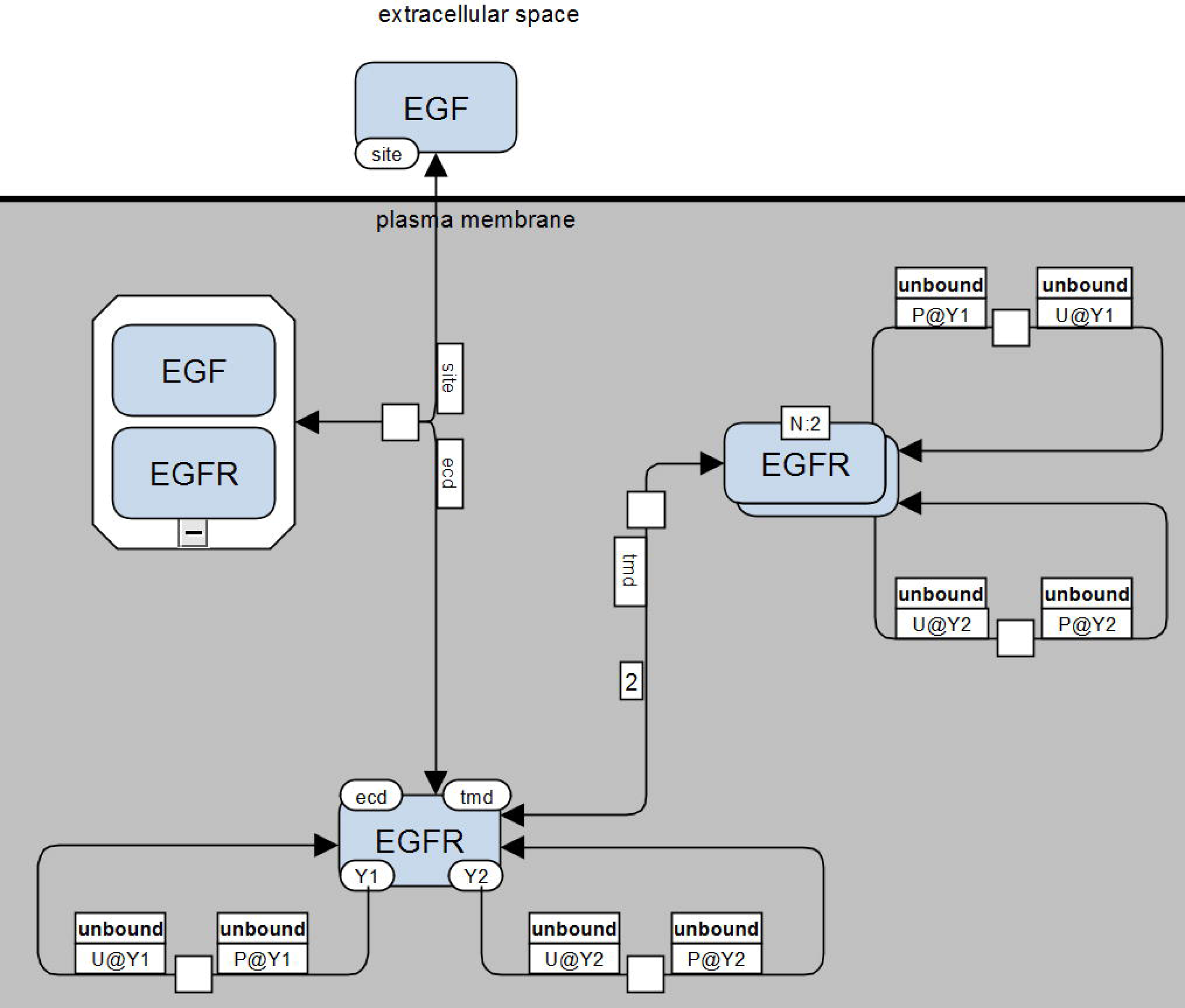
SBGN-style visualization of site-specific information of a Molecular Process Diagram. EGF site “site” binds to EGFR site “ecd”. Two EGFR molecules bind together at sites “tmd”. A single “Y1” site in EGFR dimer undergoes state change from “U” to “P”, and the same for “Y2” site. The EGFR monomer undergoes two processes, where each of unbound sites “Y1” and “Y2” can change state from “U” to “P”.

In **Figure 9** we present an SBGN PD-compliant drawing of the Molecular Process Diagram with all site-specific details included. Adding the molecular context (about sites that do not change their state during a process, but the process requires them to be in a particular state for the process to happen) in a concise but precise manner can be difficult because a rule may depend on the molecular features of reactant species in a multitude of different ways (e.g. states of sites, whether the sites in a complex are bound in a specific way, whether some sites are unbound or bound to some – possibly unspecified – molecules outside of a reactant complex, etc.). To allow for the diverse possibilities, we introduce an additional information box placed directly above the information boxes defining the sites. When such contingencies are specified, the additional box for the respective sites will be marked such as “unbound” whenever the site is unbound, “bound” when the site is bound, but without requiring a specific binding partner, or with a name (of the site it is bound to) when the site is bound and requiring a specific binding aprtner. A single box means that the site is in undefined binding status, which corresponds to a question mark in BNGL code. **Figure 10** provides examples of visualizing different combinations of molecular features used as molecular context in rules.

**Figure 9.**
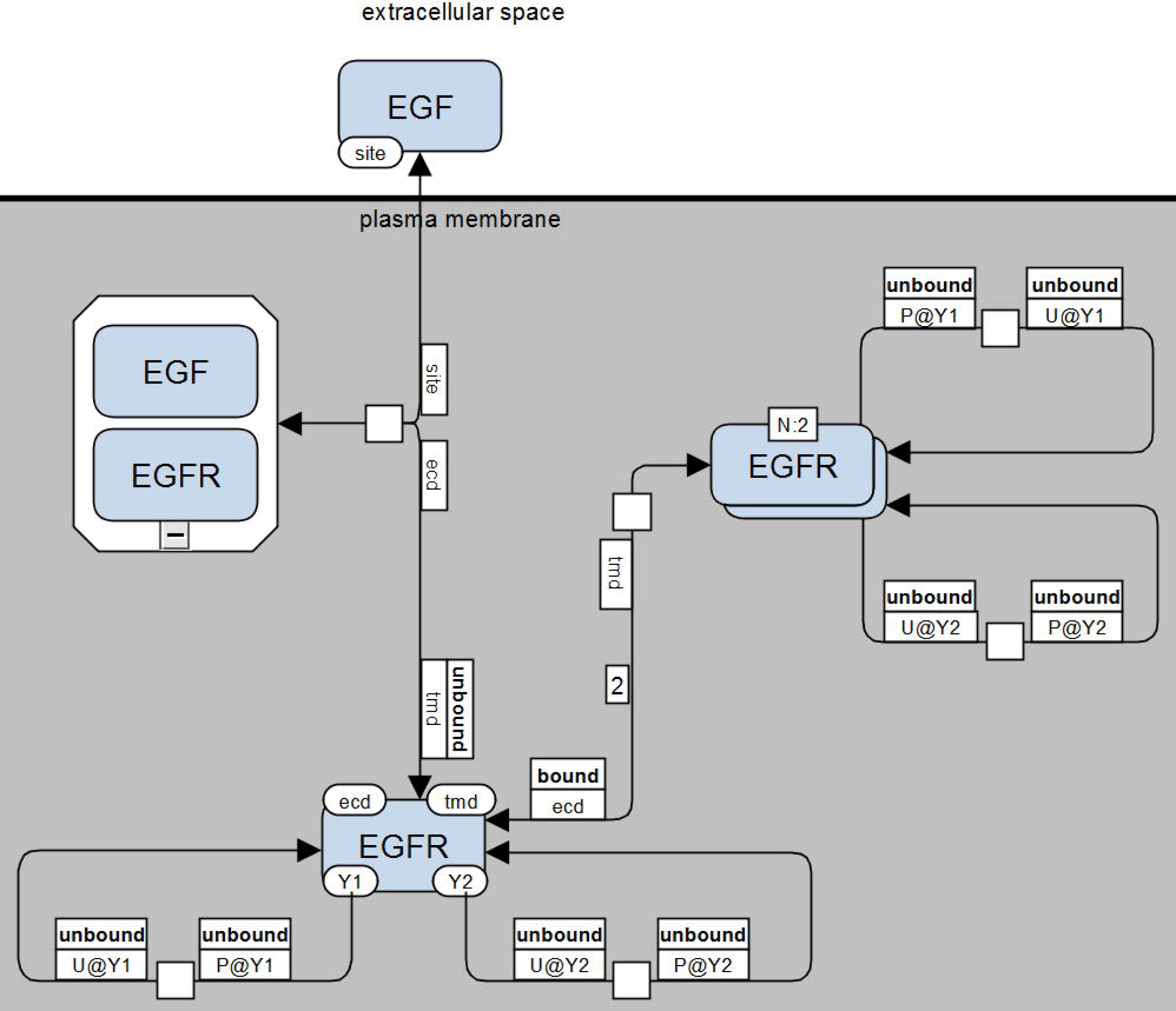
Fully-detailed visualization of the Molecular Process Diagram explicitly showing that for the EGF-EGFR binding reaction the “tmd” sites of EGFR must be unbound and for the EGFR dimerization reaction the “ecd” sites of EGFR must be bound.

**Figure 10.**
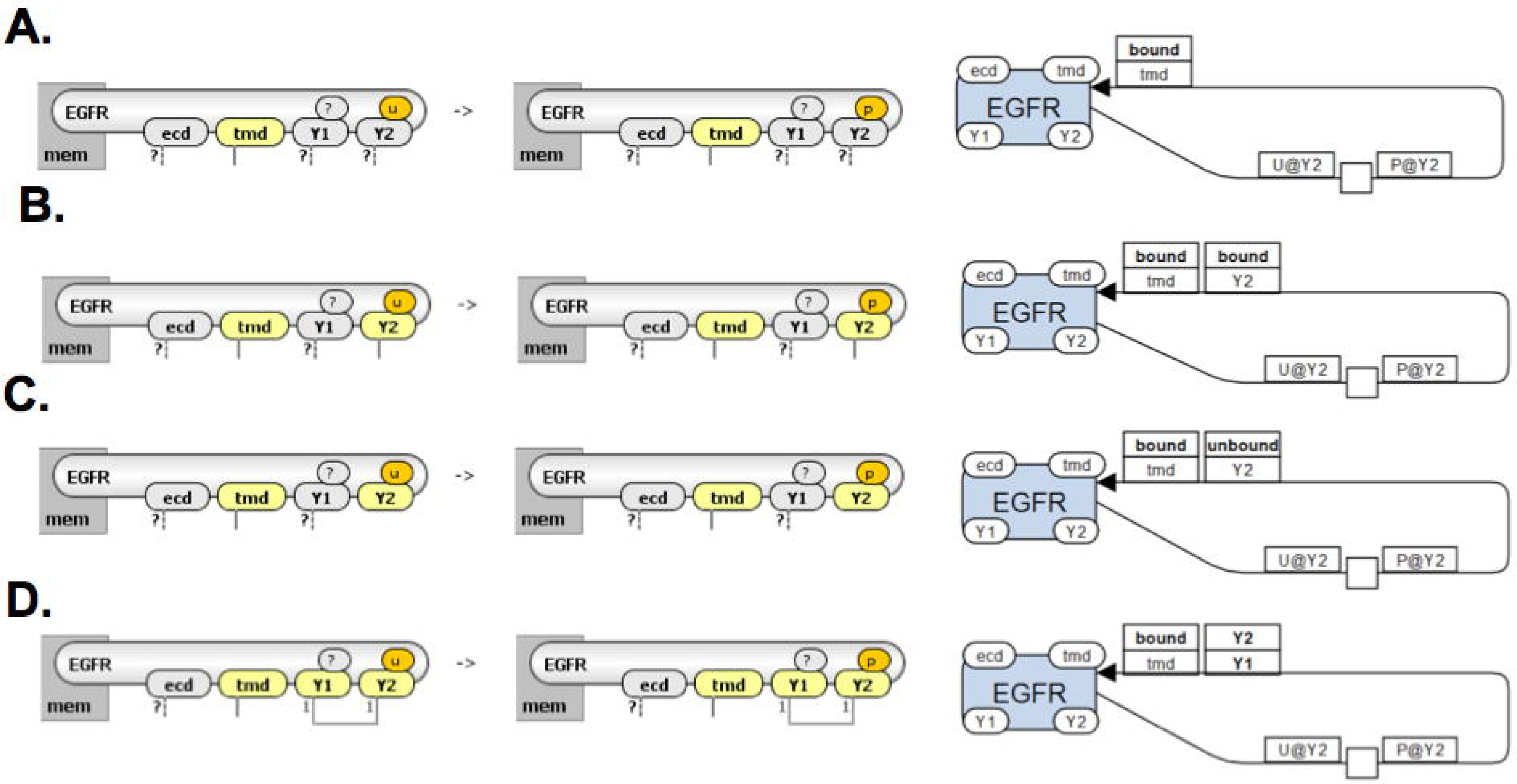
Use of site information boxes to depict different conditions on transphosphorylation reaction. All rules have a condition that transmembrane domain of EGFR (tmd) is bound. Additionally, in rule B tyrosine Y2 must be bound, in rule C tyrosine Y2 must be unbound, in rule D (hypothetical situation) conformational change lead to Y1 and Y2 being occupied together.

We should emphasize that the principles of our proposed visualization approach are simple: every reaction rule defines a transformation of molecules, thus we just need to graphically represent the changed sites before and after as well as the sites that have specific requirements for the transformation. However, the devil is in details: how can we display all site-specific information so that it will be uniquely identifiable, but compact and not cluttering the view? The arrows that connect molecular boxes with reaction rule nodes are an ideal place to include information about sites. Placing information about molecular sites on the arcs at both ends of a reaction rule glyph allows for the relationships to be defined by the type of the arc: molecular sites on consumption arcs belong to reactant molecules, and sites on production arcs belong to the product molecules. Since the context requirements of a rule belong to the molecules and not the rule, it is logical to place that information next to the molecular glyphs instead of the arcs.

Additional difficulties remain: first, when referring to a molecular site outside of a molecule, we need to uniquely associate the site with the corresponding molecule; and second, rules have a lot of redundant information (e.g. the same molecular context is repeated for both reactant and product molecules) which ideally should not be repeated in the visual representation.

To solve the first problem we link the site to its parent molecule by using a site%molecule symbol (note that in the pursuit of compactness, such marking is being used only when distinct molecules have sites with the same name; if this is not the case, the molecule identity can be uniquely recovered just by the site name). There are also situations where two or more molecules of the same type in a complex have distinct roles, in which case all of their sites must be identified as belonging to the particular instance of the molecule in the complex. For example, a model for early events in FceRI signaling (53,54) has complexes that consist of two identical receptors linked to a bivalent ligand, and two Syk kinases that transphosphorylate each other. The phosphorylation events are governed by contingencies that separately specify the states of the sites on each of the two receptor molecules. To graphically represent this rule, we introduce a site index placed into an information box that assigns a site to a specific molecule instance of the complex (see **Fig. 11**; note that there is no need to label the actual molecules in the diagram).

**Figure 11.**
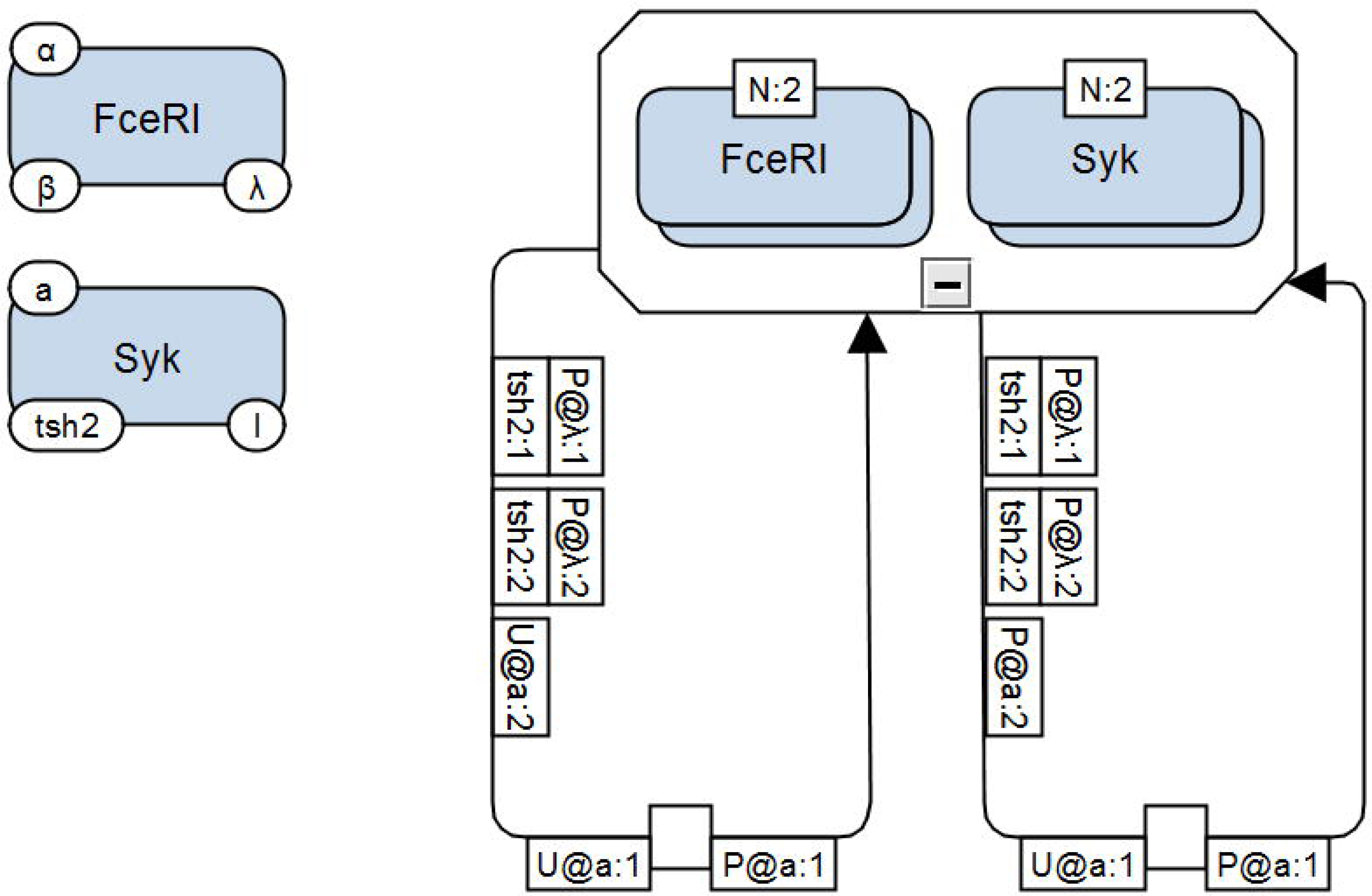
Fully-detailed Molecular Process Diagram for the two rules of Syk transphosphorylation by another Syk kinase. The molecules involved in the rules are two receptors and two Syk kinases. Both receptors have gamma chains phosphorylated and bound to tandem SH2 domains of two distinct Syk kinases. Both rules change the state of site “a” of one Syk kinase from “U” to “P”. Two rules are required in the model because the rate of phosphorylation is different depending on whether or not the site “a” of the other Syk molecule in the complex is phosphorylated or not. To express this contingency the two rules differ by requiring either “P” or “U” in the state of the site “a” of the second kinase, respectively. SWe do not care whether Syk tytosine is bound or unbound.

We should note that the identification issue above must be solved in order to maintain an unequivocal translation between the graphical representation and the rules; however, solving the second problem related to redundancy is somewhat optional: it is desirable, but not required. We did address it by introducing several possible simplifications. First, a site defining a molecular context should be shown only once next to a molecular complex node wherever it creates less visual congestion. Second, we can eliminate some of the information related to binding/unbinding transformations: in simple rules of binding-unbinding it is enough to show two unbound sites to be bound OR bound sites after binding. Choosing one option over the other would depend on which one allows for a more convenient/simple visual representation, but typically we would use the latter: the sites before binding could be marked with the “unbound” attribute, but this is omitted because there is no way the sites before binding can be in any state but unbound. However, in complicated rules where binding is coupled with a change of state, one needs to show site context on both sides of the reaction rule node (see examples in Supplementary Material).

A final point to be made is that SBGN is color-agnostic. Nevertheless, to make diagrams more easily readable, we did assign colors to the molecules, and used site information boxes of the same color as the molecule they belong to. To simplify matching of MPD representations with Virtual Cell diagrams, we chose to make glyphs for rule nodes yellow and placed the rule number inside (see supplemental material for examples).

We must emphasize that the detailed SBGN-compliant visualization of a complete Molecular Process Diagram (with all site-specific information included) has a one-to-one correspondence to the BioNetGen Language (BNGL) description of the model: one can use the SBGN PD diagram to unequivocally reconstruct the BNGL file by converting process nodes to rules, consumption arcs to reactant patterns, production arcs to product patterns, and using information boxes to correctly refine the final BNGL string. The diagrams presented here were done either automatically in VCell software or handcrafted using a general-purpose drawing tool for SBGN-compliant diagrams, yED. The SBGN-style diagrams are all provided electronically as .graphml files in the Supplemental Material. VCell models are provided in the Public Models folder of VCell software (no registration is required to access the folder).

Finally, we should point out that the above-mentioned one-to-one correspondence of the visualization to the rule-based description should not be misinterpreted as a claim that the proposed method is able to unequivocally represent any rule based model. The described visualization does cover all modeling constructs based on the foundational definition of reaction rules (transformation of chemical species selected by reactant patterns into chemical species defined by product patterns). However, rule-based languages can have a descriptive powers that extend well beyond such transformations. On the one hand, rules can be decorated with unusual features, such as the exclusion of certain molecular combinations from a set of possible reactants/products or limiting to only a subset of possible reactants/products selected by a rule (see BioNetGen commands *DeleteMolecules, exclude_reactants, exclude_products, include_reactants, include_products*). Such information could be potentially visualized (for instance as a set of additional information boxes), but for now we chose to relegate these extra conditions to future development. On the other hand, rule-based modeling often blurs the line between the declarative elements (the actual modeling assumptions) and prescriptive elements (which are essentially simulation specifications). The described conventions serve as a rule-based analogue of a reaction diagram and thus strive to cover most of the former but not the latter (such as seed species, kinetic laws, limits on species based on molecular stoichiometry, limits on reactions based on the number of iterations when applying rules, choice of simulation algorithms, etc.).

## Conclusions

Dynamical models of biological process that take into account the details of multimolecular complexes which form inside cells are no longer a rarity, as we now have a rapidly increasing body of publicly available data to provide such detailed information. Cartoon diagrams (whether ad-hoc or in a standardized format) historically served as the “gold standard” for visualizing existing knowledge and/or working hypotheses about biological systems, ranging from qualitative description of pathways to quantitative mathematical models. In the latter case, such diagrams strive to present the model of the biological system in a comprehensive but understandable way; therefore, they must be both easily readable and attentive to detail. Unfortunately, despite many recent efforts, this has proven to be a significant challenge in the case of rule-based models – an approach to model building that is nowadays frequently required in order to circumvent the otherwise intractable complexity associated with multiple possible combinations of molecular interactions when all intramolecular binding and/or modification sites are to be considered by the model.

Here we report the concept of a multilayered **Molecular Process Diagram (MPD)** that builds upon the existing Process Diagram concept and provides a *comprehensive, scalable, and intuitive* depiction of rule-based models. An **MPD** can provide the means of visualizing both the ‘big picture’ aspects of a model (molecules and molecular complexes, and the interactions among them), and the specific hypothesized conditions under which the interactions take place (detailed model assumptions). The scalability of the diagrammatic presentation is provided by three layers of information: *interacting molecules, sites modified*, and *molecular context*, allowing navigation from a birds-eye view of the system all the way to digging into the narrowly-defined site-specific context of any particular molecular interaction, which is communicated with precision and clarity as rule decorations.

We have developed a software implementation of the MPD visualization approach for the popular VCell modeling and simulation framework. Besides the benefit of visualization, which allows to easily summarize and assemble all information about interactions of interest, and outline the information flow through the biological system, the associated GUI also provides a new, biologist-friendly method to build rule-based models and specify the details of each rule (including locational information).

Since the Systems Biology Graphical Notation (SBGN) standard includes Process Diagrams as one of its three types of visualizations, we explored the possibilities of representing MPDs in an SBGN PD-compliant way. We show here a prototype approach to use existing SBGN glyphs and associated constructs to capture all three layers of information present in MPDs. The example diagrams shown in Figs. 7–10 were drawn such that stripping the additional site-specific information will always yield a standard SBGN PD cartoon. We must note that the syntax and semantics of SBGN is a community-regulated effort, and therefore the proposed visualization of all additional features present in MPDs need to be discussed among the researchers involved in standardization efforts for visualization. We can envision alternative solutions to extend the current SBGN semantics in support of MPDs, such as the use newly-designed glyphs. However, we believe that our proposed approach to extend SBGN PD is straightforward and involves minimal changes to the standard, and therefore likely to be the preferred path if MPDs were to be accepted by the community.

Finally, we must emphasize that while the MPD concept, as well as the proposed extensions to SBGN PD, were developed specifically to address the challenges of visualizing rule-based models, they can be very helpful whenever there is a need to graphically represent information about site-specific interactions among biomolecules. Mathematical models are easier to understand and evaluate when their assumptions can be easily visualized at various levels of detail and their connections to biological prior knowledge identified. We therefore expect that the ideas presented here will be not only immediately useful for the visualization of (large) rule-based models, but also for more general-purpose visualization of cell signaling systems when capturing protein substructures and site-specific details of protein interactions is of importance. The design of our approach was driven by commonly encountered scenarios when attempting to build large rule-based models of cell signaling systems. However, at present, the development of such models is not routine, and we expect that the guidelines presented here will require refinement in the future. We are dedicated to use and test this approach in our own modeling efforts and incorporate feedback from users of VCell software (which provides a fully functional implementation) and from the SBGN developer and user communities.

## Supporting information

Supplemental Data Description

## Acknowledgements

This work was supported by National Institutes of Health grants P41 GM103313 and R01 GM95485. We thank summer student Tyler McLaughlin for drafting the initial prototype drawings.

## Author Contributions

Conceptualization: Dan Vasilescu, James C. Schaff, Ion I. Moraru, Michael L. Blinov

Formal Analysis: James Greene

Funding Acquisition: Ion I Moraru, Michael L Blinov

Investigation: Michael L Blinov

Methodology: Dan Vasilescu, James C Schaff, Michael L Blinov

Project Administration: Michael L Blinov

Resources: Michael L Blinov.

Software: Dan Vasilescu, James C. Schaff, Ion I Moraru, Michael L. Blinov.

Supervision: Michael L Blinov

Validation: James Greene, Michael L Blinov

Visualization: James Greene, Michael L Blinov

Writing – Original Draft Preparation: James Greene, Michael L Blinov

Writing – Review & Editing: James Greene, Ion I Moraru, Dan Vasilescu, James C Schaff, Michael L Blinov

## References

(1) Pawson T, Nash P. Assembly of cell regulatory systems through protein interaction domains. Science 2003 Apr 18;300(5618):445–452.

(2) Pawson T. Specificity in signal transduction: from phosphotyrosine-SH2 domain interactions to complex cellular systems. Cell 2004;116(2):191–203.

(3) Seet BT, Dikic I, Zhou M, Pawson T. Reading protein modifications with interaction domains. Nature reviews Molecular cell biology 2006;7(7):473.

(4) Olsen JV, Blagoev B, Gnad F, Macek B, Kumar C, Mortensen P, et al. Global, in vivo, and site-specific phosphorylation dynamics in signaling networks. Cell 2006;127(3):635–648.

(5) Kholodenko BN, Demin OV, Moehren G, Hoek JB. Quantification of short term signaling by the epidermal growth factor receptor. J Biol Chem 1999 Oct 15;274(42):30169–30181.

(6) Siraganian RP, Zhang J, Suzuki K, Sada K. Protein tyrosine kinase Syk in mast cell signaling. Mol Immunol 2002;38(16-18):1229–1233.

(7) Chylek LA, Wilson BS, Hlavacek WS. Modeling biomolecular site dynamics in immunoreceptor signaling systems. A Systems Biology Approach to Blood: Springer; 2014. p. 245–262.

(8) Hlavacek WS, Faeder JR, Blinov ML, Posner RG, Hucka M, Fontana W. Rules for modeling signal-transduction systems. Sci.STKE 2006;2006(344):re6–re6.

(9) Mayer BJ, Blinov ML, Loew LM. Molecular machines or pleiomorphic ensembles: signaling complexes revisited. Journal of biology 2009;8(9):81.

(10) Blinov ML, Faeder JR, Goldstein B, Hlavacek WS. BioNetGen: software for rule-based modeling of signal transduction based on the interactions of molecular domains. Bioinformatics 2004;20(17):3289–3291.

(11) Sneddon MW, Faeder JR, Emonet T. Efficient modeling, simulation and coarse-graining of biological complexity with NFsim. Nature methods 2011;8(2):177.

(12) Sekar JA, Faeder JR. Rule-based modeling of signal transduction: a primer. Computational Modeling of Signaling Networks: Springer; 2012. p. 139–218.

(13) Chylek LA, Harris LA, Faeder JR, Hlavacek WS. Modeling for (physical) biologists: an introduction to the rule-based approach. Physical biology 2015;12(4):045007.

(14) Chylek LA, Holowka DA, Baird BA, Hlavacek WS. An interaction library for the FcεRI signaling network. Frontiers in immunology 2014;5:172.

(15) Moraru II, Schaff JC, Slepchenko BM, Blinov M, Morgan F, Lakshminarayana A, et al. Virtual Cell modelling and simulation software environment. IET systems biology 2008;2(5):352–362.

(16) Schaff JC, Vasilescu D, Moraru II, Loew LM, Blinov ML. Rule-based modeling with Virtual Cell. Bioinformatics 2016;32(18):2880–2882.

(17) Le Novere N, Hucka M, Mi H, Moodie S, Schreiber F, Sorokin A, et al. The systems biology graphical notation. Nat Biotechnol 2009;27(8):735.

(18) Mi H, Lazareva-Ulitsky B, Loo R, Kejariwal A, Vandergriff J, Rabkin S, et al. The PANTHER database of protein families, subfamilies, functions and pathways. Nucleic Acids Res 2005;33(Suppl_1):D284–D288.

(19) Hoops S, Sahle S, Gauges R, Lee C, Pahle J, Simus N, et al. COPASI—a complex pathway simulator. Bioinformatics 2006;22(24):3067–3074.

(20) Funahashi A, Morohashi M, Kitano H, Tanimura N. CellDesigner: a process diagram editor for gene-regulatory and biochemical networks. Biosilico 2003;1(5):159–162.

(21) Faeder JR, Blinov ML, Hlavacek WS. Rule-based modeling of biochemical systems with BioNetGen. Systems biology: Springer; 2009. p. 113–167.

(22) Harris LA, Hogg JS, Tapia J, Sekar JA, Gupta S, Korsunsky I, et al. BioNetGen 2.2: advances in rule– based modeling. Bioinformatics 2016;32(21):3366–3368.

(23) Scalable simulation of cellular signaling networks. Asian Symposium on Programming Languages and Systems: Springer; 2007.

(24) Rule-based modelling of cellular signalling. International conference on concurrency theory: Springer; 2007.

(25) Meier-Schellersheim M, Xu X, Angermann B, Kunkel EJ, Jin T, Germain RN. Key role of local regulation in chemosensing revealed by a new molecular interaction-based modeling method. PLoS computational biology 2006;2(7):e82.

(26) Cheng H, Angermann BR, Zhang F, Meier-Schellersheim M. NetworkViewer: visualizing biochemical reaction networks with embedded rendering of molecular interaction rules. BMC systems biology 2014;8(1):70.

(27) Tiger C, Krause F, Cedersund G, Palmér R, Klipp E, Hohmann S, et al. A framework for mapping, visualisation and automatic model creation of signal-transduction networks. Molecular systems biology 2012;8(1):578.

(28) Thomson TM, Benjamin KR, Bush A, Love T, Pincus D, Resnekov O, et al. Scaffold number in yeast signaling system sets tradeoff between system output and dynamic range. Proc Natl Acad Sci U S A 2011 Dec 13;108(50):20265–20270.

(29) Lopez CF, Muhlich JL, Bachman JA, Sorger PK. Programming biological models in Python using PySB. Molecular systems biology 2013;9(1):646.

(30) Blinov ML, Faeder JR, Goldstein B, Hlavacek WS. A network model of early events in epidermal growth factor receptor signaling that accounts for combinatorial complexity. BioSystems 2006;83(2-3):136–151.

(31) Blinov ML, Yang J, Faeder JR, Hlavacek WS. Depicting signaling cascades. Nat Biotechnol 2006;24(2):137.

(32) Graphical rule-based representation of signal-transduction networks. Proceedings of the 2005 ACM symposium on Applied computing: ACM; 2005.

(33) Blinov ML, Yang J, Faeder JR, Hlavacek WS. Graph theory for rule-based modeling of biochemical networks. Transactions on Computational Systems Biology VII: Springer; 2006. p. 89–106.

(34) Zhang F, Angermann BR, Meier-Schellersheim M. The Simmune Modeler visual interface for creating signaling networks based on bi-molecular interactions. Bioinformatics 2013;29(9):1229–1230.

(35) Blinov ML, Moraru II. Leveraging modeling approaches: reaction networks and rules. Advances in Systems Biology: Springer; 2012. p. 517–530.

(36) Kohn KW, Aladjem MI, Kim S, Weinstein JN, Pommier Y. Depicting combinatorial complexity with the molecular interaction map notation. Mol Syst Biol 2006;2:51.

(37) Kohn KW. Molecular interaction maps as information organizers and simulation guides. Chaos: An Interdisciplinary Journal of Nonlinear Science 2001;11(1):84–97.

(38) Chylek LA, Hu B, Blinov ML, Emonet T, Faeder JR, Goldstein B, et al. Guidelines for visualizing and annotating rule-based models. Molecular BioSystems 2011;7(10):2779–2795.

(39) Smith AM, Xu W, Sun Y, Faeder JR, Marai GE. RuleBender: integrated modeling, simulation and visualization for rule-based intracellular biochemistry. BMC Bioinformatics 2012;13(8):S3.

(40) Sekar JAP, Tapia J, Faeder JR. Automated visualization of rule-based models. PLoS computational biology 2017;13(11):e1005857.

(41) Siebenhaller M, Nielsen SS, McGee F, Balaur I, Auffray C, Mazein A. Human-like layout algorithms for signalling hypergraphs: outlining requirements. Briefings in bioinformatics 2018.

(42) Joshi-Tope G, Gillespie M, Vastrik I, D’Eustachio P, Schmidt E, de Bono B, et al. Reactome: a knowledgebase of biological pathways. Nucleic Acids Res 2005;33(suppl_1):D428–D432.

(43) Kolpakov F, Poroikov V, Selivanova G, Kel A. GeneXplain—Identification of Causal Biomarkers and Drug Targets in Personalized Cancer Pathways. Journal of biomolecular techniques: JBT 2011;22(Suppl):S16.

(44) Olivier BG, Snoep JL. Web-based kinetic modelling using JWS Online. Bioinformatics 2004;20(13):2143–2144.

(45) BioUML: visual modeling, automated code generation and simulation of biological systems. Proceedings of The Fifth International Conference on Bioinformatics of Genome Regulation and Structure; 2006.

(46) Calzone L, Fages F, Soliman S. BIOCHAM: an environment for modeling biological systems and formalizing experimental knowledge. Bioinformatics 2006;22(14):1805–1807.

(47) Demir E, Cary MP, Paley S, Fukuda K, Lemer C, Vastrik I, et al. The BioPAX community standard for pathway data sharing. Nat Biotechnol 2010;28(9):935.

(48) Dilek A, Belviranli ME, Dogrusoz U. VISIBIOweb: visualization and layout services for BioPAX pathway models. Nucleic Acids Res 2010;38(suppl_2):W150–W154.

(49) Babur Ö, Dogrusoz U, Çakır M, Aksoy BA, Schultz N, Sander C, et al. Integrating biological pathways and genomic profiles with ChiBE 2. BMC Genomics 2014;15(1):642.

(50) Gonçalves E, van Iersel M, Saez-Rodriguez J. CySBGN: a cytoscape plug-in to integrate SBGN maps. BMC Bioinformatics 2013;14(1):17.

(51) Sari M, Bahceci I, Dogrusoz U, Sumer SO, Aksoy BA, Babur Ö, et al. SBGNViz: a tool for visualization and complexity management of SBGN process description maps. PloS one 2015;10(6):e0128985.

(52) Van Iersel MP, Villéger AC, Czauderna T, Boyd SE, Bergmann FT, Luna A, et al. Software support for SBGN maps: SBGN-ML and LibSBGN. Bioinformatics 2012;28(15):2016–2021.

(53) Goldstein B, Faeder JR, Hlavacek WS, Blinov ML, Redondo A, Wofsy C. Modeling the early signaling events mediated by FcεRI. Mol Immunol 2002;38(16-18):1213–1219.

(54) Faeder JR, Hlavacek WS, Reischl I, Blinov ML, Metzger H, Redondo A, et al. Investigation of early events in FcεRI-mediated signaling using a detailed mathematical model. The Journal of Immunology 2003;170(7):3769–3781.

