## Supplemental Data Description for "Molecular Process Diagram: a precise, scalable and compact visualization of rule-based models"

The supplemental material shows examples of different visualizations of several published rule-based models. Each model is in a folder ***Model_year*** that contain the following files:

1. A file with the BNGL code (*name_year.txt*).
2. A simplified Molecular Process Diagram shown in VCell representation (*name_year_VCell.PNG*)
3. A detailed Molecular Process Diagram created in yED, shown both in JPEG (*name_year_L3.jpg*) and graphml (*name_year_L3.graphml*) formats.
4. A simplified Molecular Process Diagram created in yED, again shown both in JPEG (*name_year_L2.jpg*) and graphml (*name_year_L2.graphml*) formats.
5. A schematic visualization of rule cartoons shown in VCell representation (*name_year_rules.PNG*)
6. A model file in VCell XML encoding (*Visual_name_year.vcml*).

Each model can be loaded in the VCell modeling and simulation software (<http://vcell.org>). No registration is required; one can either open up the included .vcml files or one can use the “guest” login to browse the online VCell databse – models are located in the left bottom panel, under VCell DB -> BioModels -> PublicBioModels, scroll down to “user” MolecularPD. To see the visualization for a loaded model, one has to click on grouping molecules button in the Reaction Diagram panel:

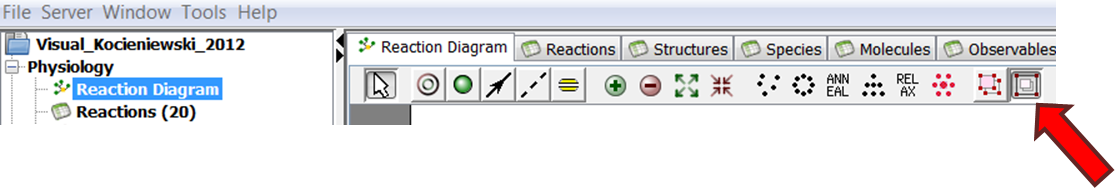

Below we will briefly discuss each model and how such visualization helps to understand its structure. To be consistent with the published models, we didn’t introduce any changes to the BNGL files, even if some rules could be simplified for clarity (e.g., get rid of “unbound” property for sites where it is irrelevant, or renaming sites to be more meaningful using shorter or more unique names).

While the use of colors in SBGN is optional, we have used them to enhance the diagrams; our convention was that sites have the same color as the molecule they belong to. Stripping colors will not fundamentally alter the diagram’s precision and readability.

**BioNetGen Example**. The model from a BioNetGen tutorial shown in this folder has 6 rules involving 3 molecules and a dummy Trash (describing a model of 69 reactions among 17 species). Diagram L2 shows that binding site s of protein S reversibly interacts with binding site t of protein T (rule 1), while gene dnaT produces a protein T (rule 4) that can be degraded into the empty pool Trash (rule 5). Protein S can be phosphorylated (rule 2) and unphosphorylated (rule 3). Two distinct rules account for different conditions for phosphorylation and dephosphorylation. Finally, protein S can dimerize through interaction with site s of another protein S. The detailed diagram L3 shows additional details: Phosphorylation (rule 2) happens only in a dimer (s – bound) and on unprotected tyrosine (tyr – unbound), while dephosphorylation happens on unbound tyrosine (tyr – unbound). An additional contingency (rule 4) specifies that T is generated with unbound site s.

**Kozer_2014**. The paper by Kozer et al., 2014 describes recruitment of the adaptor protein Grb2 to EGFR tetramers. The model illustrated here has 16 rules of interactions among EGF ligand, EGF receptor and Grb2 protein shown in diagram L2. Tetramerization is not evident in this cartoon, as it is regulated by network generation parameters and not by the rules themselves. Nevertheless, one can clearly see some important model features: (a) there are two ways EGFR could dimerize in the absence of ligands (rules 3 and 11), (b) receptor can bind to EGF-EGFR complex (rule 4), (c) EGF binds to a single receptor (rule 0), receptor-dimer (rule 1) or a complex of two EGFR and one EGF (rule 2), (d) there are multiple ways a receptor can be transformed (rules 9-10, 14), and (e) a tetramer can change its intramolecular connectivity in multiple ways (rules 6-8, 12). Note: although the original model is not compartmental, molecular complexes and rule nodes are placed in compartments, consistent with both SBGN conventions and the compartmental rule-based modeling representation in VCell.

**Kocieniewski_2012.** The paper by Kocieniewski et al., 2012 describes an interplay of double phosphorylation and scaffolding in MAPK pathways. The model has 20 rules describing interactions among 4 molecules. Diagram L2 shows that the Scaffold molecule interacts differently with MAPK, MAP2K and MAP3K proteins, each of which, once in a complex with Scaff, can undergo several unimolecular interactions. One can further look at molecular sites that are modified by each rule. For example, all unimolecular transformations are phosphorylation reactions and the detailed diagram L3 describes the contingencies for these interactions. This model illustrates sites associated with molecules: every type of MAPK molecule has sites R1, R2 and S, so each site in a rule that includes several of these molecules must reference back to the molecule (e.g. s:MAPK and s:MAP2K in rules 11-14). However, for rules that include only one molecule, there is no need to clutter the diagram (e.g. in rules 1 and 2 the information box for site s has no reference to a molecule, as the only molecule participating in these rules that has a site s is MAP3K).

**Blinov_2006**. The paper by Blinov et al., 2006 describes early events in epidermal growth factor receptor signaling. The model has 23 rules for interactions among EGF ligand, EGF receptor and Grb2, Shc and Sos proteins which results in a generated reaction network of 354 molecular species connected through 3749 unidirectional reactions. Diagram L2 illustrates these rules, while a diagram L3 provides the context details for these interactions; one can easily identify the most important molecular complexes that affect signaling. Note: although the original model is not compartmental, molecular complexes and rule nodes are placed in compartments, consistent with both SBGN conventions the compartmental rule-based modeling representation in VCell.

**Ligon_2014**. The paper by Ligon et al., 2014 describes a multi-level kinetic model of mRNA delivery via transfection of lipoplexes. The model has 33 rules describing interactions among 6 molecules (plus two “dummy molecules” Trash and Source). Diagram L2 shows that there are 11 forms of external lipoplex bound to pit (illustrated as a complex Lext-Pit) that can be lysed in an endosome with one lipoplex (molecule Lint), and that there is sequential attachment of external lipoplexes to clathrin-coated pit (rules 1-7). A detailed diagram L3 describes contingencies for these interactions. The diagram for this model illustrates that when the number of contingencies is large, they can be placed on both arcs connecting the molecule (or a complex) with the rule node. Note: in rules for Pit endocytosis (rules 8-11), there is no need to have the p site reference a specific external lipoplex, as all of them are identical.

**Dushek_2014**. The paper by Duskek et al., 2014 describes FRET biosensors. The model has 6 rules for interactions among 3 molecules which produce a network of 148 reactions involving 25 species. Diagram L2 shows that the biosensor (B) can reversibly bind the kinase (E) and the phosphatase (F) (note the use of separate rules for forward and reverse interactions in order to specify different requirements) while biosensors can dimerize and undergo modifications. This diagram illustrates complicated binding rules, when binding and site modifications happen at the same time. Rule 0 describes binding (with no state change) of E to B and the simultaneous change of biosensors’ state. As sites to be bound do not change their state, they are shown only once before the rule. Rule 1 describes binding of B to E while biosensors’ binding site is changing its state.

**Barua_2009**. The paper by Barua et al., 2009 describes a mechanism for Jak trans-phosphorylation through interaction with the Src homology 2 (SH2) domain of SH2-B, a unique adaptor protein with the capacity to homo-dimerize.

**Szymanska_2015**. The paper by Szymanska et al., 2015 describes an autophagy/translation switch based on mutual inhibition of MTORC1 and ULK1. The model describes 29 interactions among 7 molecules. Diagram L2 demonstrates that RPTOR may form complexes with MTOR, ULK1 and EIF4EBP1. RPTOR may undergo multiple dephosphorylations (rules 21-23) and multiple phosphorylations (rules 11-13) while in a complex with ULK1. Diagram L3 specifies which sites of ULK1 have to be phosphorylated to enable phosphorylation reactions 11-13.

**Faeder_2003**. The paper by Faeder et al., 2003 describes early events in FcεRI-mediated signaling. The model has 19 rules for interactions among a bivalent ligand, a monovalent FceRI receptor, and two kinases Lyn and Syk. The expanded reaction network consists of 3680 reactions involving 354 molecular species. Diagram L2 illustrates the rules and diagram L3 illustrates the contingencies. Note that many molecular complexes contain multiple instances of identical molecules, and therefore many sites have indices indicating the molecule they belong to. Because interactions within a molecular complex often depend on the connectivity of molecules within the complex, the number of contingencies for each rule can be rather long (see for example rules 4, 5, 7, 8, 10-13).

**Hat_2016**. The paper by Hat et al., 2016 describes the complex circuitry of the p53 network regulated by Mdm2, Wip1, Wip1 and PTEN. The model has 58 rules specifying interactions among 31 molecules, generating 104 reactions involving 54 species.

**Pekalski_2013**. The paper by Pekalski et al., 2013 describes regulation of innate immune response by NF-κB and NF-κB inducible inhibitors, IκBα and A20. The model has 40 rules specifying interactions among 13 molecules.

**Barua_2013**. The paper by Barua and Hlavacek, 2013 describes truncation of APC regulated by a multiprotein complex comprised of APC, Axin, β-catenin, and serine/threonine kinases CK1α and GSK -3β. The model has 29 rules specifying interactions among 5 molecules (and two “dummy molecules” Dead and I).

**Stites_2015_VCell**. The paper by Stites et al., 2015 describes phosphorylation of multiple EGFR tyrosines and assembly of adapter proteins. The model has 99 rules specifying interactions among EGF ligand, EGF receptor and 34 adapter proteins. The VCell model Visual_Stites_2015 provides a coarse-grained visualization of these interactions where due to the large number of rules, site-specific information is omitted.

**Creamer_2012_VCell**. The paper by Creamer et al., 2012 describes signaling events in ERBB signaling leading to activation of ERK and Akt. It tracks phosphorylation of 55 individual serine, threonine, and tyrosine residues, and includes 544 rules specifying interactions among 19 molecules. Even such a huge model can be visualized effectively using the different graph layout algorithms that we developed, providing a connectivity overview that can highlight the importance of specific molecular complexes.

**References**

1. Barua D, Faeder JR, Haugh JM. A bipolar clamp mechanism for activation of Jak-family protein tyrosine kinases. PLoS Computational Biology. 2009 Apr 17;5(4):e1000364.
2. Barua D, Hlavacek WS. Modeling the effect of APC truncation on destruction complex function in colorectal cancer cells. PLoS computational biology. 2013 Sep 26;9(9):e1003217.
3. Blinov ML, Faeder JR, Goldstein B, Hlavacek WS. A network model of early events in epidermal growth factor receptor signaling that accounts for combinatorial complexity. Biosystems. 2006 Feb 1;83(2-3):136-51.
4. Creamer MS, Stites EC, Aziz M, Cahill JA, Tan CW, Berens ME, Han H, Bussey KJ, Von Hoff DD, Hlavacek WS, Posner RG. Specification, annotation, visualization and simulation of a large rule-based model for ERBB receptor signaling. BMC systems biology. 2012 Dec;6(1):107.
5. Dushek O, Lellouch AC, Vaux DJ, Shahrezaei V. Biosensor architectures for high-fidelity reporting of cellular signaling. Biophysical journal. 2014 Aug 5;107(3):773-82.
6. Faeder JR, Hlavacek WS, Reischl I, Blinov ML, Metzger H, Redondo A, Wofsy C, Goldstein B. Investigation of early events in FcεRI-mediated signaling using a detailed mathematical model. The Journal of Immunology. 2003 Apr 1;170(7):3769-81.
7. Kocieniewski P, Faeder JR, Lipniacki T. The interplay of double phosphorylation and scaffolding in MAPK pathways. Journal of Theoretical Biology. 2012 Feb 21;295:116-24.
8. Kozer N, Barua D, Henderson C, Nice EC, Burgess AW, Hlavacek WS, Clayton AH. Recruitment of the adaptor protein Grb2 to EGFR tetramers. Biochemistry. 2014 Apr 21;53(16):2594-604.
9. Ligon TS, Leonhardt C, Rädler JO. Multi-level kinetic model of mRNA delivery via transfection of lipoplexes. PLoS One. 2014 Sep 19;9(9):e107148.
10. Pękalski J, Zuk PJ, Kochańczyk M, Junkin M, Kellogg R, Tay S, Lipniacki T. Spontaneous NF-κB activation by autocrine TNFα signaling: a computational analysis. PloS one. 2013 Nov 11;8(11):e78887.
11. Stites EC, Aziz M, Creamer MS, Von Hoff DD, Posner RG, Hlavacek WS. Use of mechanistic models to integrate and analyze multiple proteomic datasets. Biophysical journal. 2015 Apr 7;108(7):1819-29.
12. Szymańska P, Martin KR, MacKeigan JP, Hlavacek WS, Lipniacki T. Computational analysis of an autophagy/translation switch based on mutual inhibition of MTORC1 and ULK1. PLoS One. 2015 Mar 11;10(3):e0116550.
