## Supplemental Data files for "Molecular Process Diagram: a precise, scalable and compact visualization of rule-based models": Kozer2014.txt

#Supplemental file (a plain-text file)
#
#This file is supplementary material for the report indicated below.
#Title: Recruitment of the adaptor protein Grb2 to EGFR tetramers
#Authors: Noga Kozer, Dipak Barua, Christine Henderson, Eduoard C. Nice, 
#         Antony W. Burgess, William S. Hlavacek and Andrew H.A. Clayton

#This file is a model specification written in BNGL (Faeder et al., 2009)
#  [Methods Mol. Biol. 500:113-167].
#It can be processed by BioNetGen (http://bionetgen.org).
#The file extension should be .bngl

begin parameters

NA 6.02e14 # [=] number of molecules per nmol
f 0.01 # fraction of cell to consider in simulation, dimensionless
V 5
cellDen 6.0e8 # [=] number of cells per L
Vo 4

#See Kozer et al. (2013) for additional information about parameter values.
#  [Mol. BioSyst. 9:1849-1863]

#It should be noted that BioNetGen treats rate constants as single-site rate constants. Thus, care must be taken to specify such rate constants, i.e., rate constants that do not incorporate statistical factors (which arise, for example, because a reactant has multiple indistinguishable copies of a reactive site) or factors that arise because of reactions that have the form A+A->product(s). See Faeder et al. (2009) for more details.

LT	3.0*NA*Vo                   #  Total ligand concentration (nM). A default concentration of 30 nM was used in cases where the ligand dose was not varied. In dose-response analyses, this concentration is varied in the range 0.001-100 nM.  
RT      0.09*NA*Vo                 #  Total receptor concentration (nM). The value is based on 90,000 EGFR per cell and a density of 6*10^5 cell/mL.  
GT f*9.0e4 # number per cell

K11     4.6                  #  Equilibrium association constant (/nM). The constant characterizes binding of EGF with EGFR, which is not crosslinked with other EGFR via the ectodomain.The value is taken from Macdonald and Pike (2008) [PNAS 105:112-117], where it represents EGF binding with monomeric EGFR.   
k11r	0.02                 #  Dissociation constant (/s). This value is taken from Elleman et al. (2001) [Biochemistry 40:8930-3939]. 
k11f	0.09/(NA*Vo)                 #  Association constant (/nM/s). The value is derived from k11f = K11*k11r. 

K21     5.3                  #  Equilibrium association constant (/nM). The constant characterizes binding of EGF with EGFR that is crosslinked via ectodomain with a second EGFR, and the crosslinked receptors both are free of EGF.The value is taken from Macdonald and Pike (2008). 
k21r	0.02                 #  Single-site dissociation constaint (/s). We assume k21r to be the same as k11r. 
k21f	0.053/(NA*Vo)                #  Association constant (/nM/s). The value is derived from k21f = K21*k21r/2. Division by 2 is a correction to obtain the single-site association rate constant (k21f) from the effective equilibrium association constant (K21).     

K22     0.34                 #  Equilibrium association constant (/nM). The constant characterizes binding of EGF with EGFR that is crosslinked via ectodomain with a second EGFR and the second EGFR is occupied by EGF.The value is taken from Macdonald and Pike (2008). 
k22r	0.2                  #  Single-site dissociation constaint (/s). We assume that k22r is 10-fold larger than k11r. This assumption is consistent with the difference between K11 (or K21) and K22. 
k22f	0.136/(NA*Vo)                #  Single-site association constant in (/nM/s). The value is erived from k22f = 2*K22*k22r. Multiplication with 2 is applied to get single site association rate constant (k22f) from effective equilibrium association constant (K22).  

L20     212                  #  Equlibrium crossliinking constaint (/nM). The constant characterizes ectodomain-ectodomain interaction when both receptors are free from EGF. This parameter value is based on the value of 530 dm^2/nmol reported by Macdonald and Pike (2008) and the following assumptions: a receptor surface density of 1.175e-3 nmol/dm^2 (~20 eGFP-EGFR on average for a beam with a radius of 0.3 microns) for a cell line expressing 30,000 copies of EGFR per cell, receptor density is proportional to total receptor copy number, and a total receptor concentration of approximately 0.009 nM in the experiments of Macdonald and Pike (2008) involving cells expressing 87,000 copies of EGFR per cell (from the x-intercept of the Scatchard plot shown in the inset of Fig. 5A in the paper of Macdonald and Pike). In other words, we assume that the dimensionless quantity L20*RT is approximately 1.8 for cells with 87,000 copies of EGFR per cell. Moreover, we assume that the total number of cells used in the experiments of Macdonald and Pike (for a cell line with 87,000 copies of EGFR per cell) was such that the total concentration of EGFR was approximately 0.009 nM. If we further assume that the total number of cells used was the same in all experiments of Macdonald and Pike, then the dimensional value of L20 (in units consistent with those used for total EGF concentration) is 212 /nM (obtained by dividing the dimesionless quantity L20*RT by RT when RT is expressed in the same units as those used for total EGF concentration).
l20r	1.24                 #  Dissociation rate constant in (/s). The value is taken from Low-Nam et al. (2011) [Nat. Struct. Mol. Biol. 18:1244-1249].
l20f	526/(NA*Vo)                  #  Association rate constant in (/nM/s). Derived from l20f = 2*L20*l20r. Multiplication factor 2 is applied because the reactant pair are the same species. 

L21     244                  #  Equilibrium crosslinking constant (/nM). The constant characterizes ectodomain-ectodomain interaction when only one receptor is bound to EGF. The value is derived from the detailed balance relation, L21 = K21*L20/K11. 
l21r	0.738                #  Dissociation rate constant in (/s). The value is taken from Low-Nam et al. (2011).
l21f    180/(NA*Vo)                  #  Association rate constant (/nM/s). The value is derived from l21f = L21*l21r.

L22     18.0                 #  Equilibrium crosslinking constant (/nM). The constant characterizes ectodomain-ectodomain interaction when both receptors are bound to EGF. The value is derived from the detailed balance relation, L22 = K22*L21/K11. 
l22r	0.272                #  Dissociation rate constant in (/s). The value is taken from Low-Nam et al. (2011).
l22f    9.79/(NA*Vo)                 #  Association rate constant (/nM/s). The value is derived from l22f = 2*L22*l22r. Multiplication factor 2 is applied because the reactant pair are the same species.

k_o     6.0                  #  Rate constant for ligand-mediated activating conformation change in the cytosolic tail of EGFR (/s). The value is obtained from fitting.
k_c     1.6                  #  Rate constant for ligand-independent deactivating conformation change in the cytosolic tail of EGFR (/s). The value is obtained from fitting.    

kaf     15.4/(NA*Vo)                 #  Rate constant for EGFR tail-tail association mediated by EGFR kinase domain and juxtamembrane domains (/s). The value is obtained from fitting. 
kar     8.89                 #  Rate constant for EGFR tail-tail dissociation (/s). The value is obtained from model fitting.  

kp      1                    #  EGFR phosphorylation rate constant (/s). The value is consistent with data in Kleiman et al. (2011) [Mol. Cell 43:723-737].  
kdp     1                    #  EGFR dephoshporylation rate constant (/s). The value is consistent with data in Kleiman et al. (2011). 

chi_r   4.37e+04*(NA*Vo)             #  Enhancement factor for ring-closure reactions (nM). The value is obtained from fitting.  

# The parameter alpha does not influence simulation results; this parameter is used as a scale factor to relate simulation results to the experimental data. It multiplies the quantity (observable) defined below called "Clusters." The product is taken to be directly comparable to relative receptor cluster density measured by Kozer et al. (2013). The value of alpha is obtained from fitting.

KDg 713. # nM from Chook YM, Gish GD, Kay CM, Pai EF, Pawson T (1996) [J Biol Chem 271:30472-30478]
kmg		0.31 # /s from Chook et al. (1996)
kpg (kmg/KDg)/(NA*V) # /nM/s

end parameters


begin molecule types
EGF(rec)
EGFR(back,lig,cd~c~o,Y~u~p)
Grb2(SH2)
end molecule types


begin seed species
EGF(rec)	            LT
EGFR(back,lig,cd~c,Y~u)	    RT
Grb2(SH2) GT
end seed species


begin reaction rules

# EGF binding to EGFR not crosslinked via the ectodomain. 
EGF(rec) + EGFR(back,lig) <-> EGF(rec!1).EGFR(back,lig!1) k11f,k11r

# EGF binding to EGFR that is crosslinked via the ectodomain to a second EGFR and both crosslinked EGFR are free of EGF. 
EGF(rec) + EGFR(back!1,lig).EGFR(back!1,lig) <-> EGF(rec!2).EGFR(back!1,lig!2).EGFR(back!1,lig) k21f,k21r

# EGF binding to EGFR that is crosslinked via the ectodomain to a second EGFR and the second EGFR is bound to EGF.  
EGF(rec) + EGF(rec!1).EGFR(back!2,lig!1).EGFR(back!2,lig) <-> EGF(rec!3).EGF(rec!1).EGFR(back!2,lig!1).EGFR(back!2,lig!3) k22f,k22r

# Ectodomain-ectodomain crosslinking of two EGFR that are not tethered and both EGFR are free of EGF.
EGFR(back,lig) + EGFR(back,lig) <-> EGFR(back!1,lig).EGFR(back!1,lig) l20f,l20r 

# Ectodomain-ectodomain crosslinking of two EGFR that are not tethered and only one EGFR is free of EGF. 
EGFR(back,lig) + EGF(rec!1).EGFR(back,lig!1) <-> EGFR(back!2,lig).EGF(rec!1).EGFR(back!2,lig!1) l21f,l21r

# Ectodomain-ectodomain crosslinking of two EGFR that are not tethered and both EGFR are occupied by EGF. 
EGF(rec!1).EGFR(back,lig!1) + EGF(rec!2).EGFR(back,lig!2) <-> EGF(rec!1).EGFR(back!3,lig!1).EGF(rec!2).EGFR(back!3,lig!2) l22f,l22r

# Ectodomain-ectodomain crosslinking of two EGFR that are tethered and both EGFR are free of EGF (ring-closure reaction)
EGFR(cd~o!4,back,lig).EGFR(cd~o!2,back,lig).EGFR(cd~o!2,back!3).EGFR(cd~o!4,back!3) <-> EGFR(cd~o!4,back!1,lig).EGFR(cd~o!2,back!1,lig).EGFR(cd~o!2,back!3).EGFR(cd~o!4,back!3) chi_r*l20f, l20r

# Ectodomain-ectodomain crosslinking of two EGFR that are tethered and only one EGFR is free of EGF (ring-closure reaction)
EGFR(cd~o!4,back,lig).EGFR(cd~o!2,back,lig!+).EGFR(cd~o!2,back!3).EGFR(cd~o!4,back!3) <-> EGFR(cd~o!4,back!1,lig).EGFR(cd~o!2,back!1,lig!+).EGFR(cd~o!2,back!3).EGFR(cd~o!4,back!3) chi_r*l21f, l21r

# Ectodomain-ectodomain crosslinking of two EGFR that are tethered and both EGFR are occuplied by EGF (ring-closure reaction)
EGFR(cd~o!4,back,lig!+).EGFR(cd~o!2,back,lig!+).EGFR(cd~o!2,back!3).EGFR(cd~o!4,back!3) <-> EGFR(cd~o!4,back!1,lig!+).EGFR(cd~o!2,back!1,lig!+).EGFR(cd~o!2,back!3).EGFR(cd~o!4,back!3) chi_r*l22f, l22r

# EGF-induced activating conformation change of EGFR cytosolic tail
EGFR(lig!+,cd~c) -> EGFR(lig!+,cd~o)  k_o

# EGF-independent deactivating conformatoin change of EGFR cytosolic tail
EGFR(cd~o) -> EGFR(cd~c) k_c

# Crosslinking of conformationally-modified cytosolic tails of EGFR that are not tethered
EGFR(cd~o) + EGFR(cd~o) <-> EGFR(cd~o!1).EGFR(cd~o!1) kaf, kar

# Crosslinking of conformationally-modified cytosolic tails of EGFR that are tethered (ring-closure reaction)
EGFR(cd~o,back!1).EGFR(cd~o!3,back!1).EGFR(cd~o!3,back!2).EGFR(cd~o,back!2) <-> EGFR(cd~o!4,back!1).EGFR(cd~o!3,back!1).EGFR(cd~o!3,back!2).EGFR(cd~o!4,back!2) chi_r*kaf, kar

# EGFR phosphorylation
EGFR(cd~o!1).EGFR(cd~o!1,Y~u) -> EGFR(cd~o!1).EGFR(cd~o!1,Y~p) kp 

# EGFR dephosphorylation
EGFR(Y~p) -> EGFR(Y~u)   kdp

# Grb2 binds phosphorylated receptor
Grb2(SH2) + EGFR(Y~p) <-> Grb2(SH2!1).EGFR(Y~p!1) kpg,kmg

end reaction rules


begin observables

# Unbound EGF
Molecules EGFfree EGF(rec)

# Ligand-free EGFR
Molecules EGFRfree EGFR(lig)

 
Molecules pEGFR  EGFR(Y~p!?)

# Grb2
Molecules Grb2_pEGFR EGFR(Y~p!1).Grb2(SH2!1)

# Free Grb2
Molecules Grb2Free  Grb2(SH2)

# Grb2-EGFR monomer
Species Grb2EGFRMonomer EGFR(cd,back,Y~p!1).Grb2(SH2!1)

# Grb2-EGFR dimer
Species Grb2EGFRDimer1  EGFR(cd,back!1,Y~p!2).Grb2(SH2!2).EGFR(cd,back!1,Y)
Species Grb2EGFRDimer2  EGFR(cd~o!1,back,Y~p!2).Grb2(SH2!2).EGFR(cd~o!1,back,Y)
Species Grb2EGFRDimer3  EGFR(cd,back!1,Y~p!2).Grb2(SH2!2).EGFR(cd,back!1,Y~p!3).Grb2(SH2!3)    
Species Grb2EGFRDimer4  EGFR(cd~o!1,back,Y~p!2).Grb2(SH2!2).EGFR(cd~o!1,back,Y~p!3).Grb2(SH2!3)


end observables

#the max_stoich setting below prohibits oligomers larger than tetramers
generate_network({overwrite=>1,max_stoich=>{EGF=>4,EGFR=>4}});

#simulate_ode({t_end=>120,n_steps=>100}); 
