## Supplementary figures and images for "Molecular Process Diagram: a precise, scalable and compact visualization of rule-based models"

### Barua_2009_L2.jpg

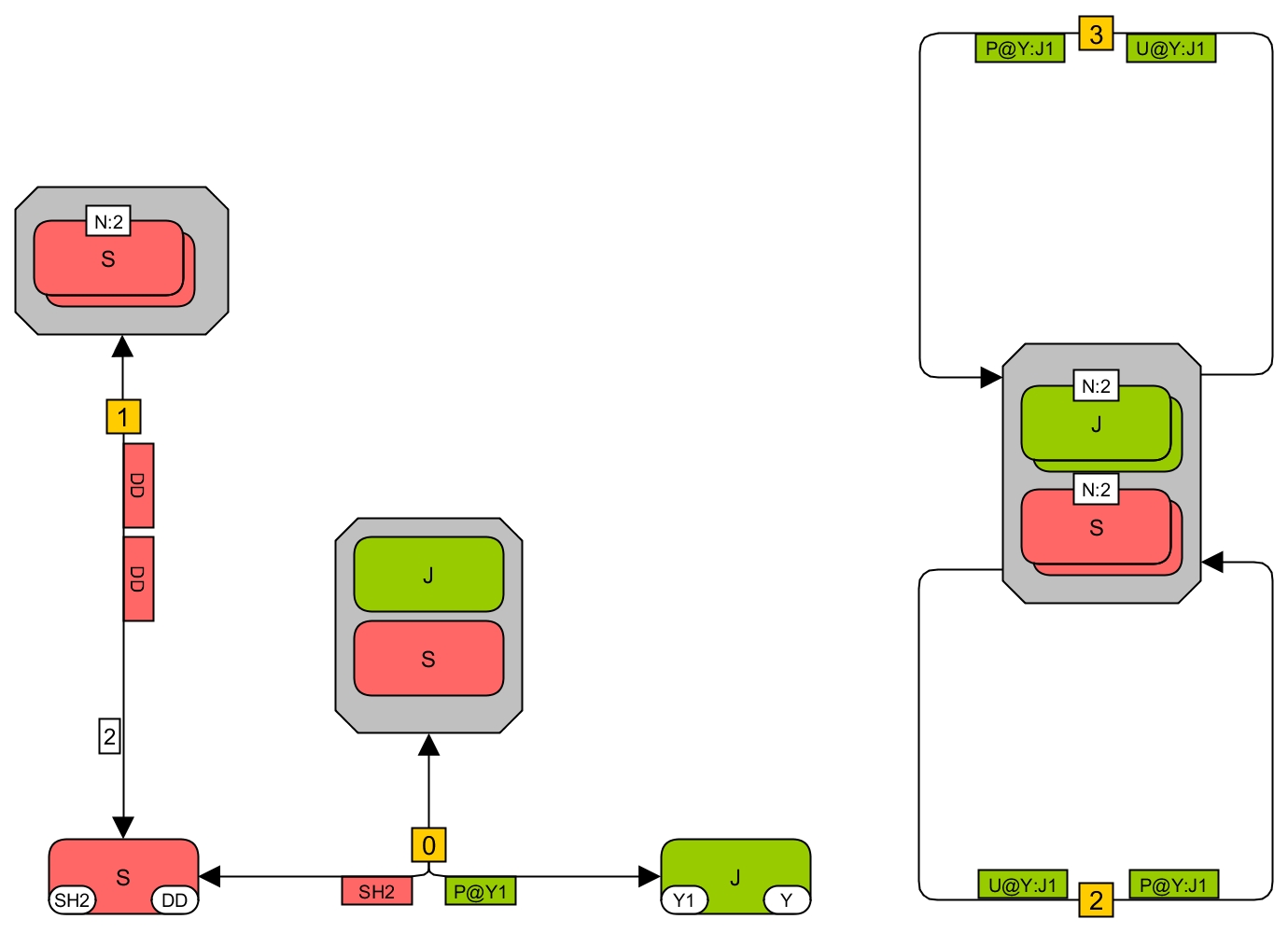

### Barua_2009_L3.jpg

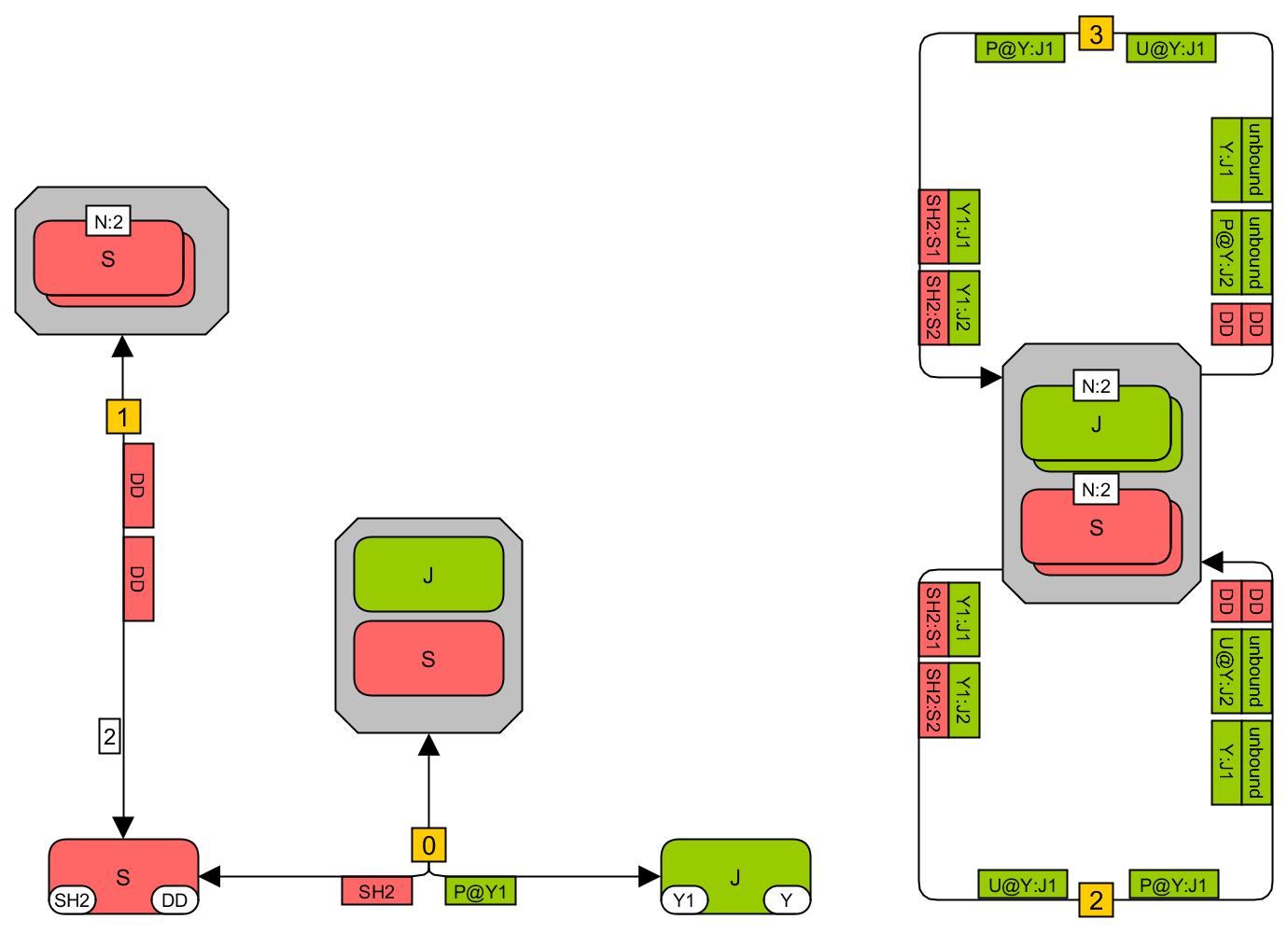

### Barua_2009_rules.PNG

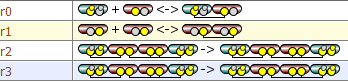

### Barua_2009_VCell.PNG

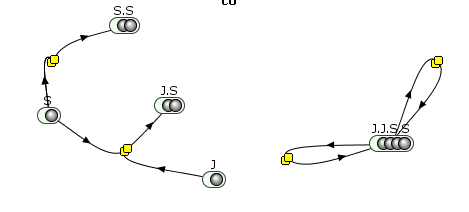

### Barua_2013-L2.jpg

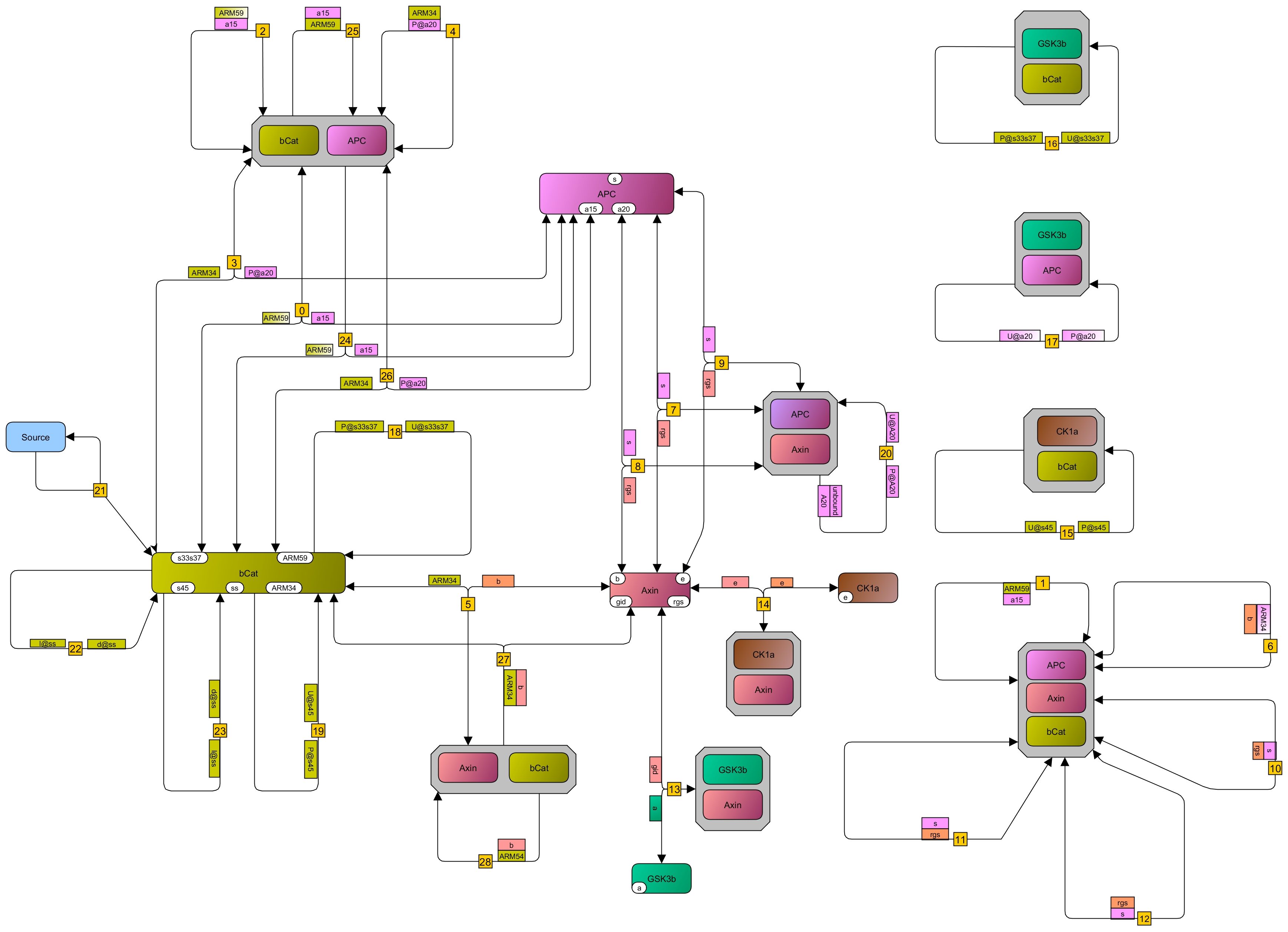

### Barua_2013-L3.jpg

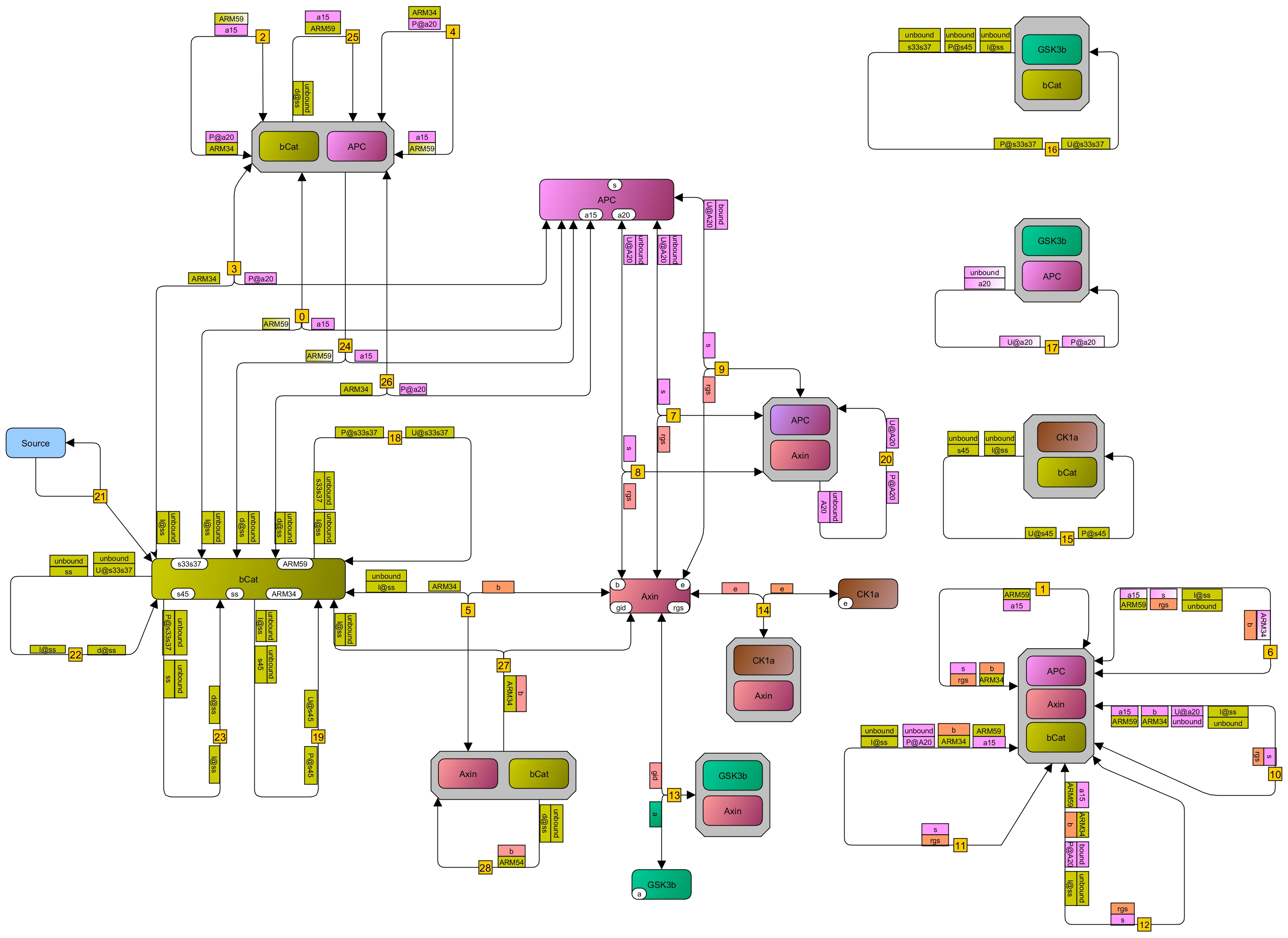

### Barua_2013_rules.PNG

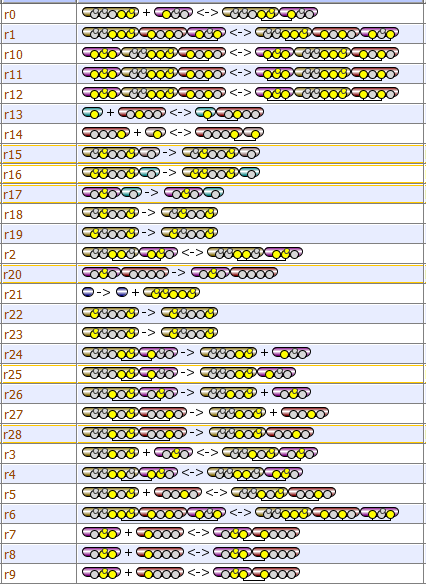

### Barua_2013_VCell.PNG

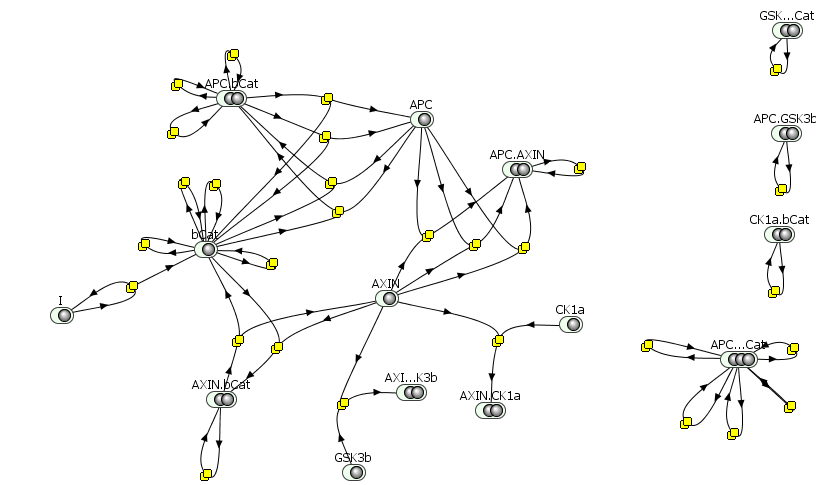

### BioNetGen_example_L2.jpg

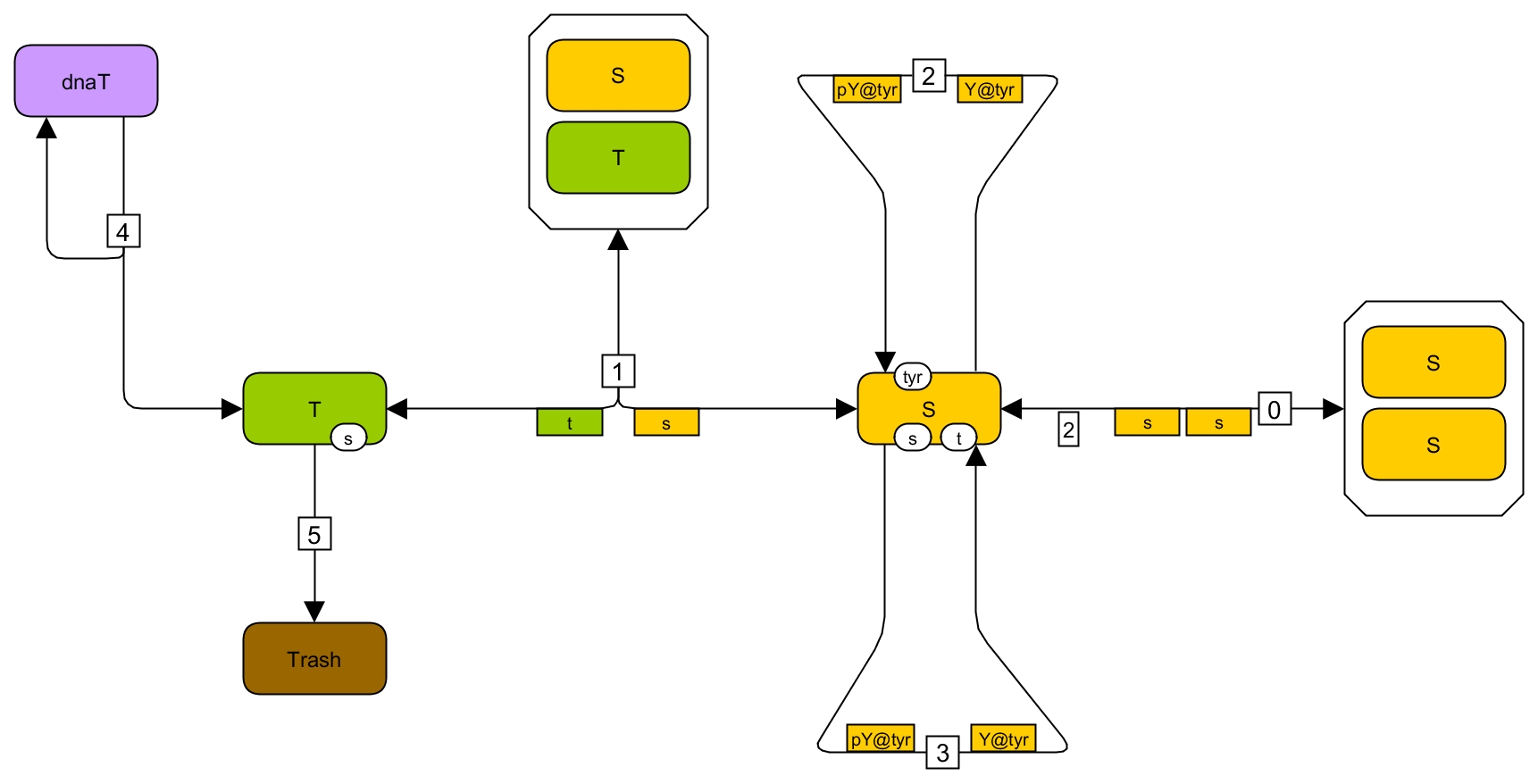

### BioNetGen_example_L3.jpg

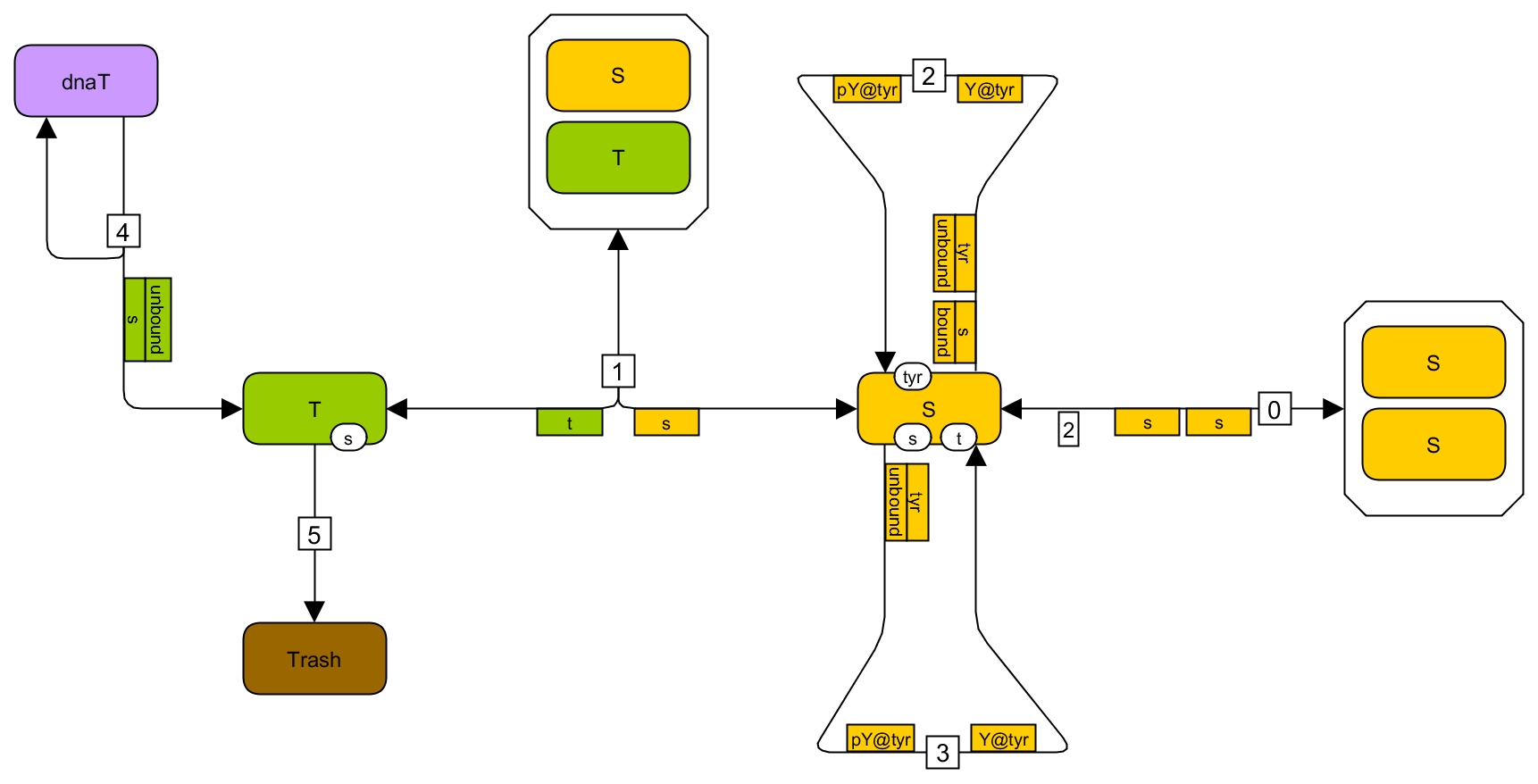

### BioNetGen_example_rules.PNG

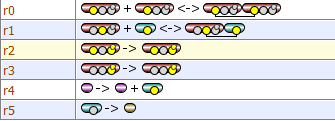

### BioNetGen_example_VCell.PNG

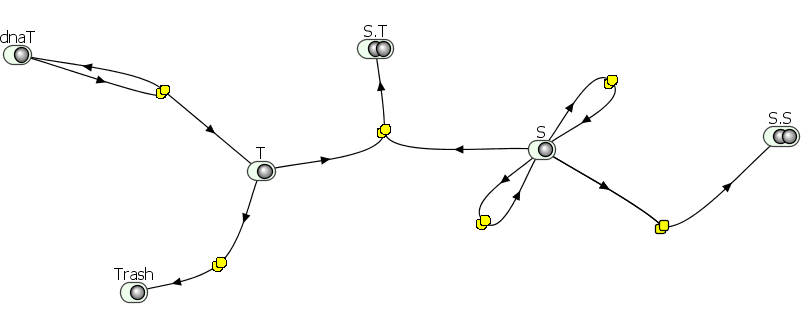

### Blinov_2006_L2.jpg

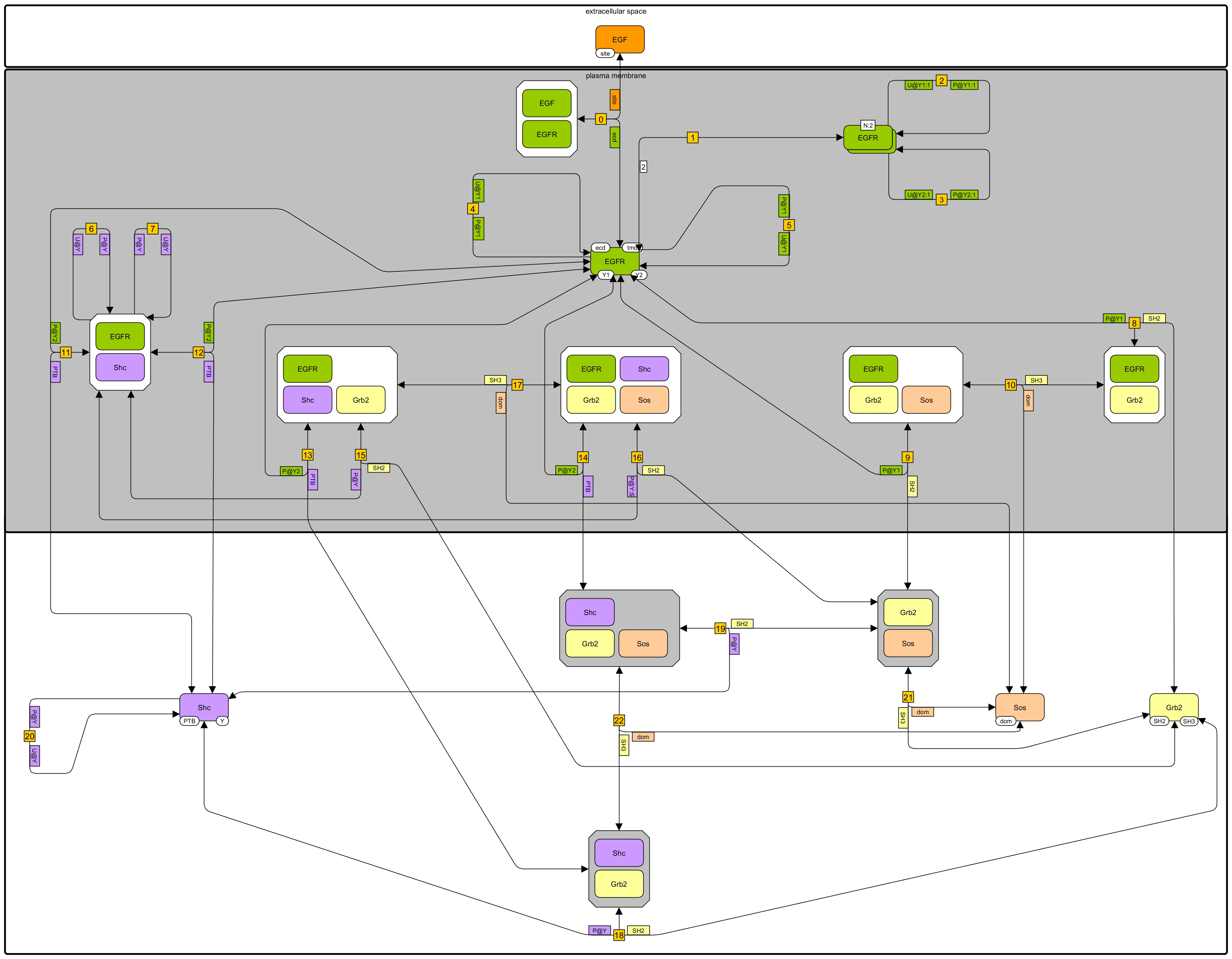

### Blinov_2006_L3.jpg

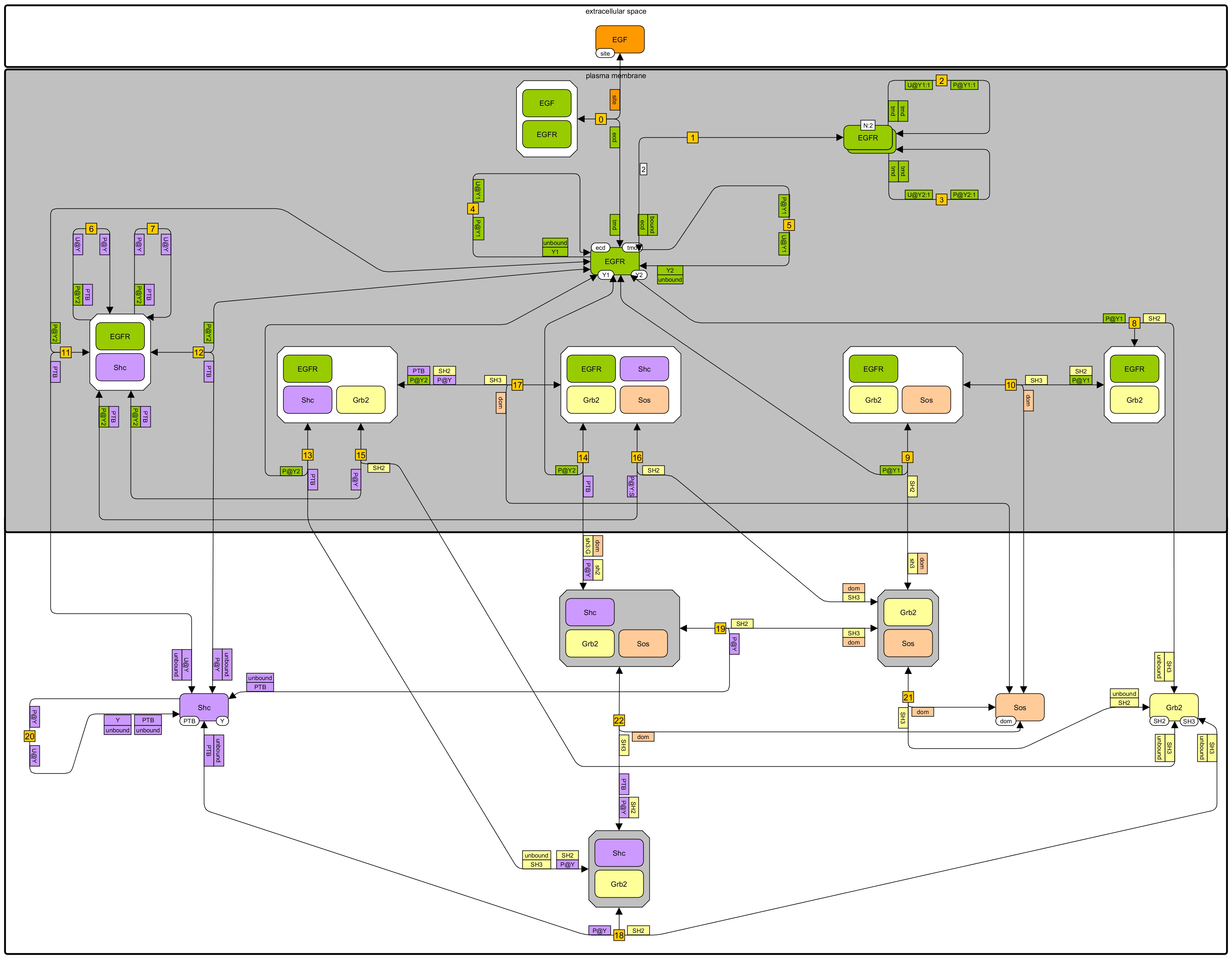

### Blinov_2006_rules.PNG

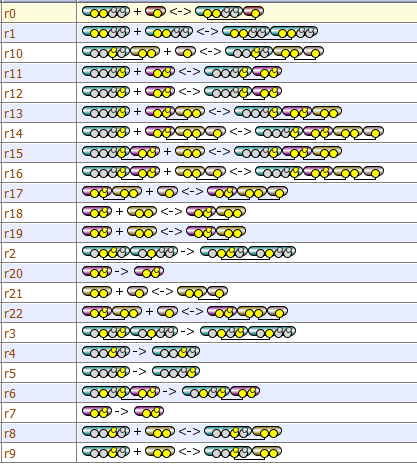

### Blinov_2006_VCell.PNG

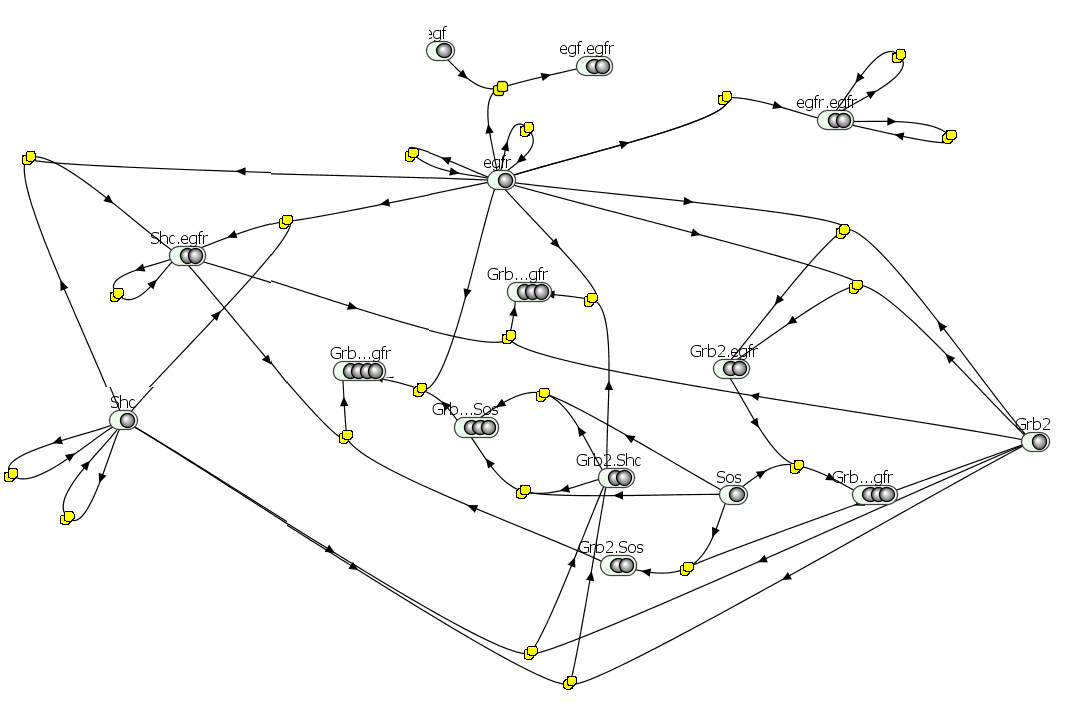

### Creamer_2012_VCell.PNG

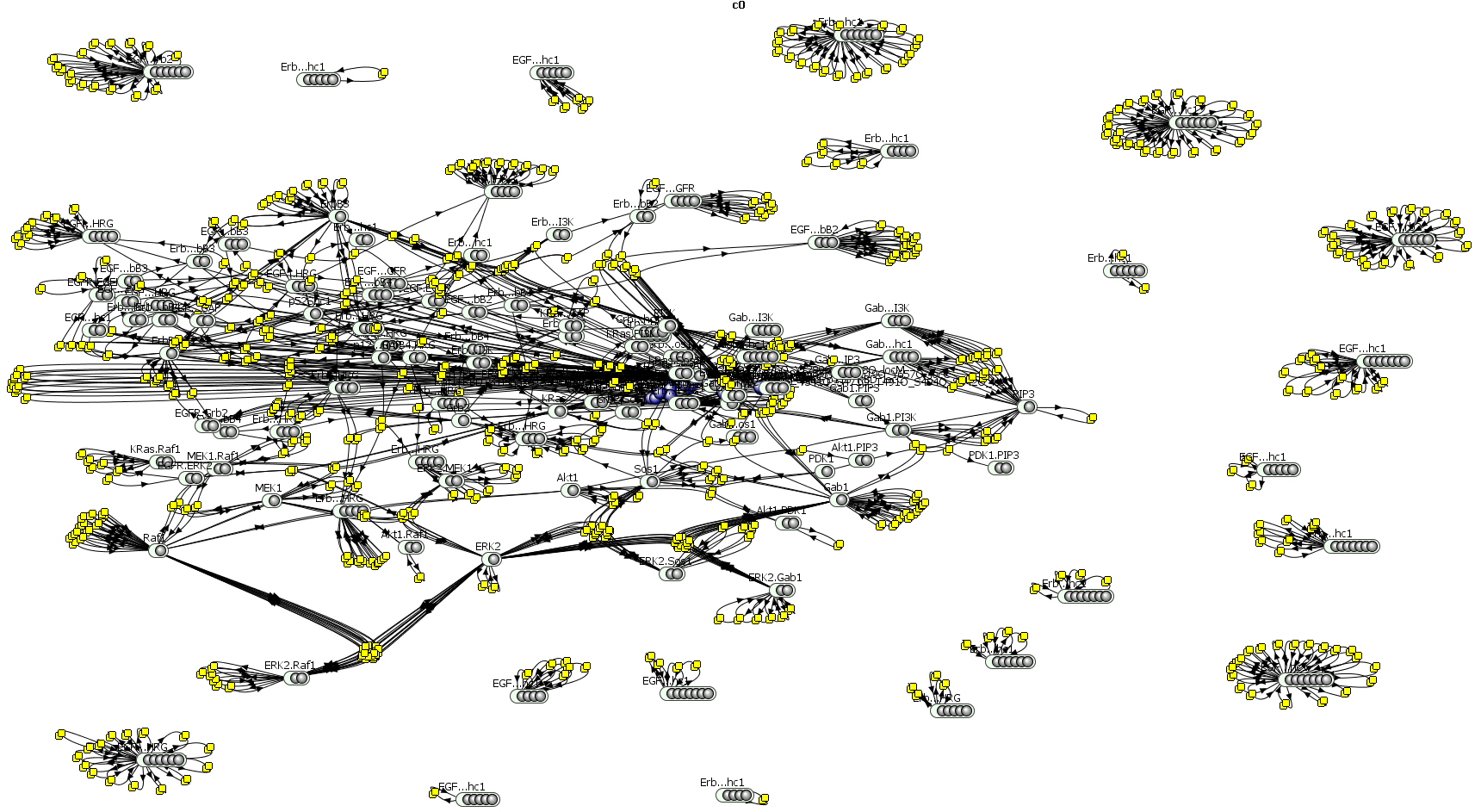

### Dushek_2014_L2.jpg

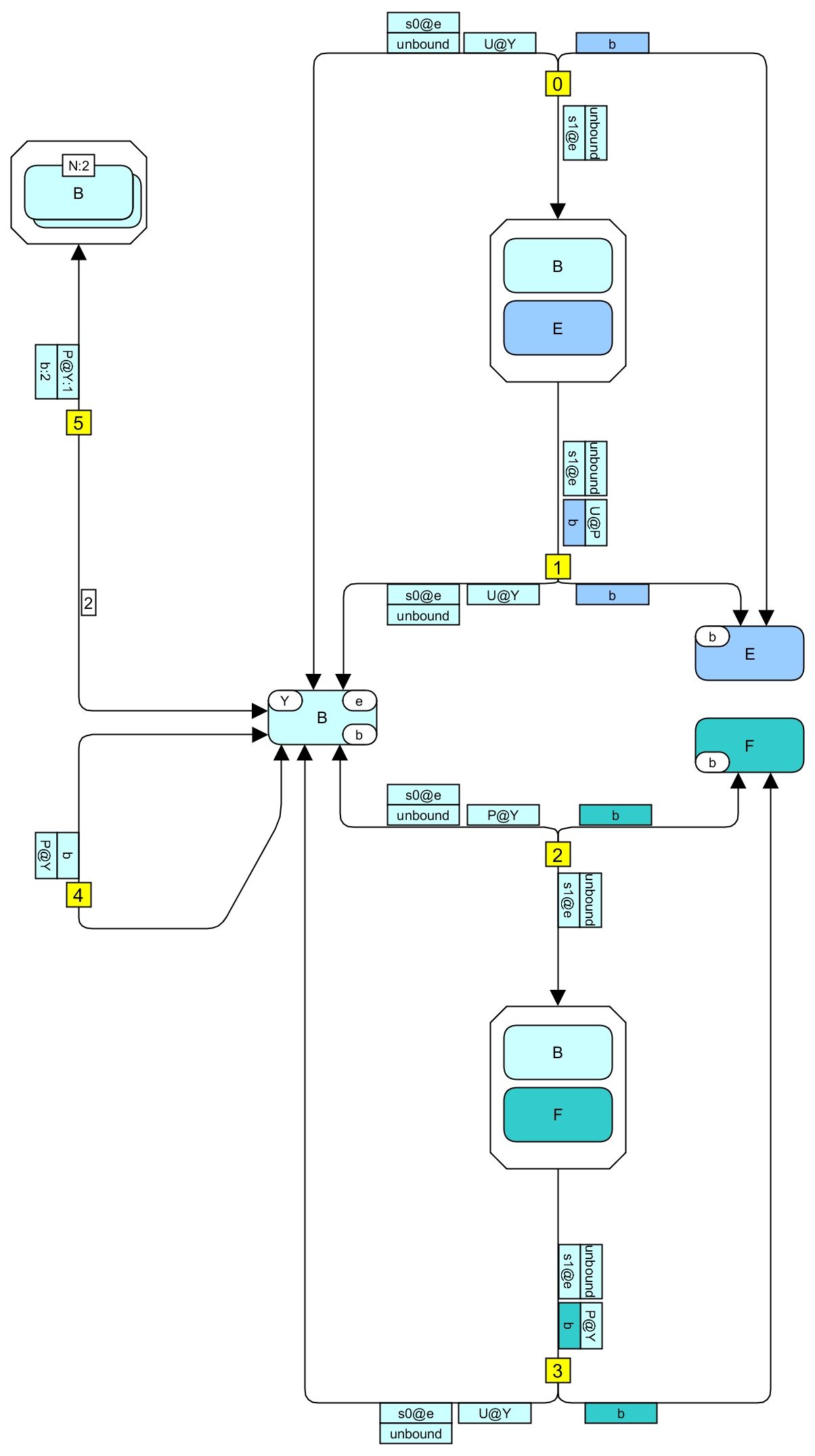

### Dushek_2014_L3.jpg

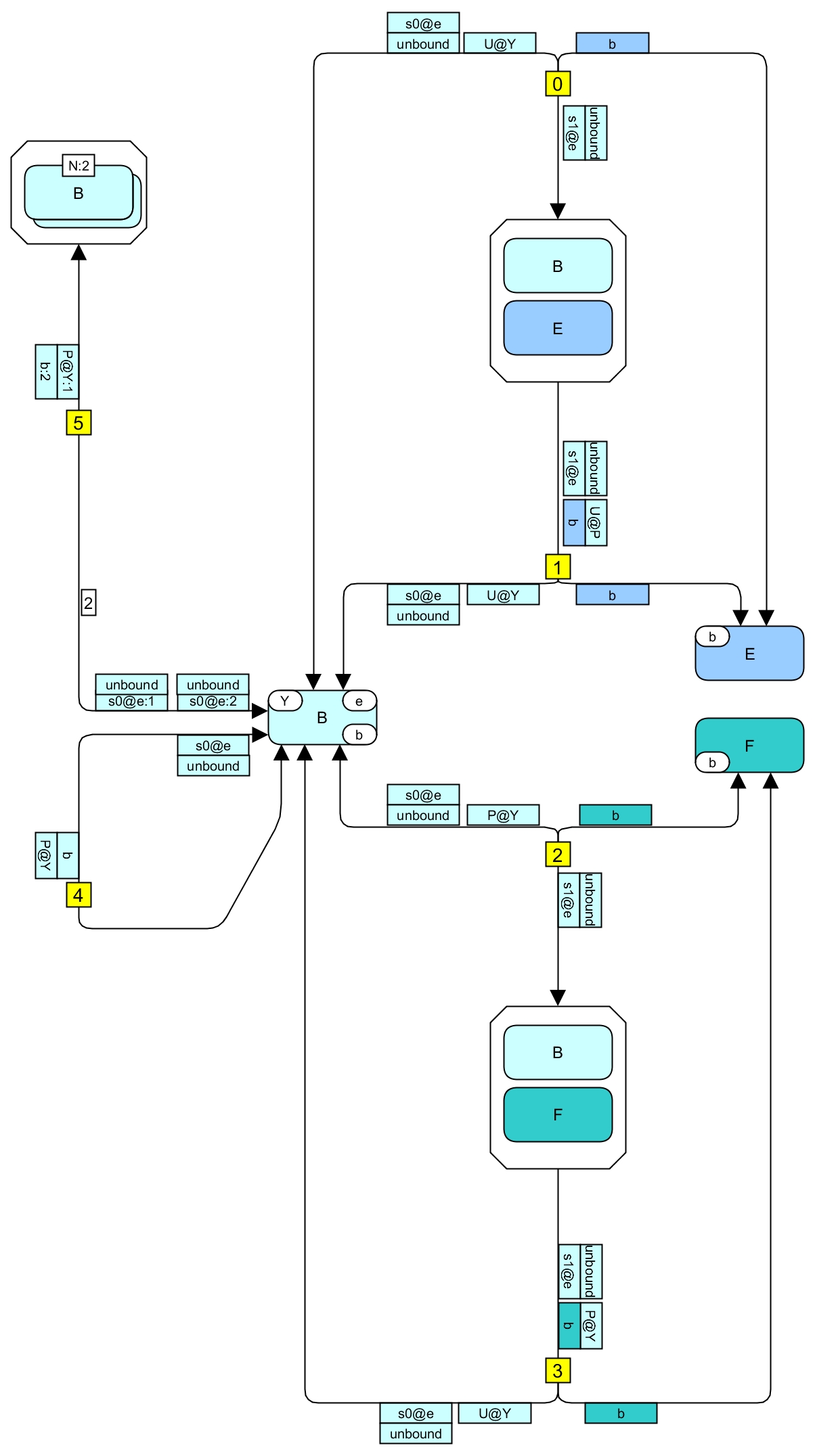

### Dushek_2014_rules.PNG

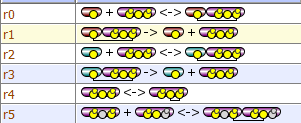

### Dushek_2014_VCell.PNG

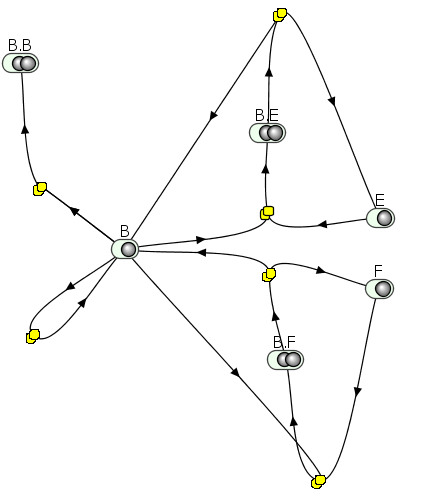

### Faeder_2003_L2.jpg

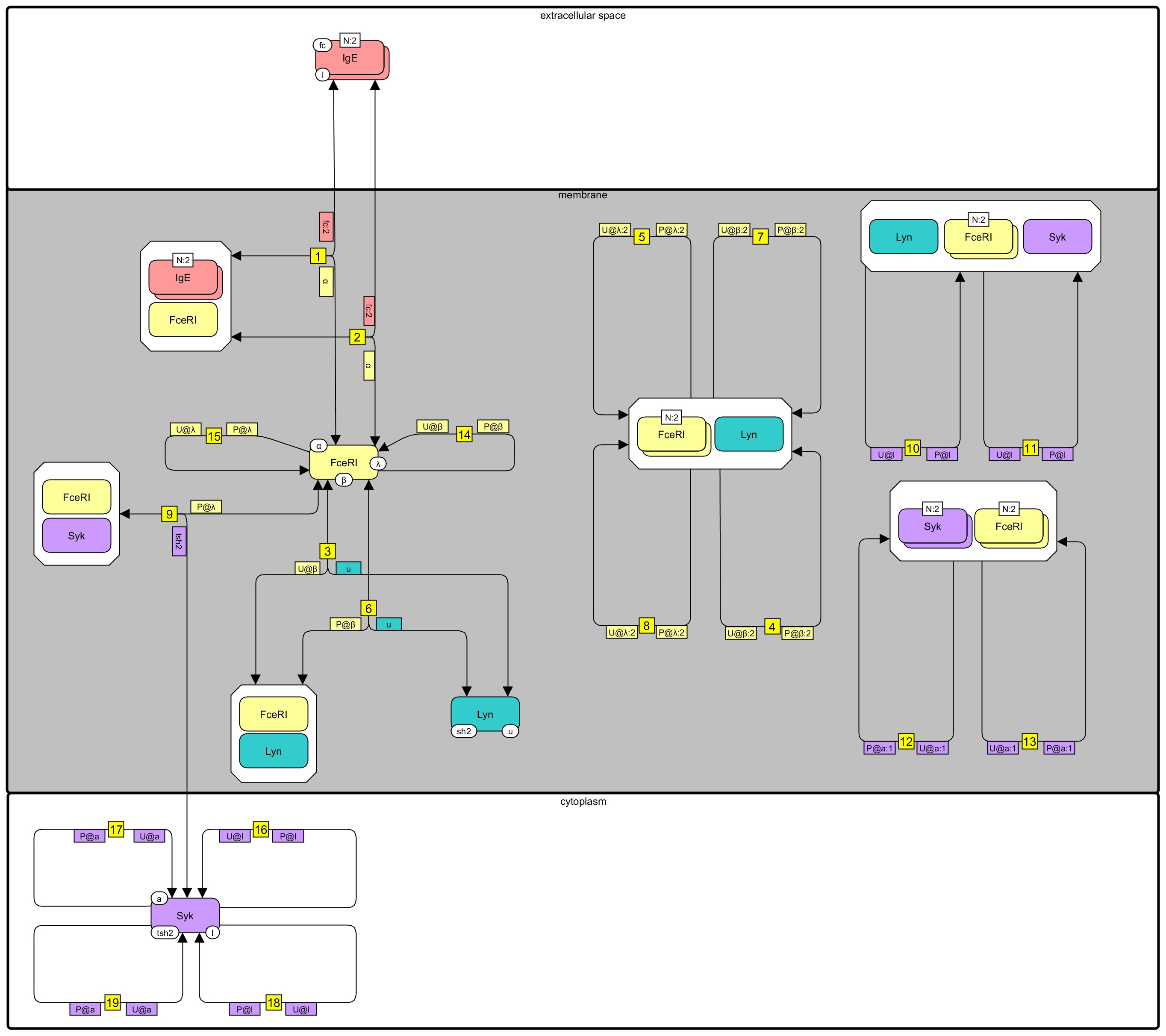

### Faeder_2003_L3.jpg

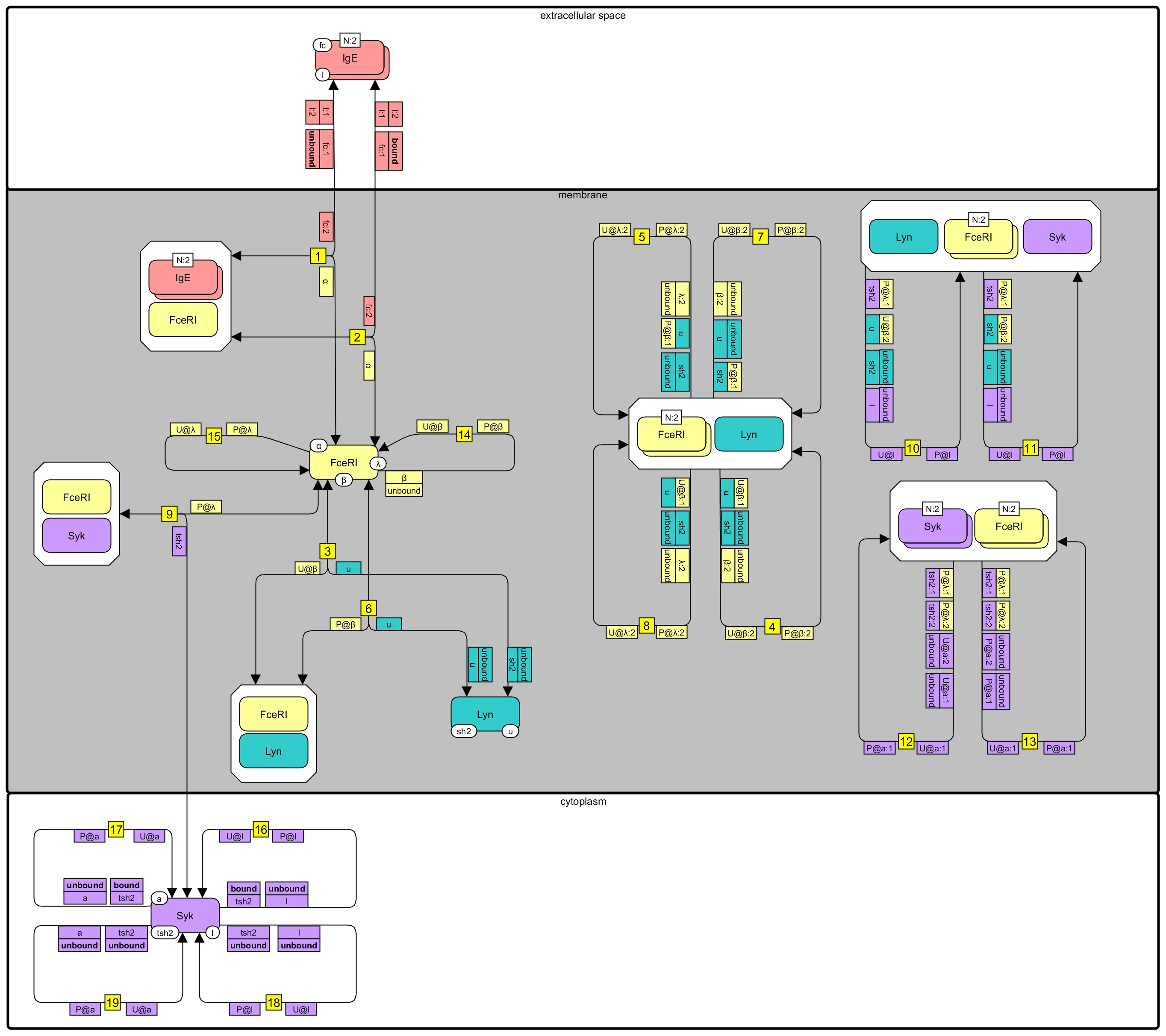

### Faeder_2003_rules.PNG

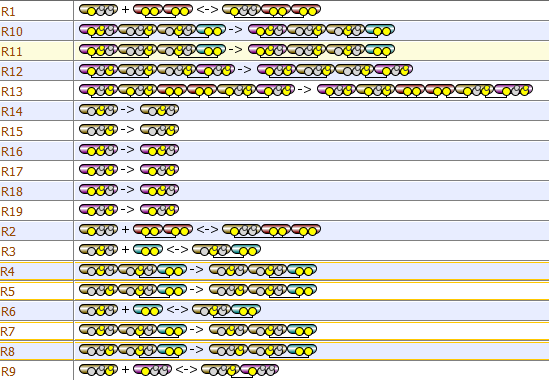

### Faeder_2003_VCell.PNG

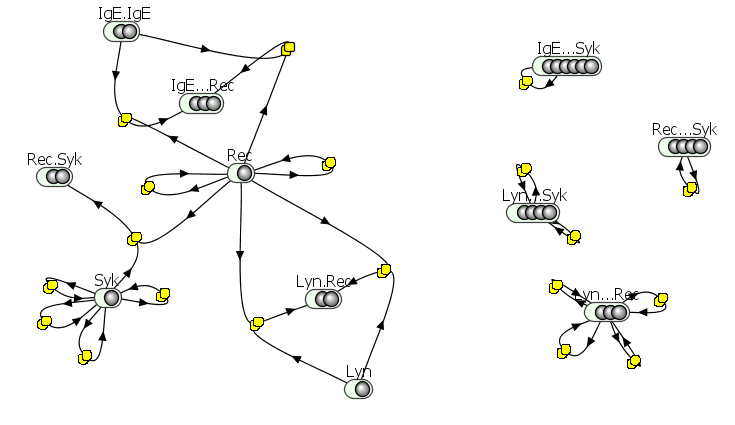

### Hat_2016_L2.jpg

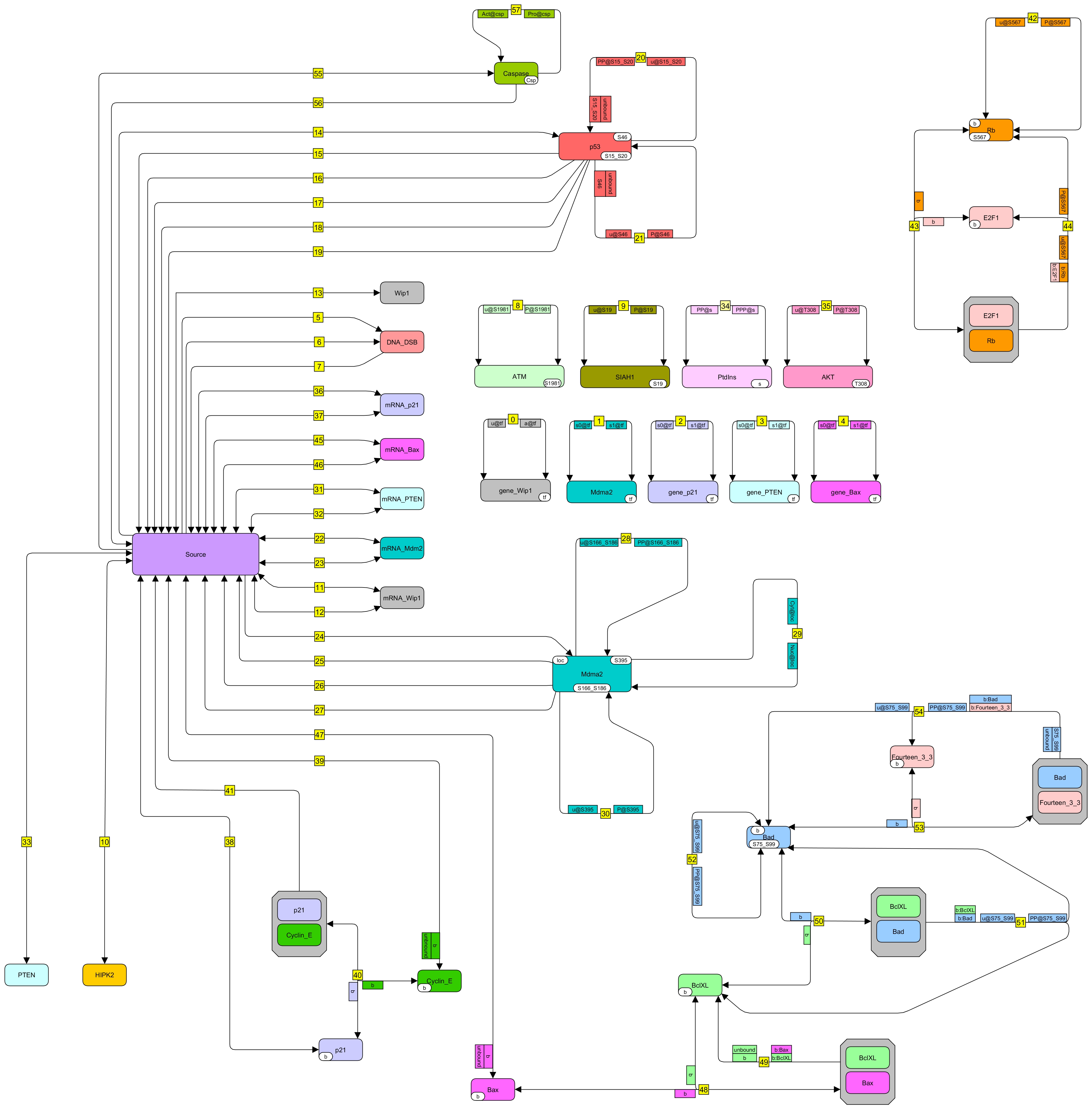

### Hat_2016_L3.jpg

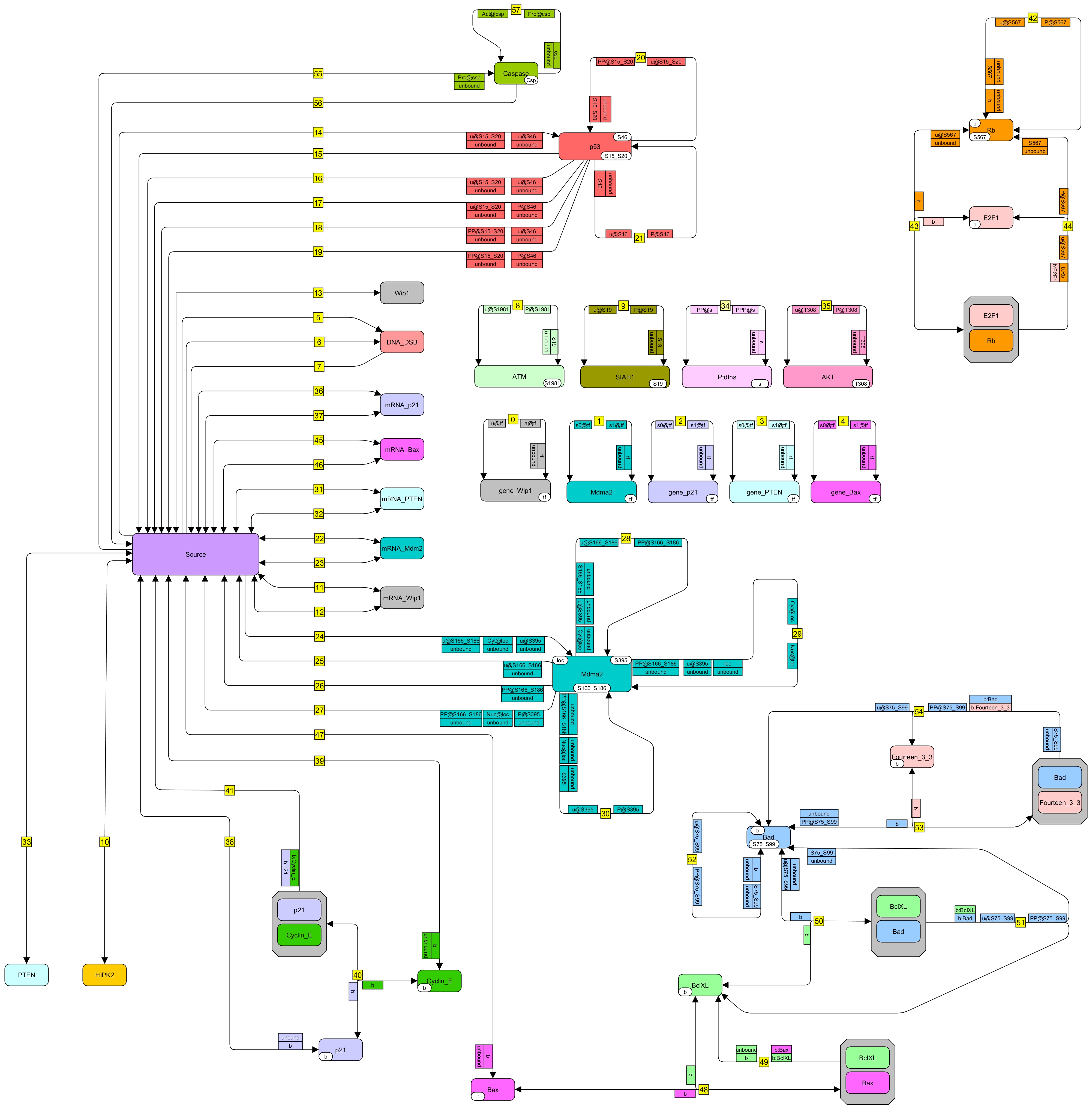

### Hat_2016_rules.PNG

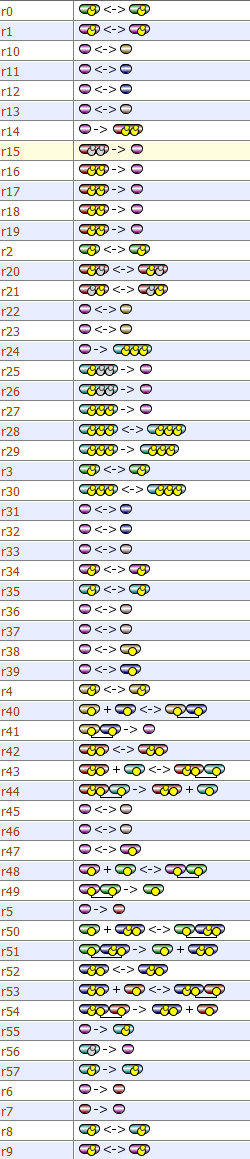

### Hat_2016_Vcell.PNG

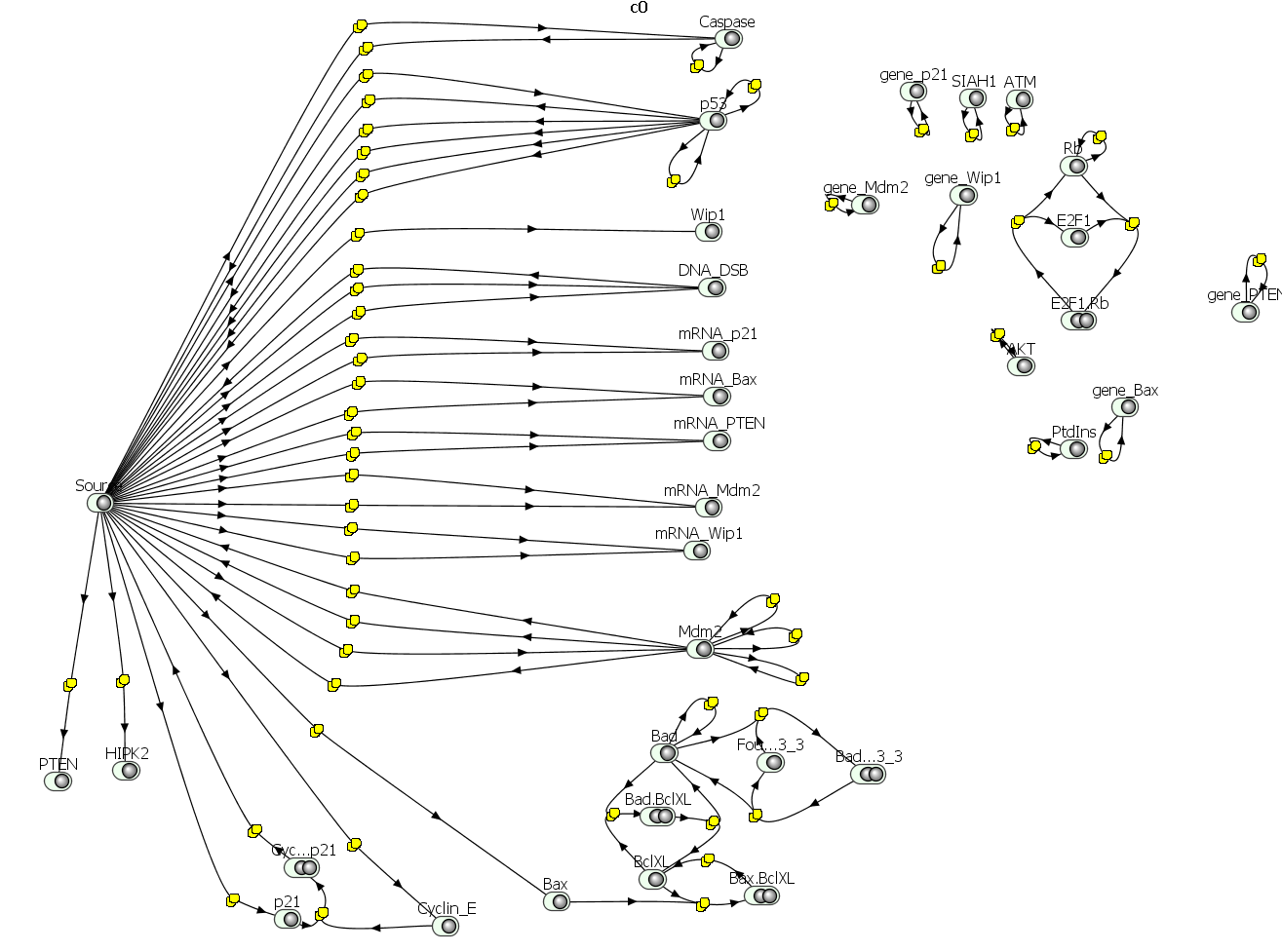

### Kocieniewski_2012_L2.jpg

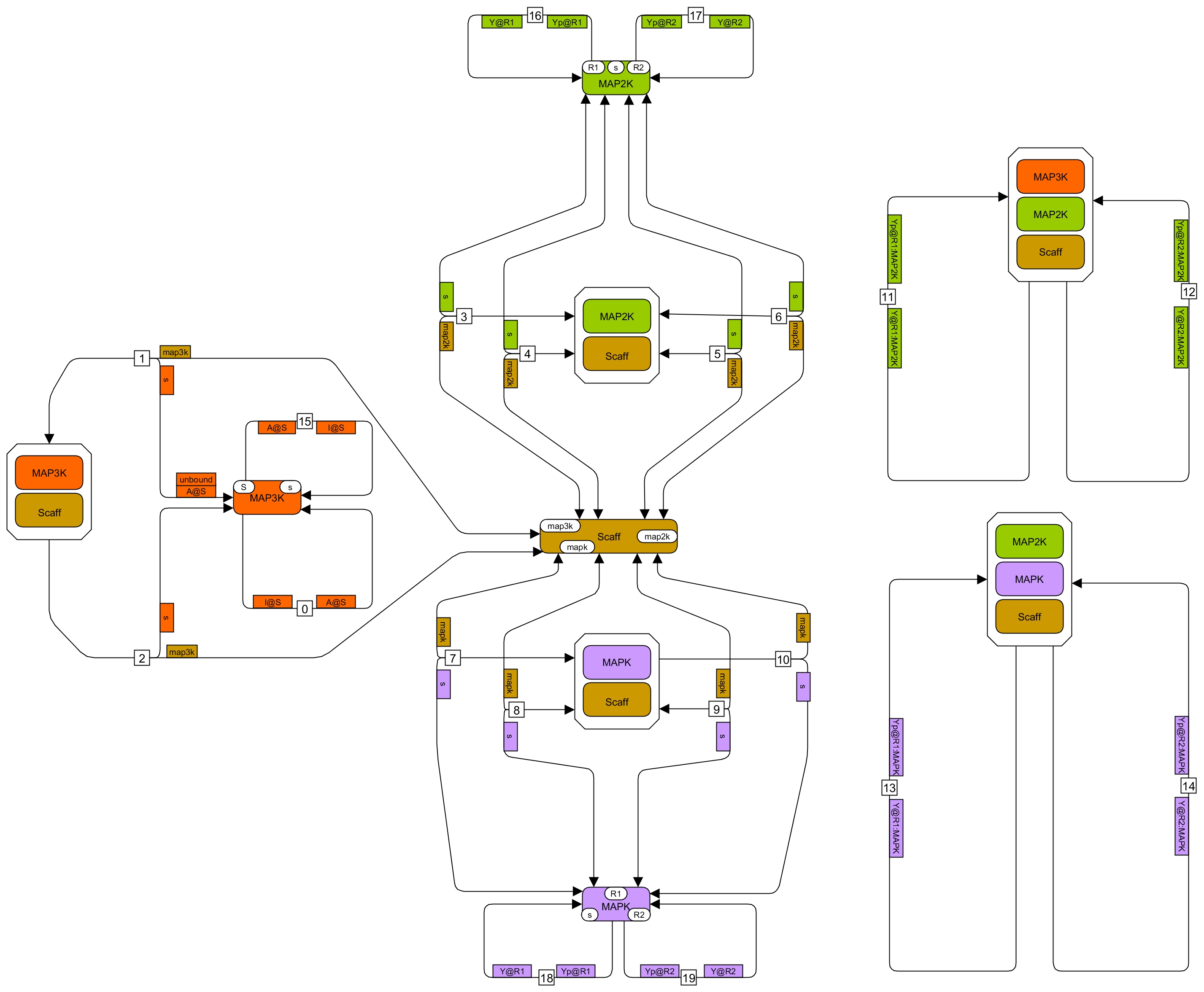

### Kocieniewski_2012_L3.jpg

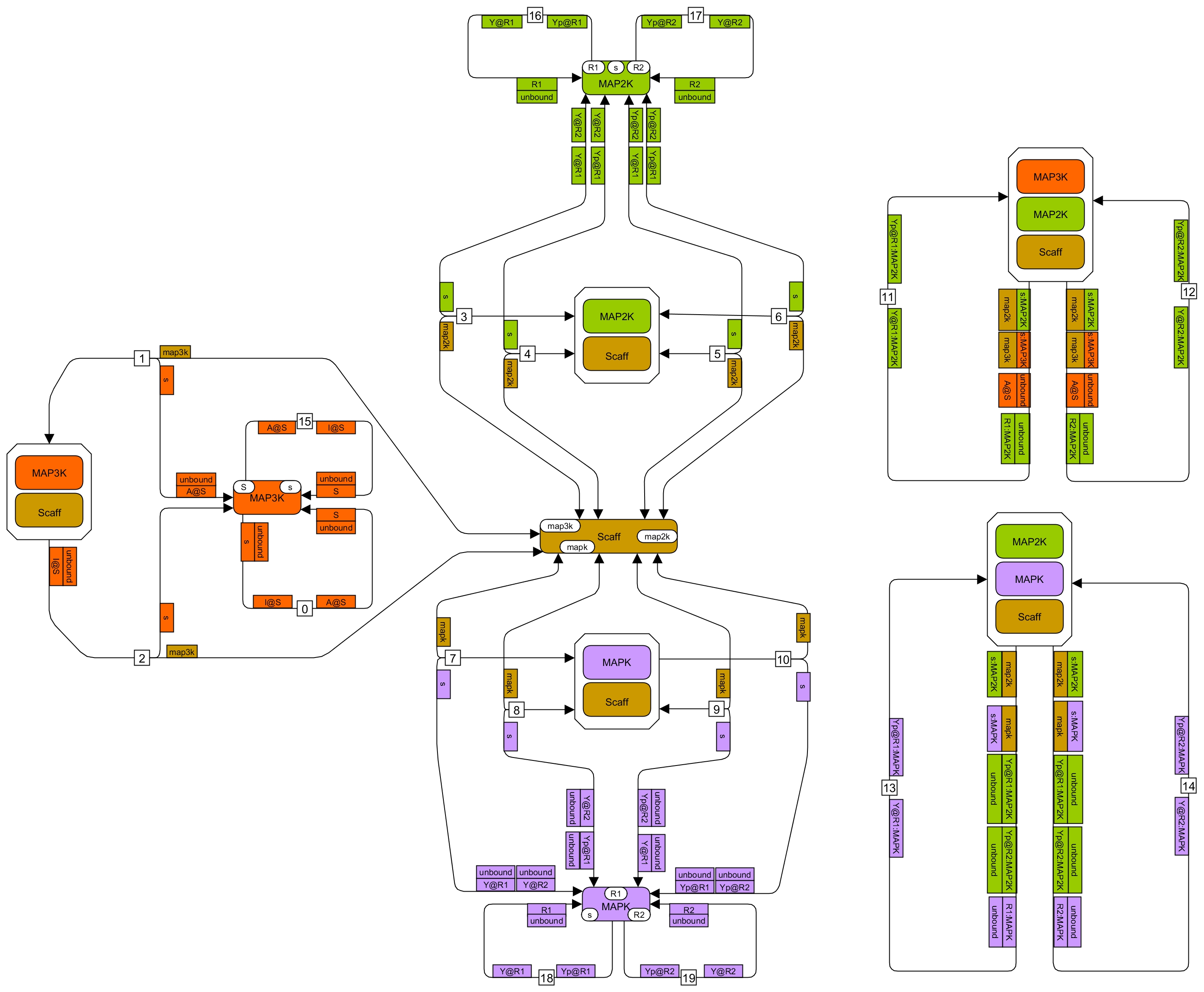

### Kocieniewski_2012_rules.jpg

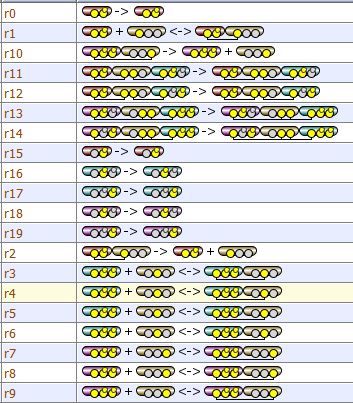

### Kocieniewski_2012_VCell.jpg

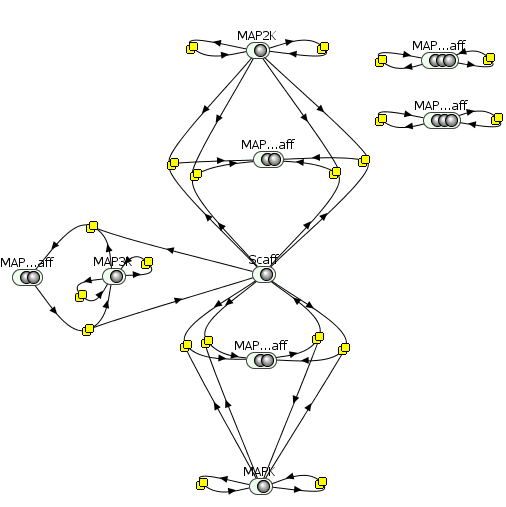

### Kozer_2014_L2.jpg

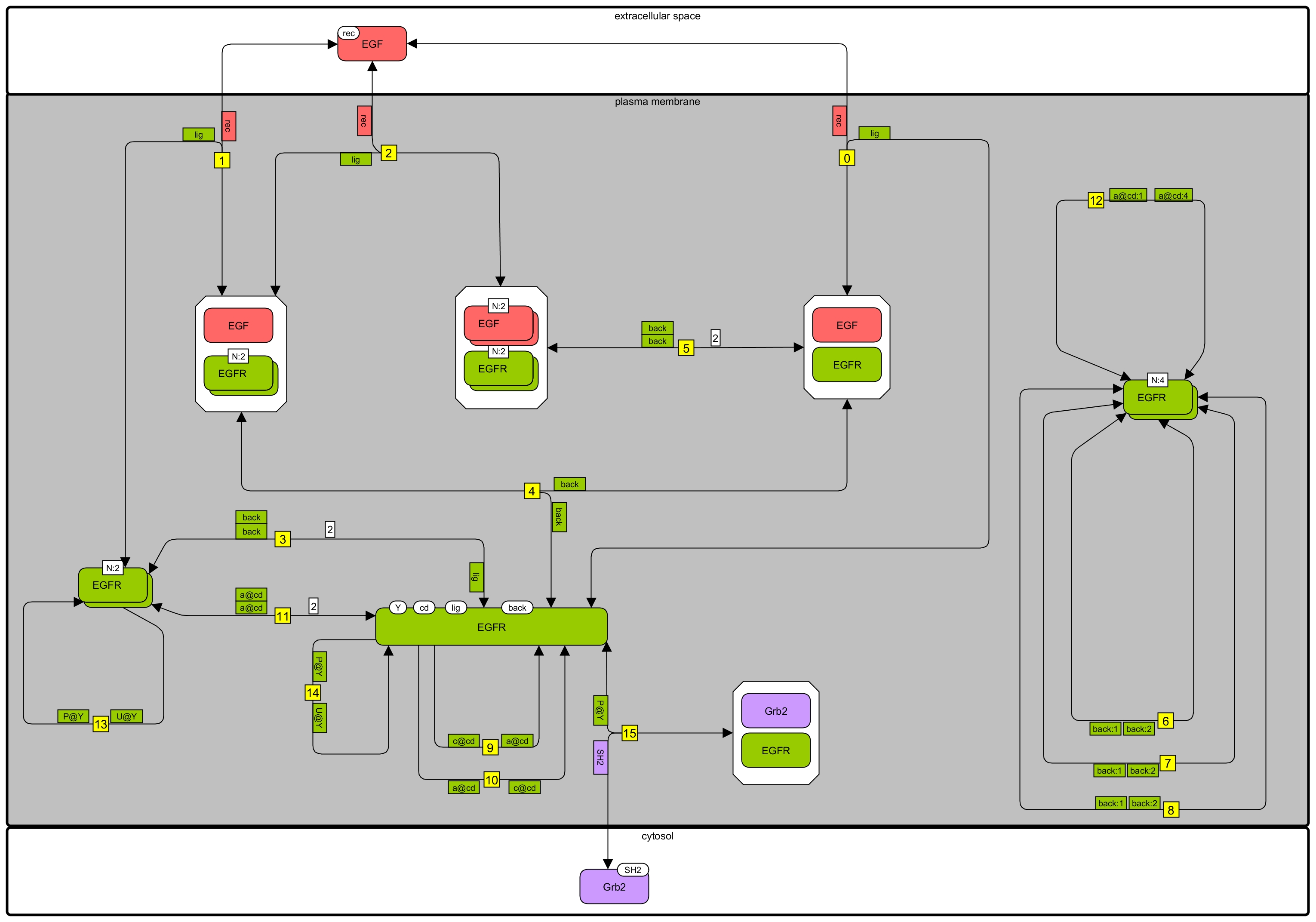

### Kozer_2014_L3.jpg

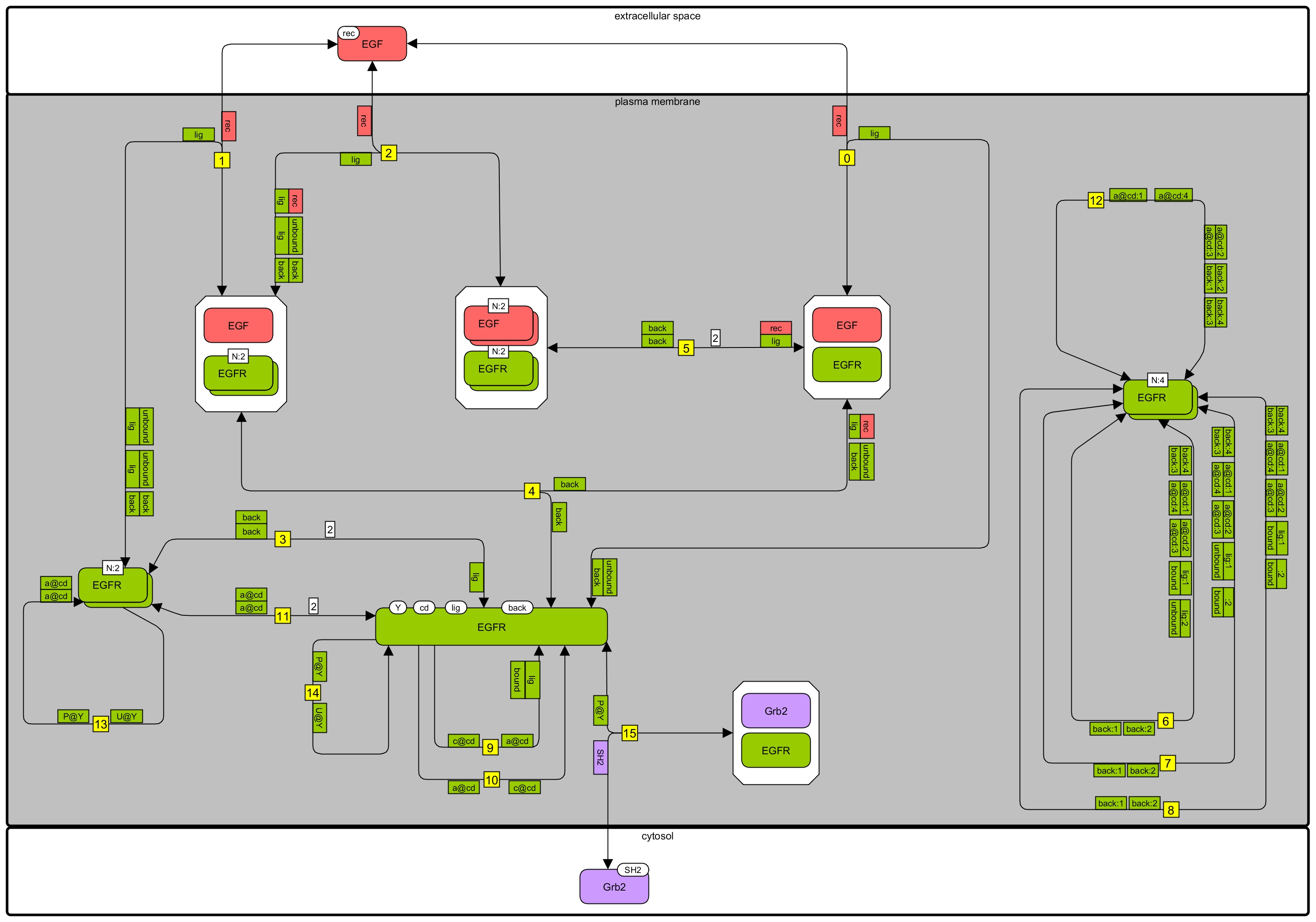

### Kozer_2014_rules.PNG

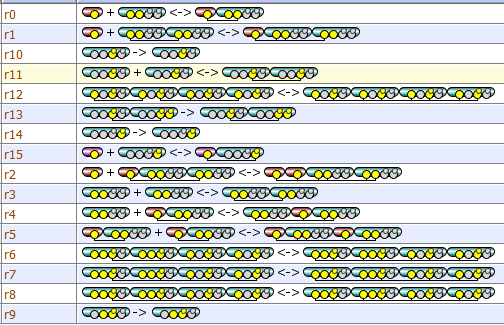

### Kozer_2014_VCell.PNG

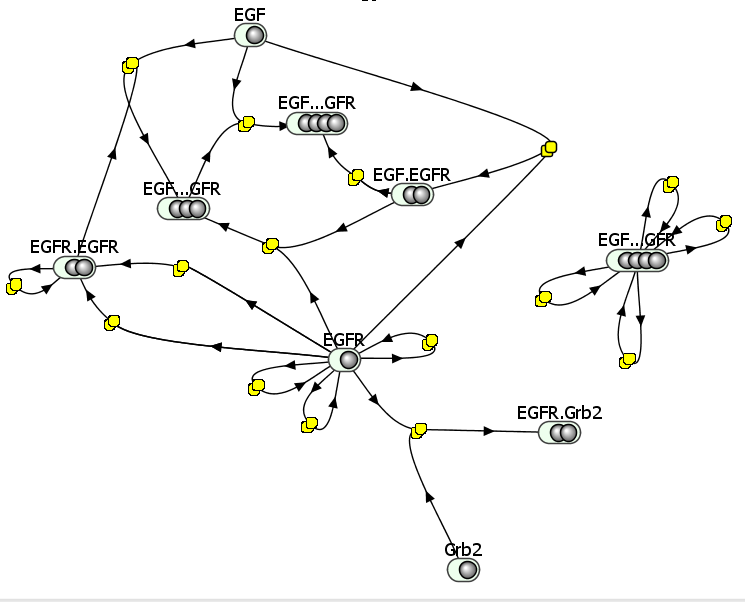

### Ligon_2014_L2.jpg

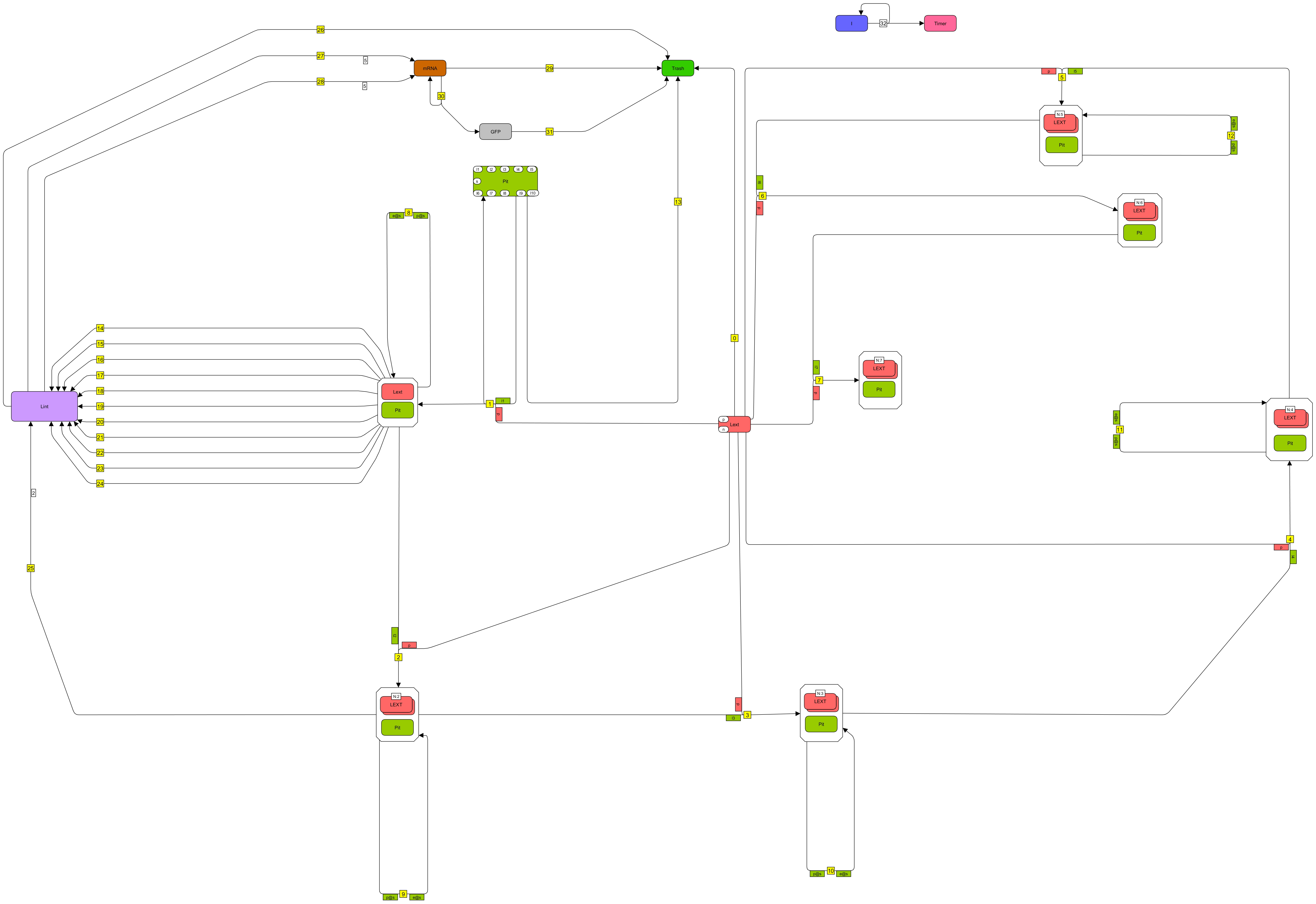

### Ligon_2014_L3.jpg

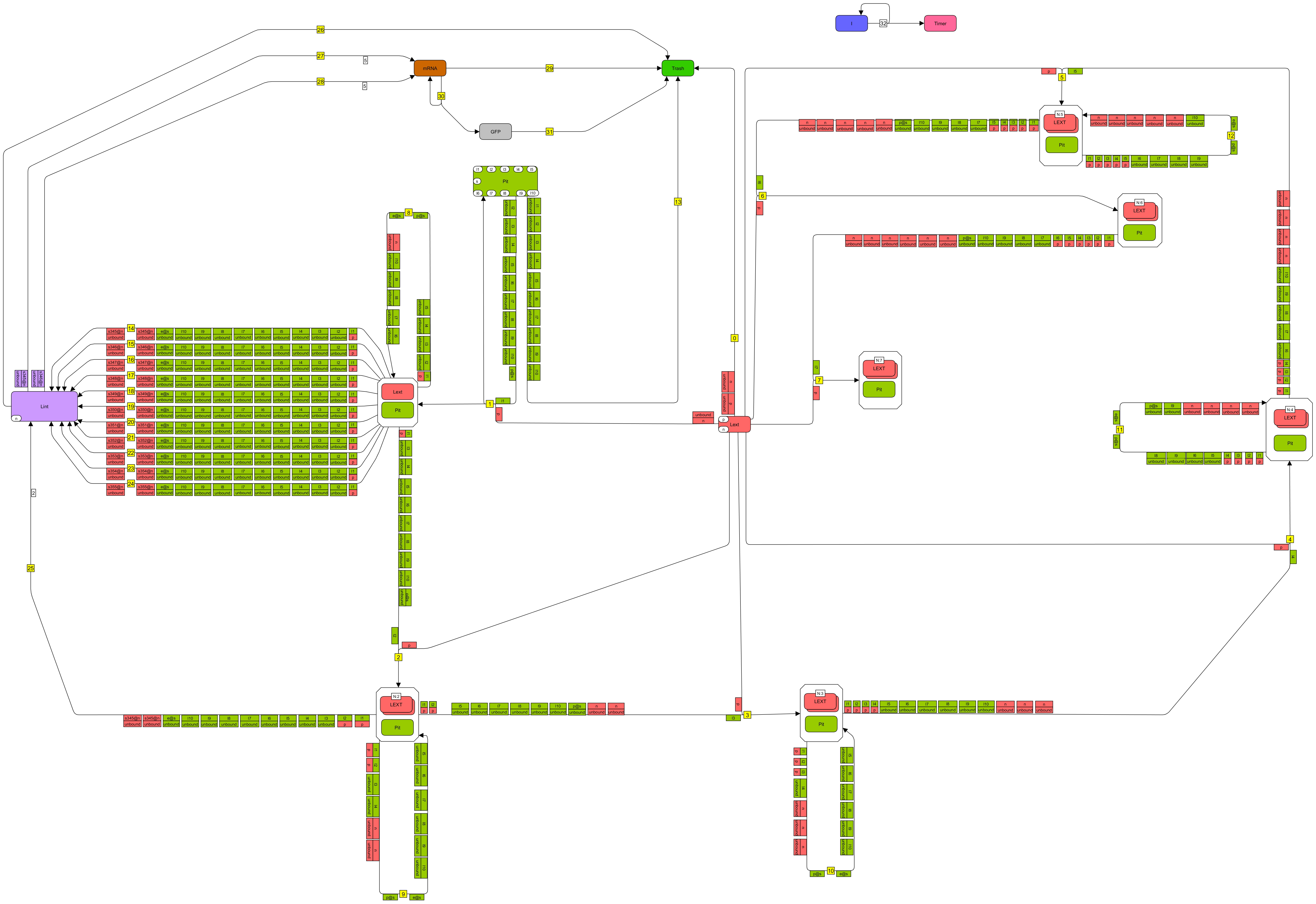

### Ligon_2014_rules.PNG

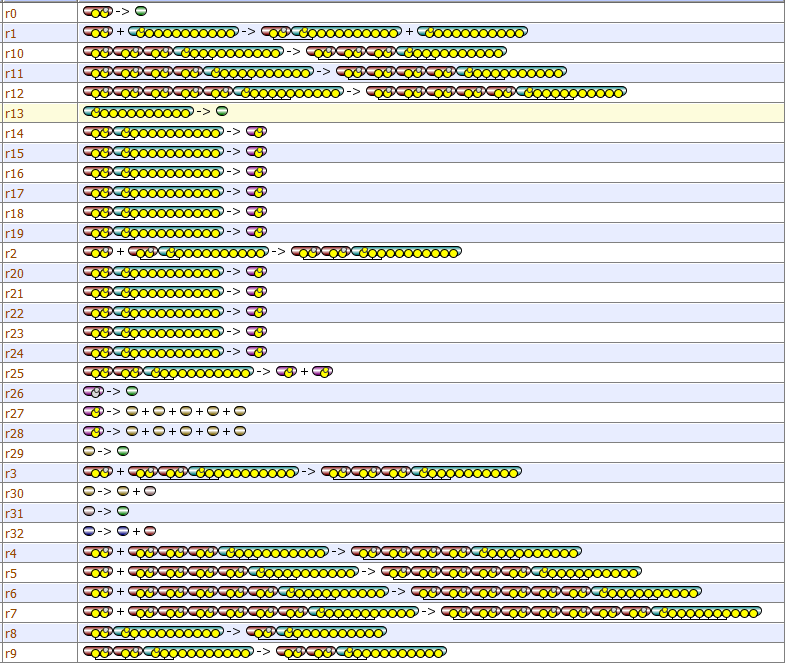

### Ligon_2014_VCell.PNG

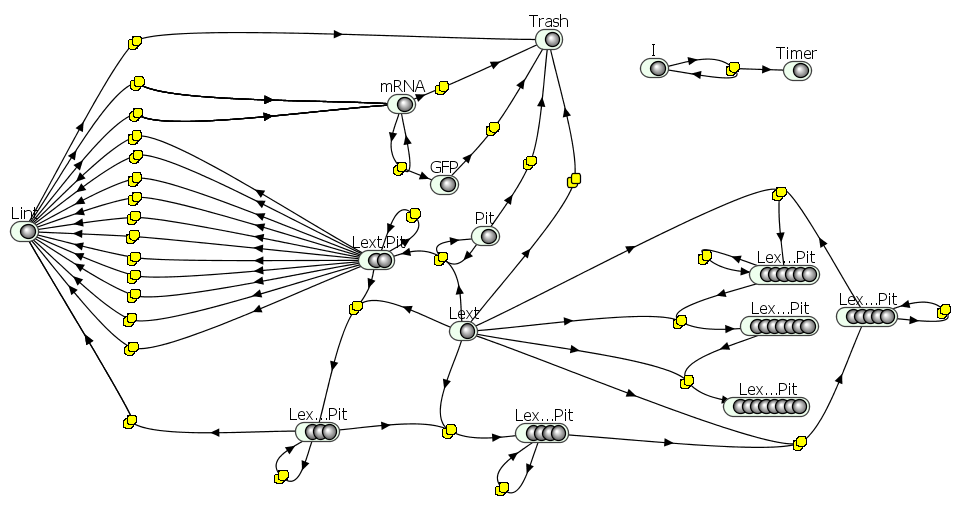

### Pekalski_2013_L2.jpg

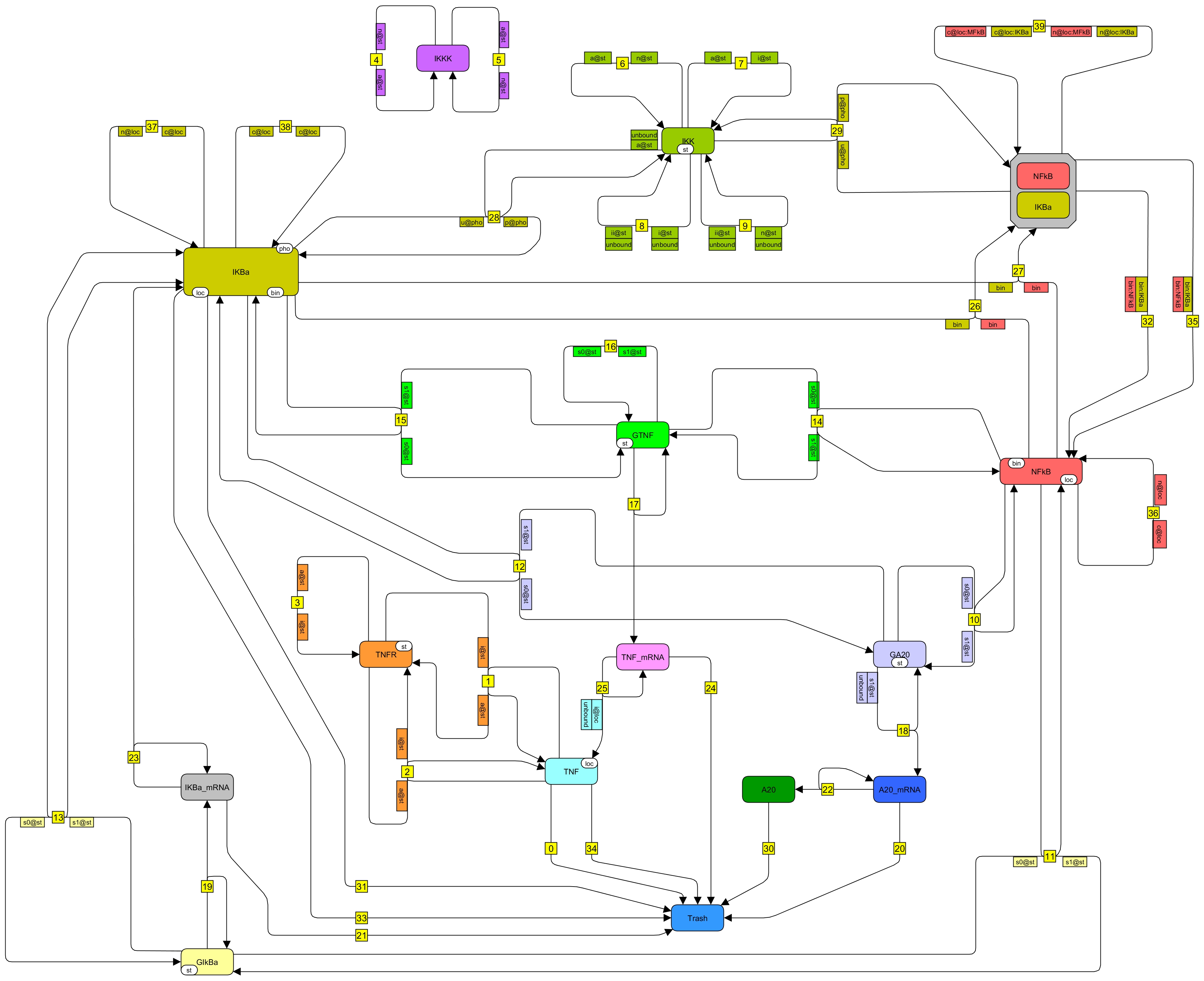

### Pekalski_2013_L3.jpg

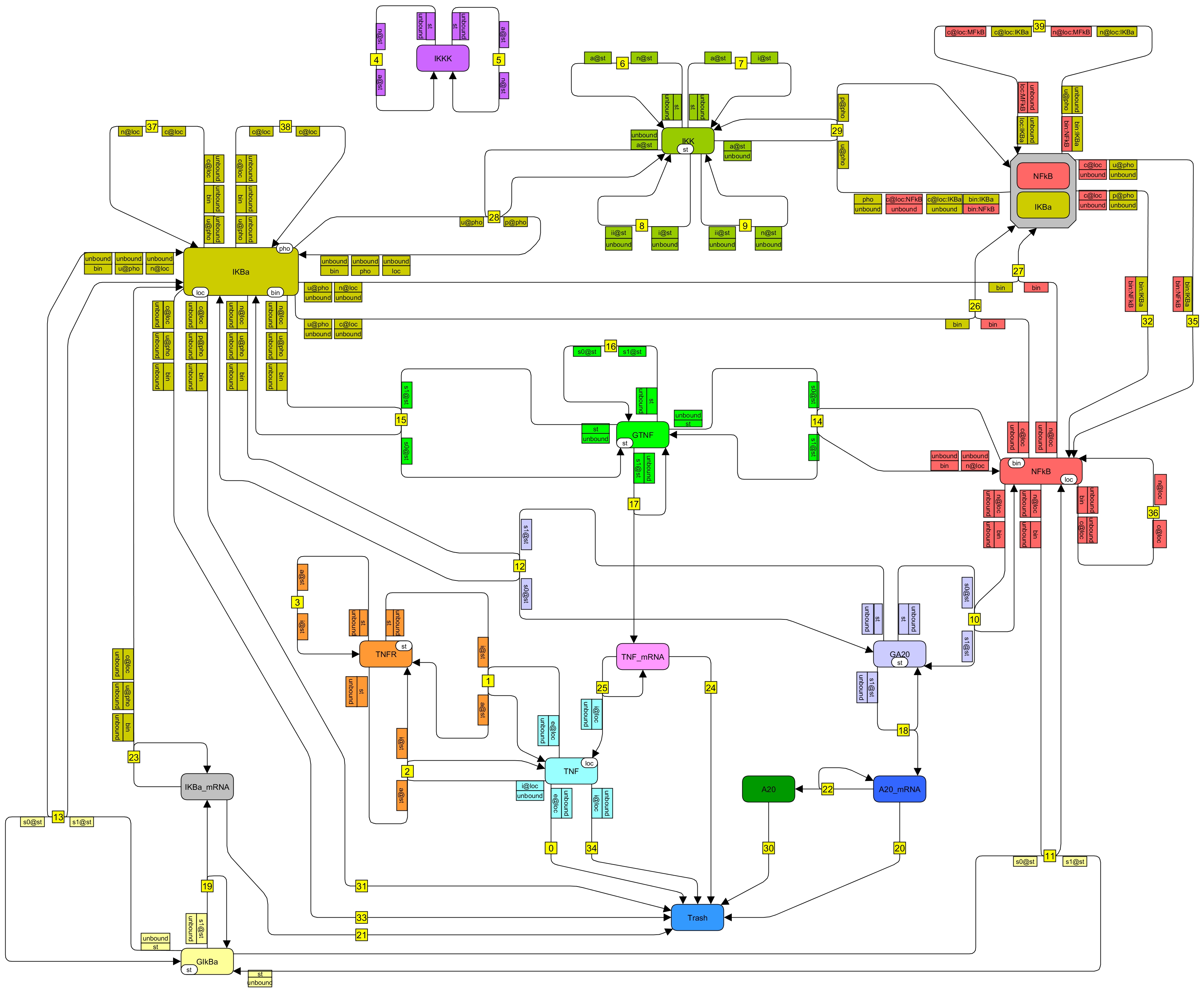

### Pekalski_2013_rules.PNG

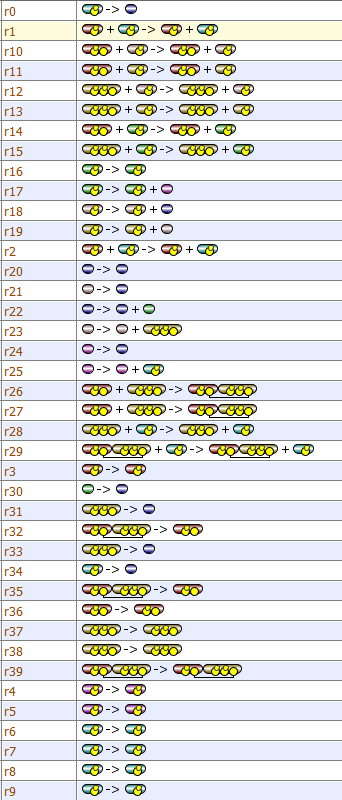

### Pekalski_2013_VCell.PNG

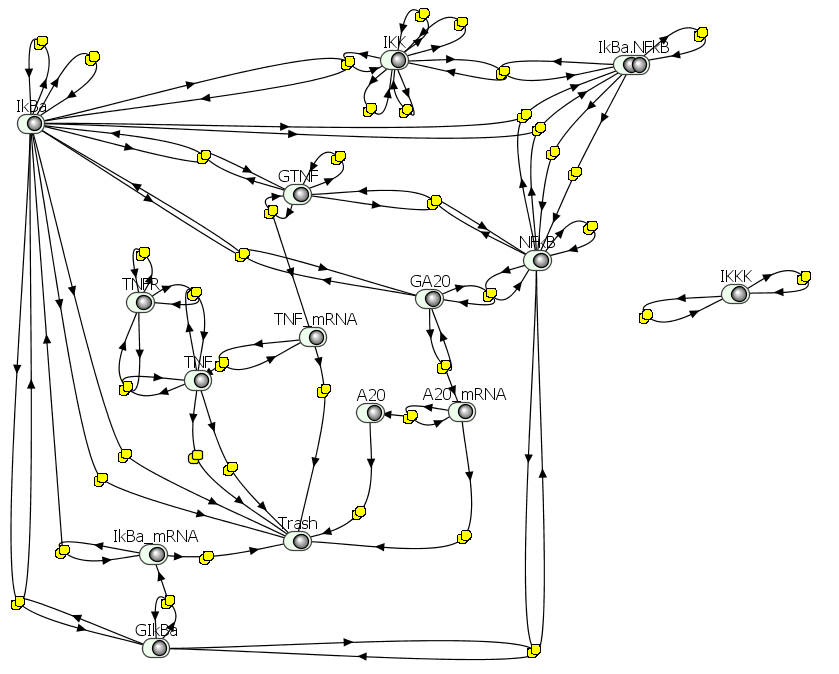

### Stites_2015_rules1.PNG

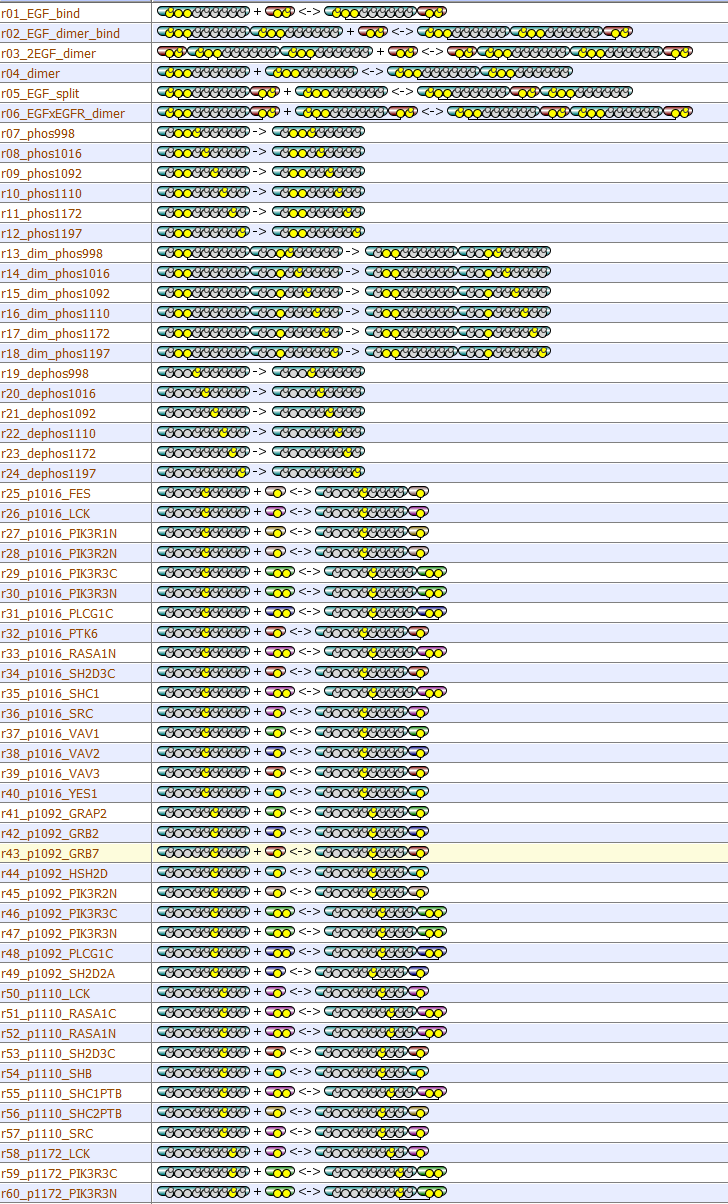

### Stites_2015_rules2.PNG

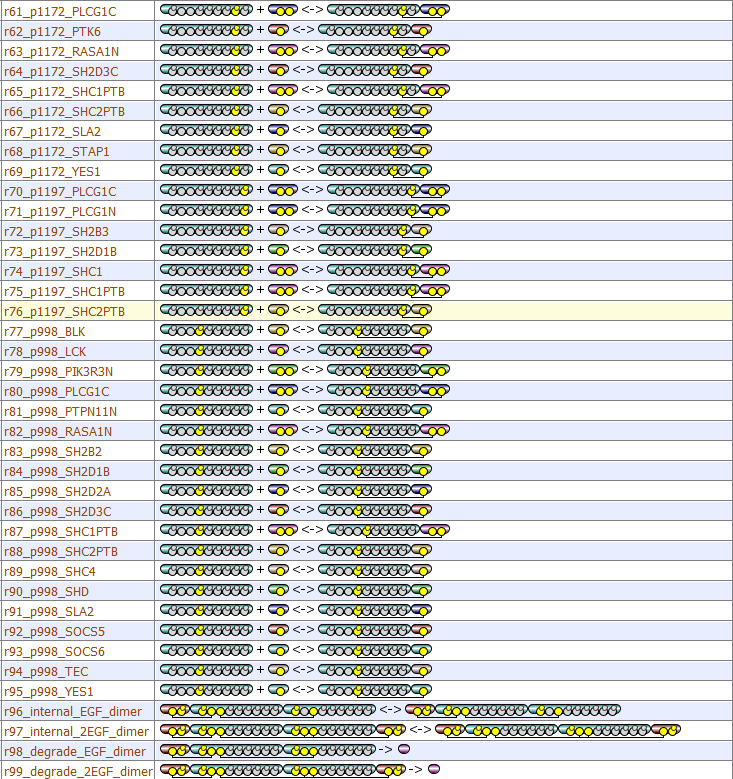

### Stites_2015_VCell.PNG

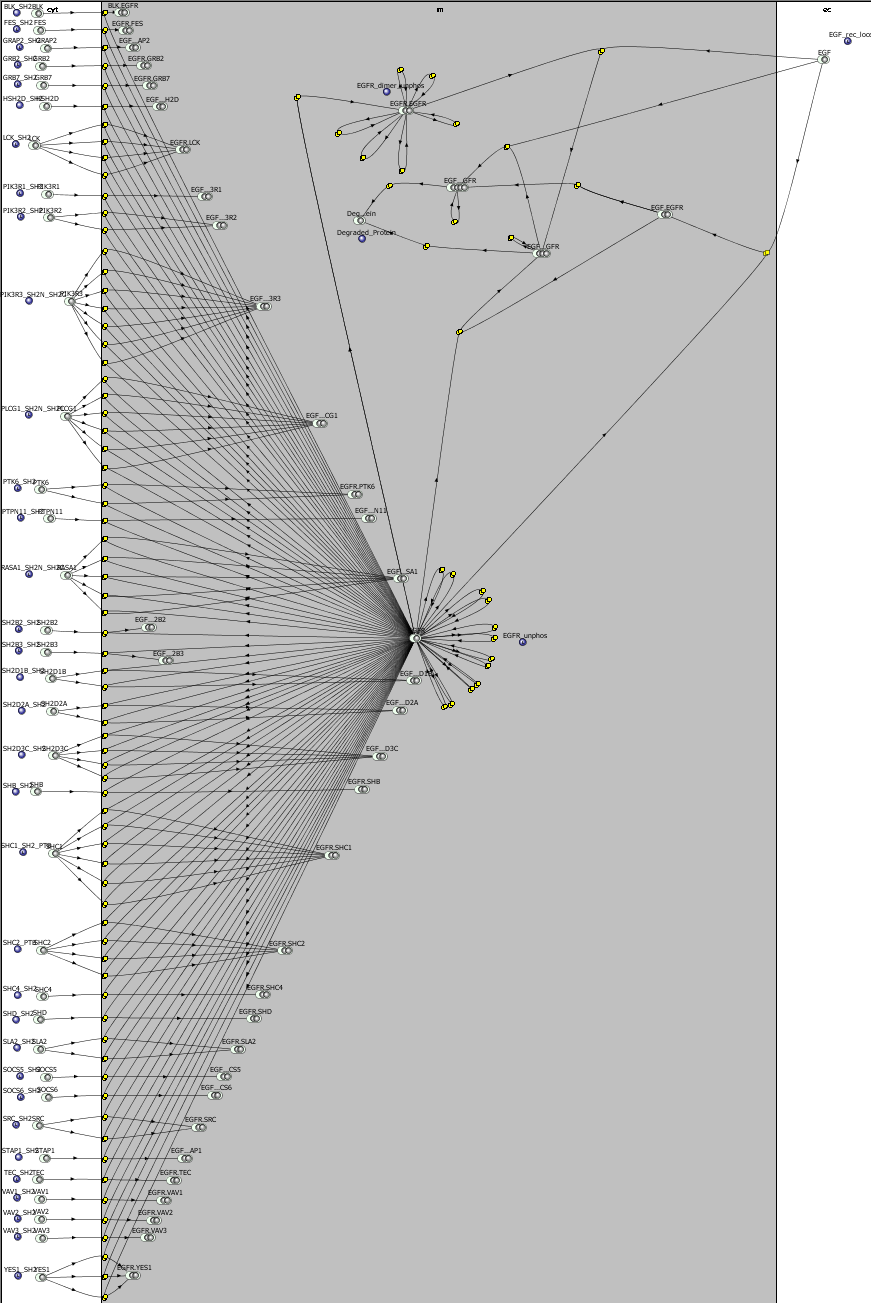

### Szymanska_2015_L2.jpg

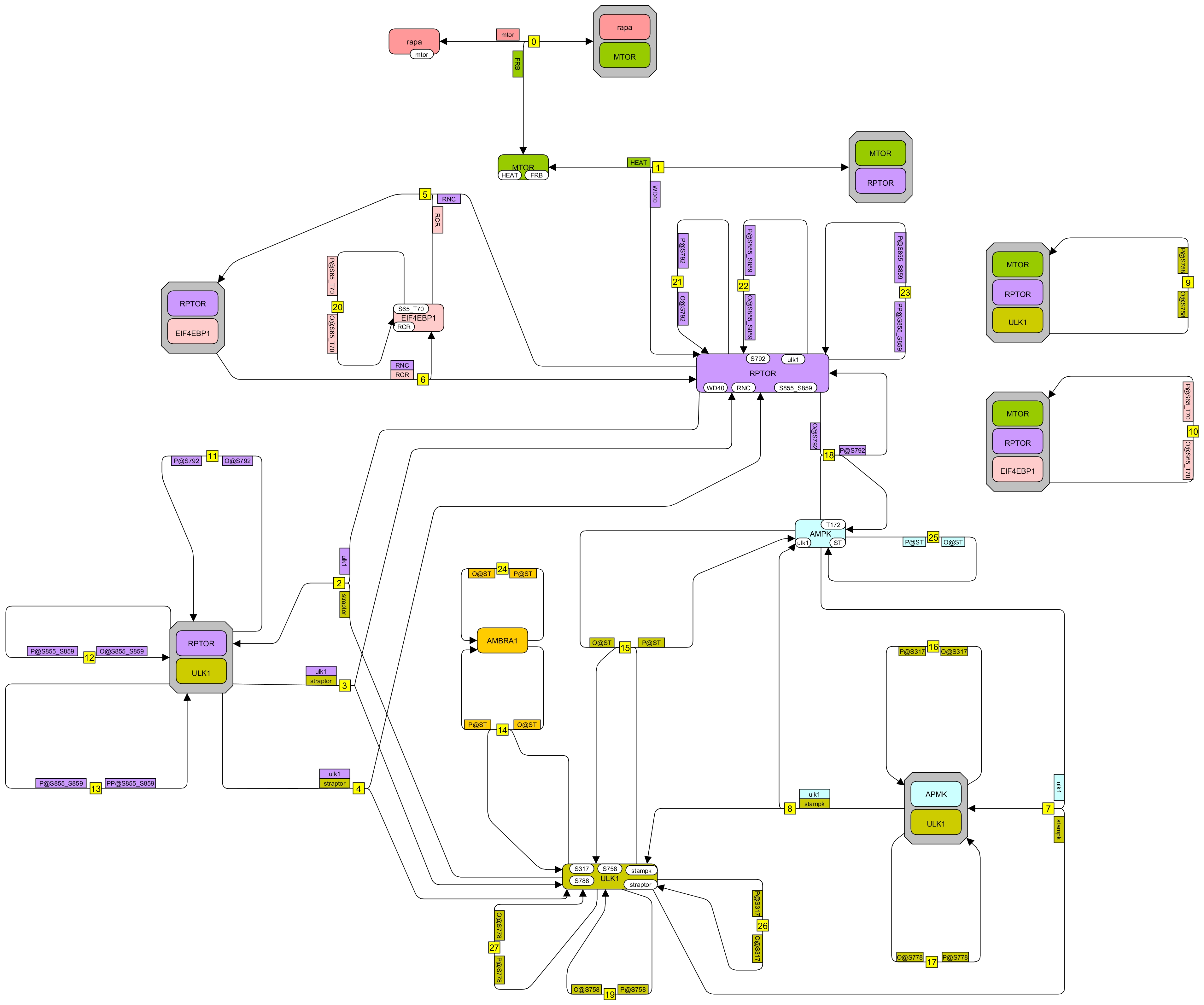

### Szymanska_2015_L3.jpg

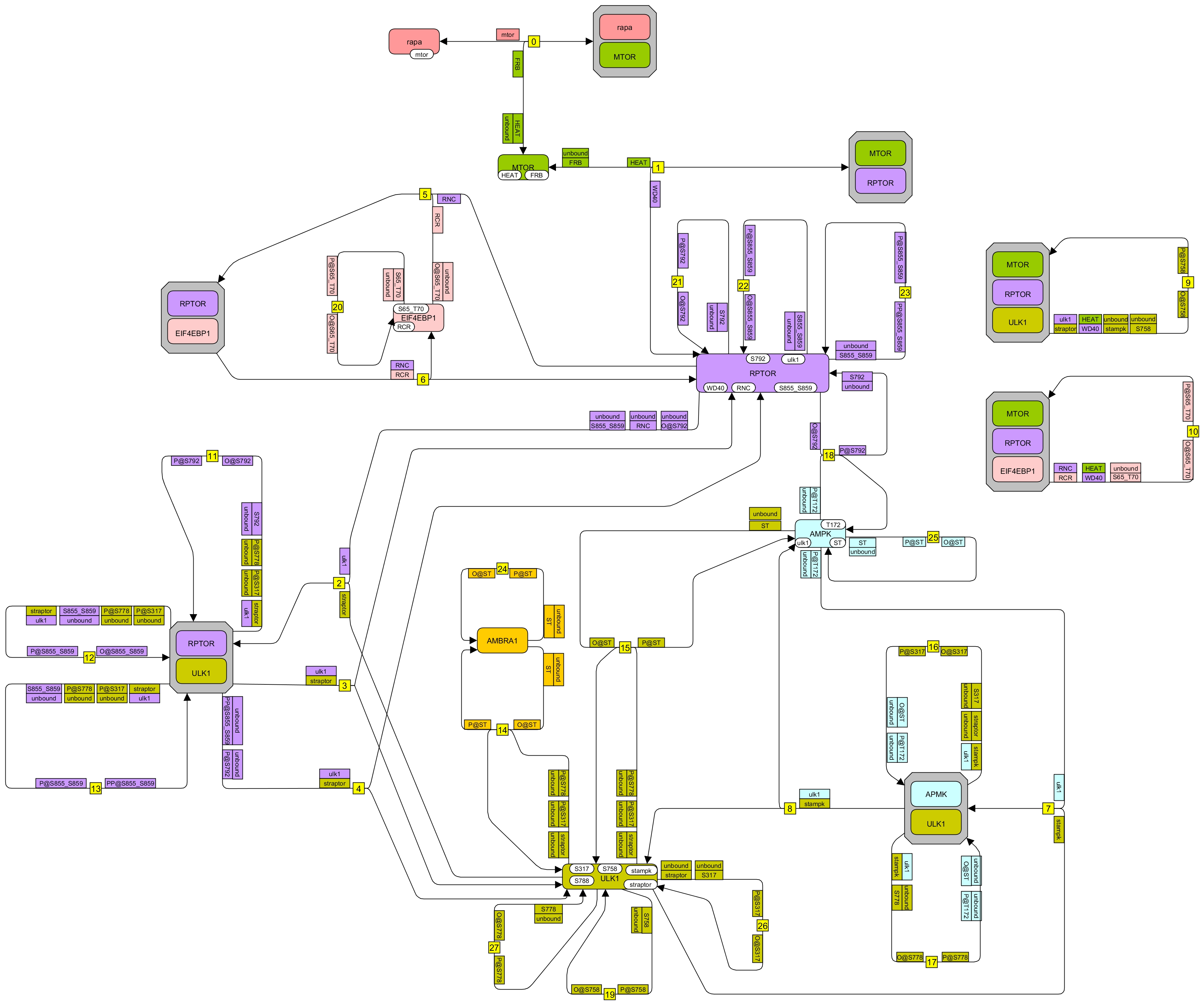

### Szymanska_2015_rules.PNG

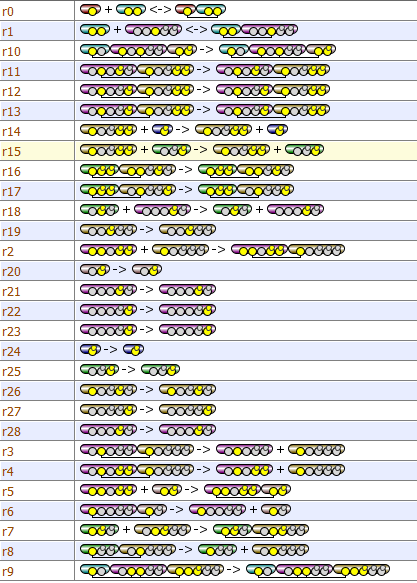

### Szymanska_2015_VCell.PNG

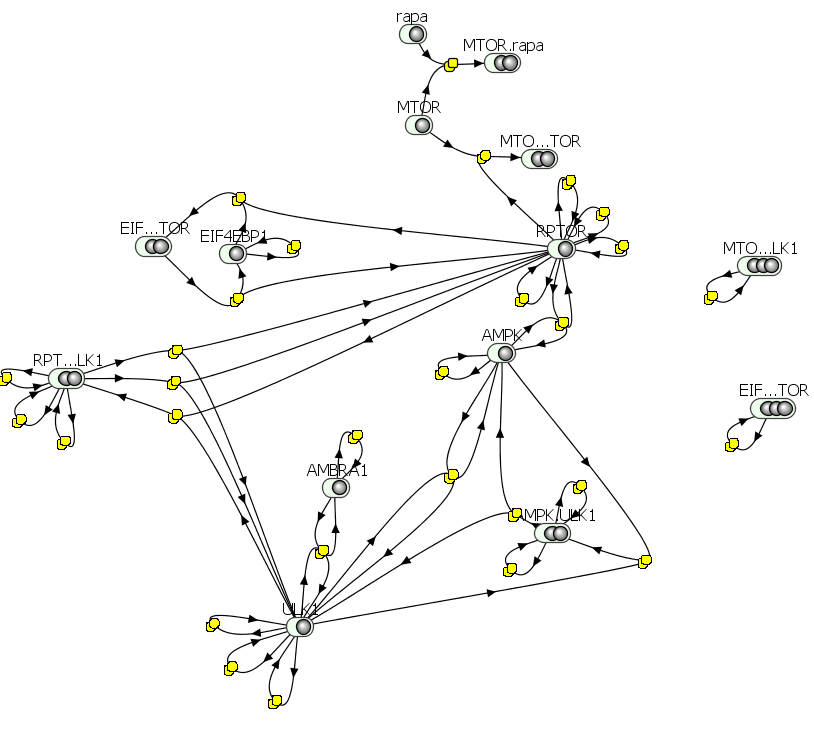
